# Elucidation and *de novo* Reconstitution of Glyceollin Biosynthesis

**DOI:** 10.1101/2025.02.07.636353

**Authors:** Yunlong Sun, Cong Chen, Chao Lin, Hao Zhang, Jiazhang Lian, Benke Hong

## Abstract

Glyceollins are phytoalexins produced by soybeans in response to stressors such as pathogen invasion, injury, and environmental challenges. In addition to their antibacterial and antifungal activities, these compounds have attracted significant attention for their potential anticancer, anti-inflammatory, and antioxidant properties. However, their limited accessibility—due to the challenges and high costs associated with organic synthesis or purification from treated soybean seedlings—has hindered further physiological and biochemical studies. Moreover, the incomplete understanding of glyceollin biosynthesis, particularly the final cyclization steps, remains a major barrier to elucidating the physiological functions of glyceollins and achieving sustainable production through synthetic biology. In this study, we identified previously uncharacterized genes encoding two reductases for 7,2’,4’-trihydroxyisoflavanol (THI) biosynthesis and five P450 enzymes responsible for the final oxidative cyclization in glyceollin I, II, and III biosynthesis, thereby completing the entire glyceollin biosynthetic pathway. By reconstructing the pathway *de novo* through synthetic biology, we achieved the successful production of glyceollins from simple carbon sources in baker’s yeast. This work advances the understanding of glyceollin biosynthesis in soybean, facilitates sustainable production in microbial hosts and offers new opportunities for their application in agriculture and biology.

## Introduction

Soybean (*Glycine max*), one of the most important crop plants in the world for dietary protein and oil production, often suffers yield loss due to attacks by pathogens such as *Phytophthora sojae* (Whitham et al., 2016; Prayogo et al., 2021; Rao et al., 2023). In response to these threats, soybean plants leverage their natural resistance mechanisms to produce isoflavonoid phytoalexins such as glyceollins (Burden and Bailey, 1975; Robert Lyne et al., 1976). In addition to their antibacterial and antifungal activities (Kim et al., 2010; Araya-Cloutier et al., 2018), glyceollins have garnered significant attention because of their anticancer, anti-inflammatory, and antioxidative properties (Bamji and Corbitt, 2017; Pham et al., 2019; Yue et al., 2023), with particular potential for targeting breast cancer through the oestrogen receptor signalling pathway (Salvo et al., 2006; Zimmermann et al., 2010; Payton-Stewart et al., 2010; Lecomte et al., 2017). Unlike the ubiquitous isoflavonoids daidzein and genistein, which are constitutively present in soybeans (Medic et al., 2014), glyceollins are synthesized only under specific environmental triggers, such as pathogen infection or exposure to other elicitors. Soybean predominantly produces three glyceollin isomers, namely, glyceollin I, glyceollin II, and glyceollin III. Pure forms of glyceollins can be purified from soybeans, but the methods for large-scale isolation are time-consuming and unsuitable for mass production, which has driven interest in alternative production methods (Luniwal et al., 2011; Ciesielski and Metz, 2020). Although the chemical synthesis of glyceollins I and II has been reported (Khupse and Erhardt, 2008; Khupse et al., 2011; Luniwal et al., 2011; Kohno et al., 2014; Malik et al., 2015; Ciesielski and Metz, 2020), the complex synthetic routes and high manufacturing costs have limited their practical applications for biological use.

Since the discovery of glyceollin I in soybean in 1972 (Sims et al., 1972), the biosynthetic mechanism of glyceollins has been extensively investigated (**Figure 1**) (Yue et al., 2023). These compounds are synthesized from phenylalanine (**1**), which is converted into the isoflavone daidzein *via* eight known genes. Daidzein (**2**) is transformed into 2’-hydroxydaidzein (**3**) by CYP81E (Uchida et al., 2015), followed by reduction to 2’-OH dihydrodaidzein (**4**) by isoflavone reductase (IFR) (Fischer et al., 1990b; Uchida et al., 2017). A subsequent reduction, mediated by an uncharacterized reductase, produces 7,2’,4’-trihydroxyisoflavanol (THI, **5**), which is converted to 3,9-OH pterocarpan (**6**) by pterocarpan synthase (PTS) (Uchida et al., 2017) and further modified to glycinol (**7**) by CYP93A1 (Schopfer et al., 1998). Glycinol (**7**) undergoes dimethyl allylation at the C2 or C4 position, which is catalysed by the prenyltransferase G2DT (Yoneyama et al., 2016) or G4DT (Akashi et al., 2009; Sukumaran et al., 2018), resulting in the formation of 4-glyceollidin (**8**) or glyceocarpin (**9**), respectively. However, the final cyclization steps remain unresolved, as the genes encoding the responsible enzymes have not been identified. It is unclear whether these steps are catalysed by a single gene or multiple paralogous genes. Studies with soybean cell culture microsomes have suggested the involvement of cytochrome P450 (CYP) enzyme(s) capable of converting 4-glyceollidin (**8**) and glyceocarpin (**9**) into glyceollins I (**10**), II (**11**), and III (**12**) (Welle and Grisebach, 1988).

**Figure 1:**
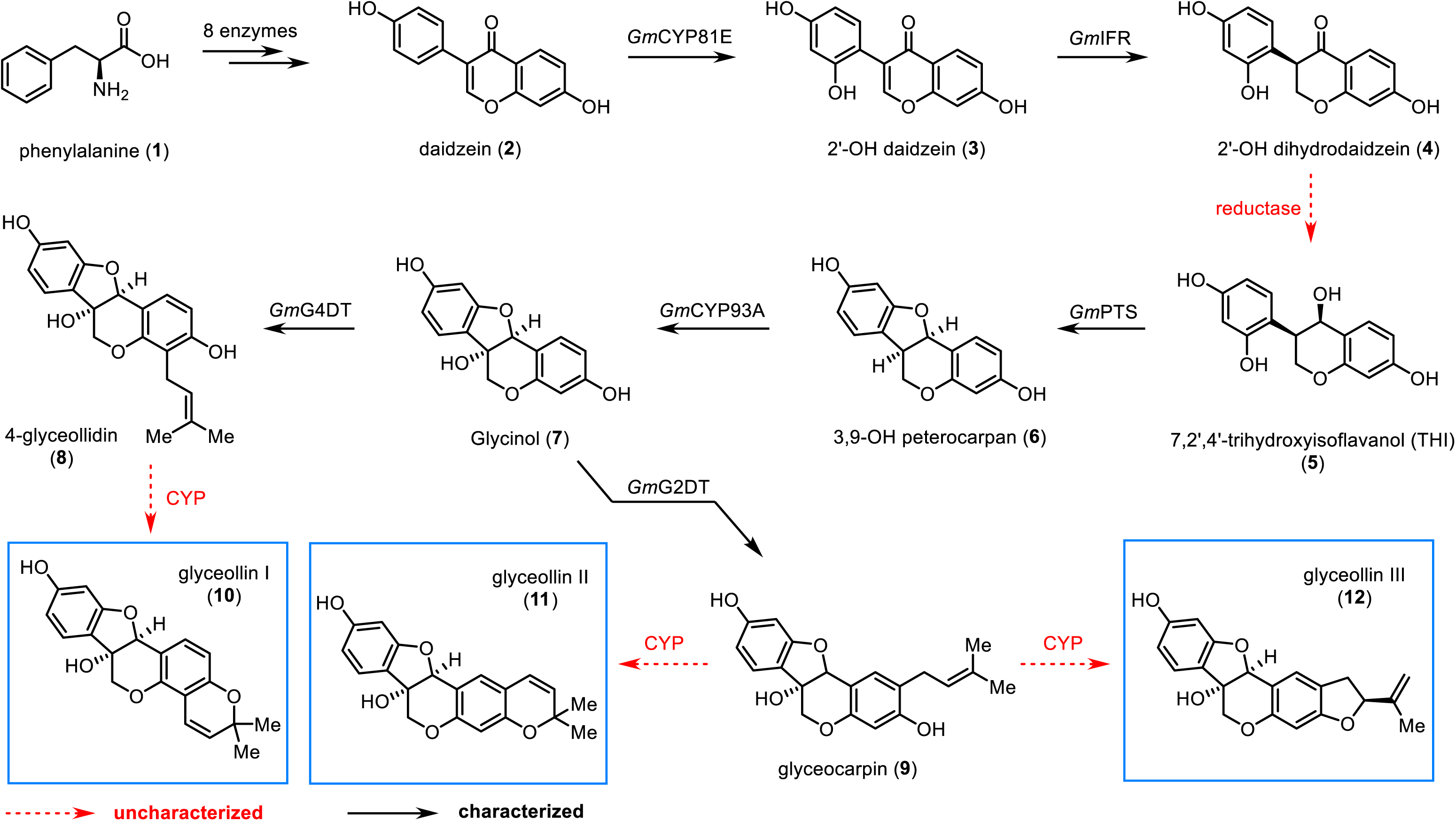
Proposed biosynthetic pathway of glyceollins I, II, and III. Dash arrows represent uncharacterized step, black arrows represent characterized steps.

In this study, we elucidated the complete biosynthetic pathway for glyceollins I, II, and III by identifying two reductases responsible for synthesizing THI and five CYP enzymes that catalyse the final steps of glyceollin biosynthesis. We confirmed these findings through both heterologous expression in *N. benthamiana* and *in vitro* enzyme assays. Leveraging these insights, we successfully reconstituted the entire glyceollin biosynthetic pathway in *Saccharomyces cerevisiae* (baker’s yeast), enabling *de novo* production of glyceollins from simple carbon sources. This platform provides a foundation for optimizing glyceollin production at an industrial scale, paving the way for future agricultural and biological investigations.

## Results

### Identification and characterization of THI synthase (THIS)

The enzyme responsible for converting 2’-OH dihydrodaidzein (**4**) to THI (**5**) in soybean remains unidentified. Similar reductive reactions have been reported with vestitone reductase (*Ms*VR) from *Medicago sativa* and sophorol reductase (*Ps*SR) from *Pisum sativum* (**Supplementary Fig. 1**) (Guo et al., 1994; DiCenzo and VanEtten, 2006). To identify candidate genes encoding this reductase, we performed a BLAST search against the *Glycine max* genome using the protein sequences of *Ms*VR as a query. The search revealed seven candidate genes with sequence identities to *Ms*VR ranging from 63.6% to 79.2% **(Supplementary Table 1)**. To clone these genes, cDNA was prepared from soybean roots treated with Triton X-100, a known inducer of glyceollin biosynthesis (Yoshikawa, 1978). Two of these genes, Glyma.18G220500 and Glyma.09G269600, which share 97.6% identity, were successfully cloned from the cDNA into a binary vector for plant expression. To characterize their function, the genes were transiently expressed in *N. benthamiana* leaves, followed by infiltration with synthetic substrate **4**. Analysis of crude leaf extracts using quadrupole-time-of-flight (QTOF)-HRMS revealed the production of THI (**5**) and 3,9-OH pterocarpan (**6**) on the basis of retention time (Rt) and mass fragmentation patterns (**Figure 2a, 2b, Supplementary Figure 2**). Notably, microsomal supernatants from elicitor-induced soybean cell cultures can catalyse the direct conversion of **4** to **6** (Fischer et al., 1990a). To determine whether Glyma.18G220500 and Glyma.09G269600 might be bifunctional enzymes capable of catalysing both steps (**4** → **5** → **6**), synthetic intermediate **5** was infiltrated into *N. benthamiana* leaves expressing an empty vector. The successful detection of 3,9-OH pterocarpan (**6**) suggested that **5** was spontaneously converted to **6** within the cellular environment or through the action of an endogenous *N. benthamiana* enzyme (**Figure 2a, 2b**). To further characterize the biochemical functions of Glyma.18G220500 and Glyma.09G269600, we expressed and purified *N*-terminal His6-tagged Glyma.18G220500 and Glyma.09G269600 proteins for *in vitro* enzyme assays. These assays confirmed the formation of **5**, with no detectable formation of **6**, suggesting that Glyma.18G220500 and Glyma.09G269600 are specifically responsible for the reduction of **4** to **5** (**Figure 2a, 2c**). These enzymes were therefore named THI synthase 1 (*Gm*THIS1) and THI synthase 2 (*Gm*THIS2). *In vitro* assays revealed that *Gm*THIS1 exhibited higher activity than GmTHIS2 (**Supplementary Figure 3**). The remaining five genes (Glyma.18G220600, Glyma.09G269500, Glyma.09G269400, Glyma.13G203900, and Glyma.12G238100) could not be cloned from cDNA and were instead synthesized and expressed in tobacco for functional analysis. Glyma.18G220600, Glyma.09G269500, Glyma.09G269400, and Glyma.13G203900 were found to catalyse the conversion of 2’-OH dihydrodaidzein (**4**) to 7,2’,4’-trihydroxyisoflavanol (THI, **5**), albeit with lower activity than *Gm*THIS1 and *Gm*THIS2 in tobacco (**Supplementary Figure 4**). Additionally, both *Ms*VR and *Ps*SR were shown to catalyse the conversion of **4** to **5** in the *N. benthamiana* transient expression system and in *in vitro* assays using purified enzymes (**Supplementary Figure 5**). However, chromatograms revealed undesired and uncharacterized byproducts, indicating possible nonspecific activities. These results reveal that *Gm*THIS1 and *Gm*THIS2 exhibit high specificity for 2’-OH dihydrodaidzein (**4**), distinguishing them from vestitone and sophorol reductases, which display nonspecific activity and byproduct formation. The identification of THI synthases (*Gm*THIS1 and *Gm*THIS2) underscores the evolutionary diversification of leguminous reductases in isoflavonoid biosynthesis, providing new insights into the enzymatic specialization within this metabolic pathway.

**Figure 2:**
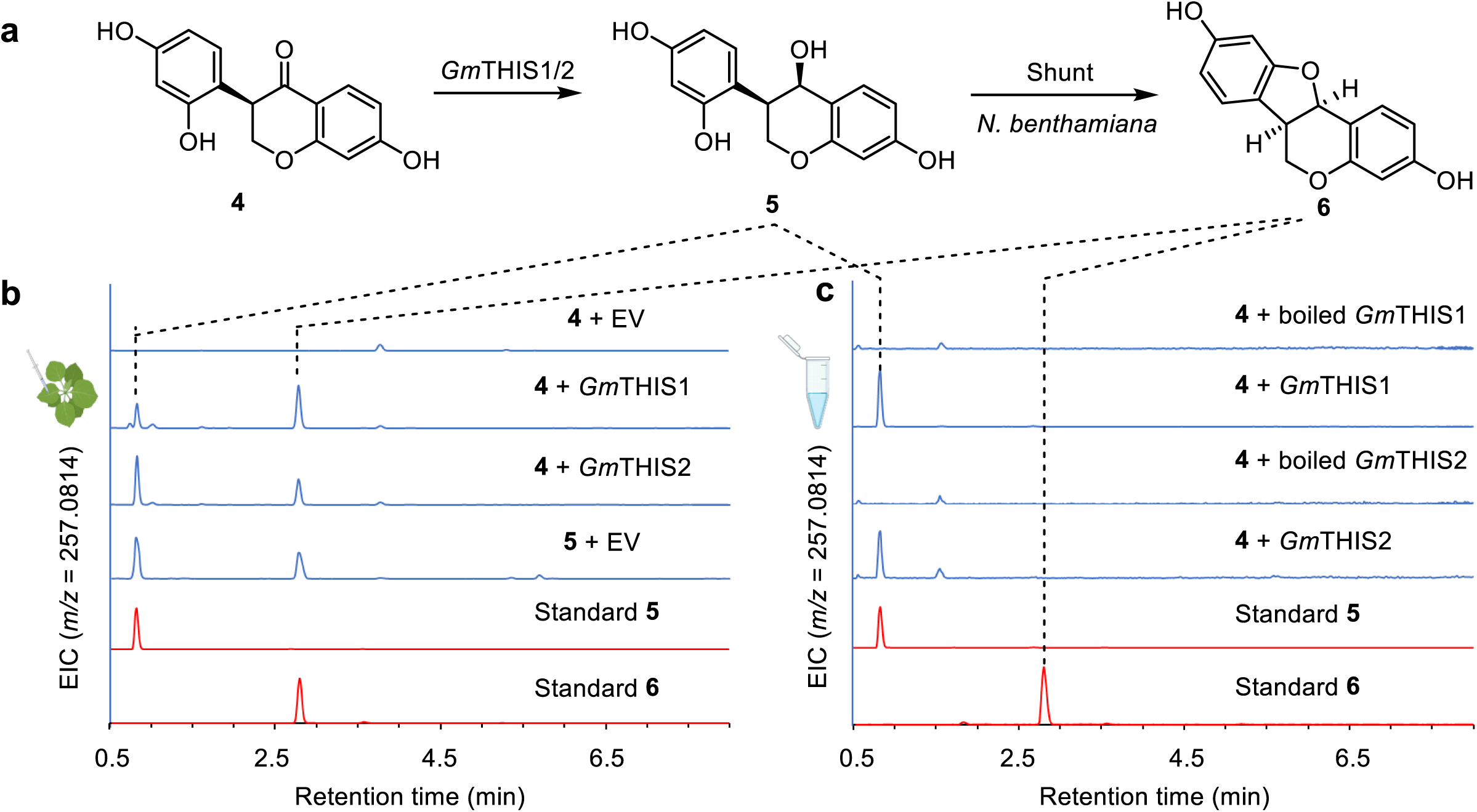
Functional characterization of *Gm*THIS1/2. (a) Scheme of the conversion from **4** to **5**, where the step from **5** to **6** represents a shunt in *N. benthamiana*. (b) Transient expression of *Gm*THIS1/2 with the substrate **4**. Extracted ion chromatograms (EICs) for **5** ([M-H_2_O+ H]^+^ = *m/z* 257.0814) and **6** ([M + H]+ = *m/z* 257.0814) are displayed. (c) *In vitro* experiments of *Gm*THIS1/2 using purified proteins. EICs for **5** and **6** are displayed.

### Identification and characterization of glyceollin I synthases (GISs)

The final steps in the biosynthesis of glyceollins I, II, and III are proposed to involve oxidation of the prenyl unit followed by cyclization, catalysed by cytochrome P450 enzymes, to form 5- and 6-membered rings. As most biosynthetic genes are upregulated upon infection by *Phytophthora sojae* (**Supplementary Table 2**), we used publicly available transcriptomic data from soybean hairy roots treated with a wall glucan elicitor (WGE) from *P. sojae* (Jahan et al., 2020; Khatri et al., 2022) to identify candidate CYP genes. From this dataset, 28 upregulated CYP genes whose expression levels were at least twofold greater than those of the control were selected for further characterization (**Supplementary Table 3**). We cloned these genes from Triton X-100-induced soybean cDNA and transiently expressed them in *N. benthamiana* leaves. When two of these candidates, Glyma.11G062500 and Glyma.11G062600, were expressed in the presence of synthetic 4-glyceollidin (**8**), a new peak with an *m/z* (M+H)^+^ = 339.12 and a retention time (Rt) = 6.57 was observed. This peak, which was absent in the empty vector control, along with a fragment ion at *m/z* (M+H-H_2_O)^+^ = 321.11, matched the synthetic standard of glyceollin I (**10**) (**Figure 3a, Supplementary Figure 6**). Therefore, we named these two CYPs glyceollin I synthases 1 and 2 (*Gm*GIS1 and *Gm*GIS2). To confirm their functions, we conducted feeding experiments in the *S. cerevisiae* WAT11 strain using substrate **8**. While *Gm*GIS1 and *Gm*GIS2 share 87.3% protein sequence identity and exhibit comparable activity in *N. benthamiana*, glyceollin I (**10**) was undetectable when *Gm*GIS1 was expressed in *S. cerevisiae* (**Figure 3a**). To figure out the potential cause of the loss of function of *Gm*GIS1 in yeast, we examined the subcellular localization of *Gm*GIS1 and *Gm*GIS2 via confocal microscopy. The results revealed that *Gm*GIS1 was not localized to the endoplasmic reticulum (ER), while *Gm*GIS2 showed partial ER localization in yeast (**Supplementary Figure 7**).

**Figure 3:**
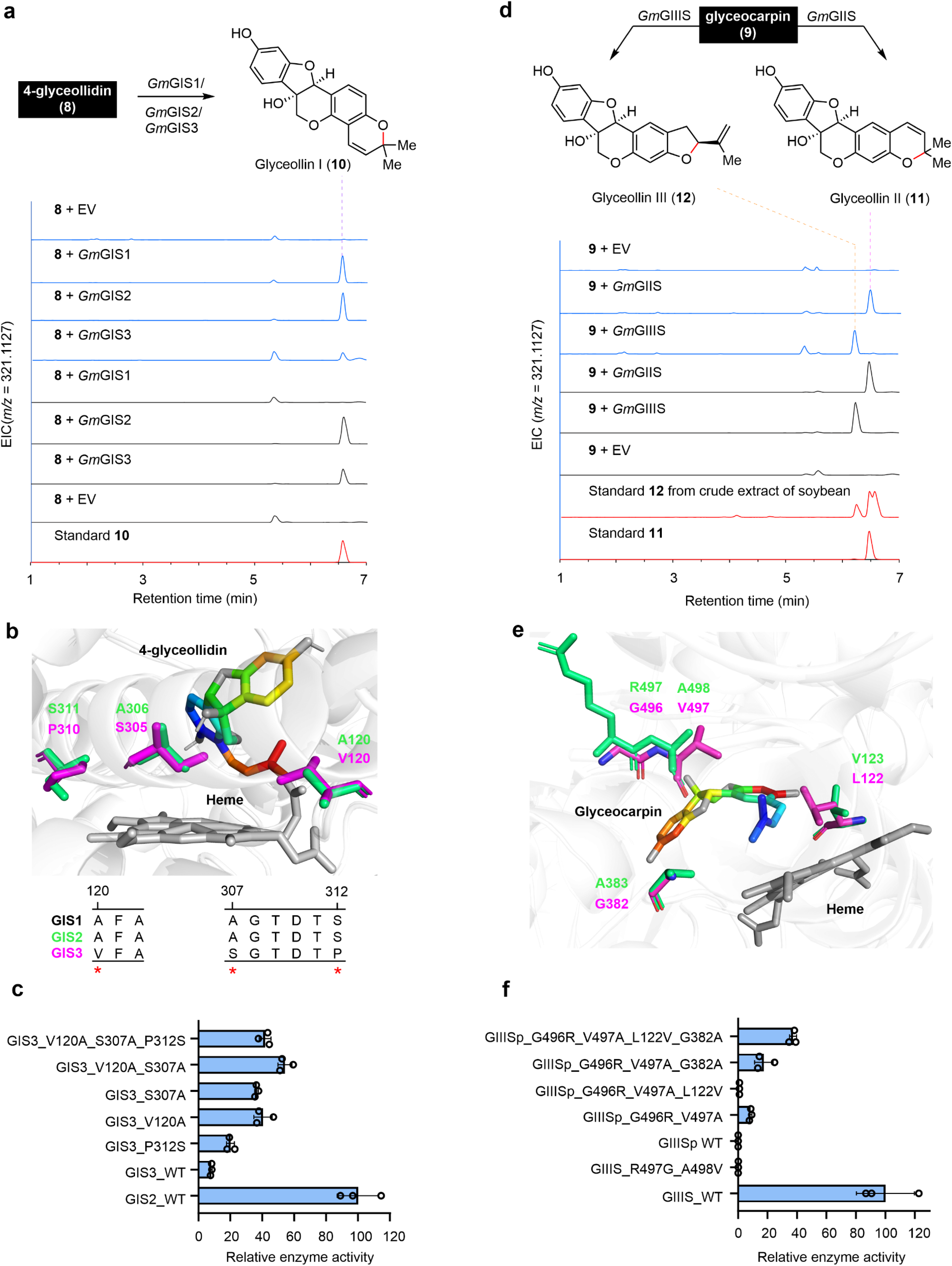
Functional characterization of *Gm*GIS1/2/3 and *Gm*GIIS, *Gm*GIIIS, *Gm*GIIISp. (a) Transient expression in *N. benthamiana* (blue) and feeding experiments in *S. cerevisiae* (black) showing the conversion of **8** to **10**.EIC for **10** ([M-H_2_O+ H]^+^ = *m/z* 321.1127) is displayed. (b) A predicted model of *Gm*GIS2 and *Gm*GIS3, highlighting the interacting substrates and cofactors. (c) Comparison of activity for *Gm*GIS2, *Gm*GIS3, and their mutants. Average LC-MS peak area of product **10** are shown (d) Transient expression in *N. benthamiana* (blue) and feeding experiments in *S. cerevisiae* (black), showing *Gm*GIIS and *Gm*GIIIS converting **9** into **11** and **12**, respectively. EICs for **11** ([M-H_2_O+ H]^+^ = *m/z* 321.1127) and **12** ([M-H_2_O+ H]^+^ = *m/z* 321.1127) are presented (e) A predicted model of *Gm*GIIIS and *Gm*GIIISp, focusing on the substrates and cofactors. (f) Comparison of activity for *Gm*GIIIS, *Gm*GIIISp, and their mutants. Average LC-MS peak area of product **12** are shown. For bar graphs, plotted values represent the mean value ± standard deviations (n = 3 biologically independent samples)

We identified a homologue of *Gm*GIS1/2 in the soybean genome, Glyma.01G179700, which shares 94.0% identity with *Gm*GIS2 and 85.0% identity with *Gm*GIS1 (**Supplementary Figure 8**). Notably, this gene is not upregulated upon WGE treatment (**Supplementary Table 2**) and could not be cloned from cDNA. To investigate its potential function, we synthesized the gene and transiently expressed it in *N. benthamiana*. Surprisingly, Glyma.01G179700 (designated *Gm*GIS3) exhibited significantly lower activity than *Gm*GIS2 in this system (**Figure 3a**). Protein sequence comparisons of *Gm*GIS1, *Gm*GIS2, and *Gm*GIS3 revealed amino acid variations at 15 positions, with *Gm*GIS3 differing from the other two (**Supplementary Figure 8**). Homology models revealed that *Gm*GIS3 possesses three distinct residues within its active site (V120A, S307A, and P312S) compared with *Gm*GIS1 and *Gm*GIS2 (**Figure 3b**). These differences may contribute to the decreased activity observed in *Gm*GIS3. We hypothesized that these natural mutations altered the volume of the active site pocket, affecting substrate accessibility and the reaction process. To test this hypothesis, we performed site-directed mutagenesis to replace V120, S307, and P312 in *Gm*GIS3 with residues found in *Gm*GIS1/2 (A120, A307, and S312) either individually or in different combinations (**Figure 3b, Supplementary Figure 9**). Transient expression of the mutants in tobacco revealed that, compared with wild-type *Gm*GIS3, the single-site mutant *Gm*GIS3_P312S resulted in a twofold increase in product formation. In contrast, the *Gm*GIS3_V120A mutant presented a sixfold increase, whereas the GmGIS3_S307A mutant presented a fivefold increase. Notably, compared with wild-type *Gm*GIS3, the dual-site mutant GmGIS3_V120A-S307A resulted in an approximately eightfold increase in product formation. (**Figure 3c, Supplementary Figure 9**). Finally, the triple-site mutation (*Gm*GIS3_V120A-S307A-P312S) also enhanced product formation, although to a lesser extent than the *Gm*GIS3_V120A-S307A variant did (**Figure 3c, Supplementary Figure 9**). The functional characterization of cytochrome P450 enzymes, including *Gm*GIS1, *Gm*GIS2, and their paralogue *Gm*GIS3, revealed their pivotal roles in the final oxidative cyclization leading to glyceollin I (**10**). Notably, the decreased activity of *Gm*GIS3, compared with that of *Gm*GIS1 and *Gm*GIS2, underscores the functional divergence among these paralogues. The successful restoration of *Gm*GIS3 activity through site-directed mutagenesis of key active site residues provides valuable insights into the structural determinants of enzymatic function, paving the way for future protein engineering efforts.

### Identification and characterization of glyceollin II (GIIS) and III (GIIIS) synthases

Glyceollins II (**11**) and III (**12**) are isomeric pterocarpinoid phytoalexins that share a characteristic 6α-hydroxy-pterocarpan ring system. The distinction between these two compounds lies in the arrangement of the C2 prenyl group in the biosynthetic precursor glyceocarpin (**9**), which leads to the formation of distinct heterocyclic structures: a dihydropyran ring in glyceollin II (**11**) and a hydrofuran ring in glyceollin III (**12**). Transient expression of Glyma.01G135200 with synthetic glyceocarpin (**9**) as a substrate revealed a peak at Rt = 6.48 min that coeluted with the synthetic standard of glyceollin II (**11**) (**Figure 3d, Supplementary Figure 10**). Therefore, Glyma.01G135200 was named glyceollin II synthase (*Gm*GIIS). Transient expression of Glyma.13G285300 in *N. benthamiana* combined with infiltration of glyceocarpin (**9**) resulted in the emergence of a novel peak at Rt = 6.25 min, which was distinct from those of glyceollins I and II. Mass spectral analysis of this peak revealed a fragmentation pattern consistent with that of glyceollin I or II, with a parent ion at *m/z* (M+H)^+^ = 339.12 and a fragment ion at *m/z* (M-H_2_O+H)^+^ = 321.11 (**Figure 3d**, **Supplementary Figure 11**). We proposed this compound as glyceollin III, as its retention time corresponds to the peak detected in the LC MS data from elicitor-induced soybean roots (**Supplementary Figure 11, 12**) and aligns with previously reported data in the literature (Jahan et al., 2020; Khatri et al., 2024). Accordingly, we designated Glyma.13G285300 glyceollin III synthase (*Gm*GIIIS). *Gm*GIIS and *Gm*GIIIS share 70.8% protein sequence identity. To further confirm their functions, we conducted feeding experiments in *S. cerevisiae*, which demonstrated their respective roles in the biosynthesis of glyceollins II and III (**Figure 3d**).

The soybean genome contains a homologue of *Gm*GIIIS, Glyma.15G203500, which shares 92.7% protein sequence identity with *Gm*GIIIS and is upregulated in response to *P. sojae* WGE infection (**Supplementary Table 2**). However, transient expression of Glyma.15G203500 in *N. benthamiana* leaves revealed no detectable activity, indicating that it functions as a pseudo-glyceollin III synthase and was designated *Gm*GIIISp. To identify candidate amino acid residues critical for the inactivity of *Gm*GIIISp, we generated a structural model of the enzyme. We found that four amino acids in the binding pocket differed between *Gm*GIIISp and *Gm*GIIIS (**Figure 3e**). These residues from *Gm*GIIIS were introduced into *Gm*GIIISp in an attempt to recover its activity. We performed transient expression assays of the resulting mutants in *N. benthamiana* leaves with coinfiltration with **9**. A single mutation, L122V or G382A, failed to restore activity. However, mutagenesis of residues 496-497 (GV to RA) restored approximately 10% of *Gm*GIIIS activity, indicating the importance of these two amino acids for enzymatic function (**Figure 3f, Supplementary Figure 13**). Indeed, reverse mutations at positions 497-498 (RA497-498GV) in *Gm*GIIIS resulted in a loss of function (**Figure 3f, Supplementary Figure 13**). Further investigation revealed additional mutations in the *Gm*GIIISp-GV496-497RA mutant. Introducing the L122V mutation abolished the residual activity, whereas substituting G382A instead of L122V slightly improved the activity compared to that of *Gm*GIIISp-GV496-497RA. Finally, combining all four mutations (G496R, V497A, L122V, and G382A) into *Gm*GIIISp resulted in the quadruple mutant achieving more than 40% of the activity of *Gm*GIIIS (**Figure 3f, Supplementary Figure 13**). The identification of glyceollin II and III synthases (*Gm*GIIS and *Gm*GIIIS) demonstrated the diversification of cytochrome P450 enzymes in soybean. The regioselective cyclization of glyceocarpin to distinct glyceollin isomers underscores the intricate evolution of enzyme-substrate specificity. Interestingly, *Gm*GIIISp functions as a pseudoenzyme due to critical active site substitutions, providing new insights into the evolutionary trajectory of glyceollin biosynthetic genes. Collectively, these findings offer a molecular basis for further enzyme engineering to increase glyceollin production. While this manuscript was in preparation, an independent study reported the enzymatic activities of *Gm*GIS1/2 and *Gm*GIIIS through *in vitro* assays, and these results were supported by *in planta* gene silencing experiments (Khatri et al., 2024).

### *De novo* biosynthesis of daidzein in yeast

Following the complete elucidation of the glyceollin biosynthetic pathway, we aimed to achieve *de novo* glyceollin biosynthesis in *S. cerevisiae*. Considering the lengthy pathway (16 genes) and the presence of a rate-limiting prenylation step (Li et al., 2015; Luo et al., 2019; Yang et al., 2024), we segmented the whole pathway into three modules: the daidzein module (from *L*-tyrosine to daidzein), the glycinol module (from daidzein to glycinol), and the glyceollin module (from glycinol to glyceollins). Unlike the biosynthetic pathway in soybean, which starts from *L*-phenylalanine, we reconstituted the pathway in yeast starting from L-tyrosine, as it already contains a 4-OH group.

For the daidzein module (**Figure 4a**), we first redirected the metabolic flux towards *L*-tyrosine by integrating copies of *ARO4^K229L^*(the feedback inhibition-insensitive 3-deoxy-D-arabino-heptulosonic acid 7-phosphate synthase mutant) and *ARO7^G141S^* (the feedback inhibition-insensitive mutant of chorismate mutase) while knocking out a competing enzyme encoding the gene *ARO10* (phenylpyruvate decarboxylase). We subsequently inserted expression cassettes containing *FjTAL* (tyrosine ammonia lyase from *Flavobacterium johnsoniae*), *Pc4CL2* (4-coumaroyl-CoA synthase 2 from *Petroselinum crispum*), *PhCHS*∼*GmCHR5* (*Petunia hybrida* chalcone synthase fused with *Glycine max* chalcone reductase CHR5), *GmCHR5* (an extra copy of chalcone reductase CHR5 from *G. max*), and *PhCHI* (chalcone isomerase from *Petunia hybrida*). The resulting strain GYN1 produced 17.12 mg/L isoliquiritigenin, 0.64 mg/L liquiritigenin, and 5.62 mg/L naringenin (**Figure 4b**). The inefficiency of isoliquiritigenin isomerization prompted us to explore other CHI orthologues with increased activity. Liu et al. previously achieved 77.7 mg/L liquiritigenin production using *GmCHIB2* (chalcone isomerase from *G. max*) (Liu et al., 2024). Consequently, we integrated *GmCHIB2* into GYN1, resulting in the construction of strain GYN2 with a notable increase in the liquiritigenin titre (24.57 mg/L, **Figure 4b**). To construct a daidzein-producing strain, we further introduced two downstream genes, *2-HIS* (2-hydroxyisoflavanone synthase) and *HID* (2-hydroxyisoflavanone dehydratase), into GYN2 to generate strains GYN3 and GYN4. The former included an optimized combination of *Ge*2-*HIS* (*Glycyrrhiza echinata*) and *GmHID* (*G. max*) as previously reported (Liu et al., 2024), whereas the latter incorporated *AmIFS* (isoflavone synthase from *Astragalus membranaceus*) and *GmHID*. After one day of induction with galactose, both strains presented similar daidzein (**3**) titres of approximately 24 mg/L, reflecting comparable catalytic efficiency between *Ge*2-HIS and *Am*IFS (**Figure 4b**). Therefore, we chose GYN3 as our platform strain for subsequent studies.

**Figure 4:**
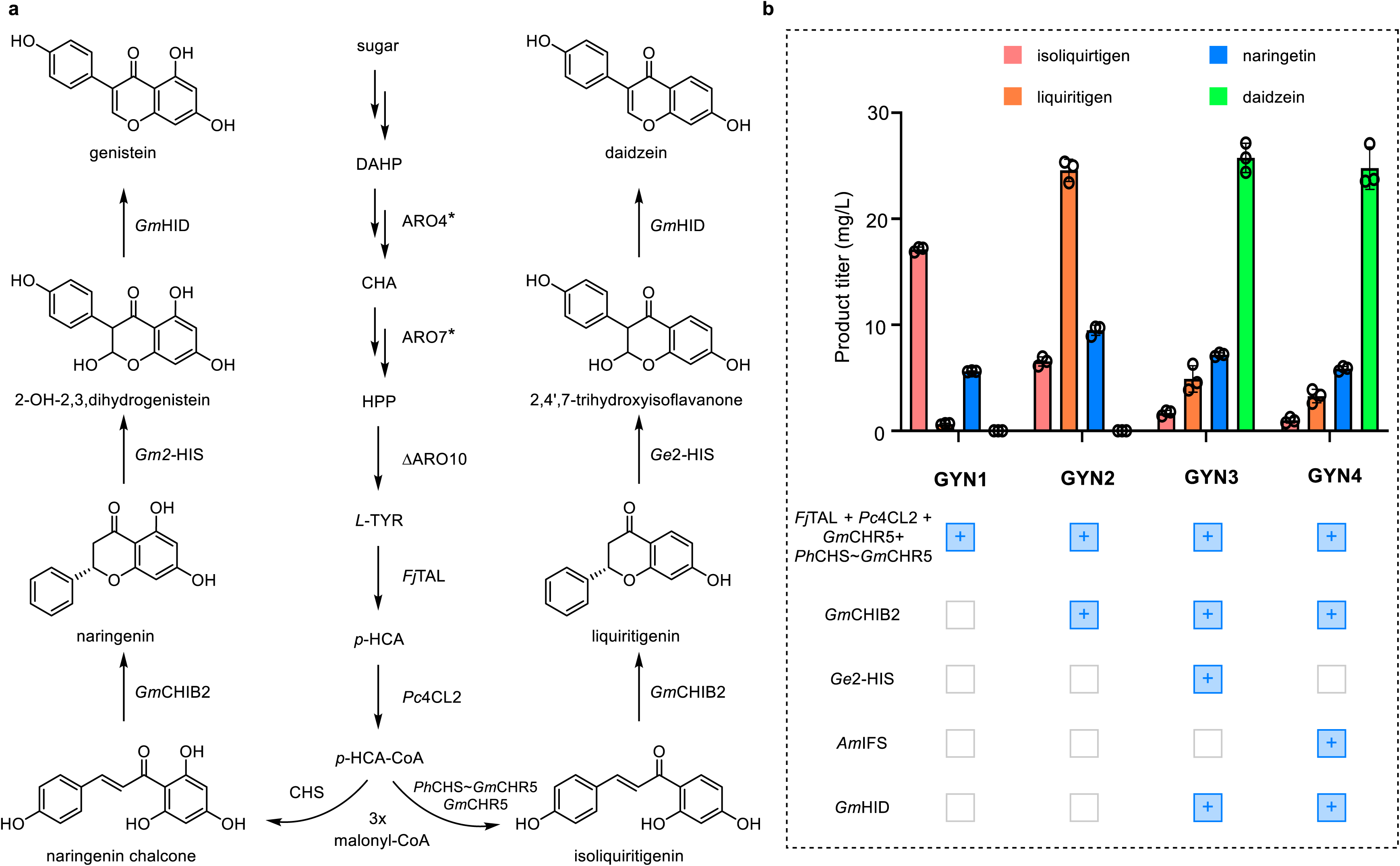
Reconstitution of the daidzein module for *de novo* biosynthesis of daidzein. (a) Schematic diagram illustrating the engineered metabolic pathway for the construction of daidzein-producing strains. (b) Metabolic modification of the daidzein module improved the titres of isoliquiritigenin, liquiritigenin, naringenin, and daidzein. DAHP: 3-deoxy-D-arabino-heptulosonic acid 7-phosphate, CHA: chorismic acid, PPA: prephenate, PPY: phenylpyruvate, HPP: para-hydroxy-phenylpyruvate, *p*-PAC: para-phenylacetaldehyde, L-TYR: L-tyrosine, *p*-HCA: *p*-coumaric acid, *p*-HCA-CoA: p-coumaroyl-CoA, ARO4*: DAHP synthase mutant (K229L), ARO7*: chorismate mutase mutant (G141S), ARO10: phenylpyruvate decarboxylase, *Fj*TAL: *F. johnsoniae* tyrosine ammonia lyase, *Pc*4CL2: *P. crispum* 4-coumaroyl-CoA synthase 2, *Ph*CHS-GmCHR5: *P. hybrida* chalcone synthase fused with *G. max* chalcone reductase, *Gm*CHR5: *G. max* chalcone reductase, *Gm*CHIB2: *G. max* chalcone isomerase, *Ge*2-HIS: *G. echinate* 2-hydroxyisoflavanone synthase, *Am*IFS: *A. membranaceus* isoflavone synthase, *Gm*HID: *G. max* 2-hydroxyisoflavanone dehydratase. Data are shown as mean value ± standard deviations (n = 3 biologically independent samples).

### *De novo* biosynthesis of glycinol in yeast

In the glycinol module (**Figure 5**, **Supplementary Figure 14**), we sequentially integrated *GmCYP81E*, *GmIFR*, *GmTHIS1*, and *GmPTS* expression cassettes into GYN3, resulting in the construction of strains GYN5, GYN6, GYN7, and GYN8 **(Supplementary Table 3)**. As shown in **Figure 5**, most of the daidzein (**2**) in GYN5 was converted to 2’-OH daidzein (**3**), which was subsequently transformed into 2’-OH dihydrodaidzein (**4**) in GYN6. However, the conversion ratio of 2’-OH daidzein (**3**) remained below 50%. Notably, in GYN7, we detected both 7,2’,4’-trihydroxyisoflavanol (**5**) and 3,9-OH pterocarpan (**6**), which is consistent with the results observed in *N. benthamiana* (**Figure 2**). With the introduction of *Gm*PTS, we observed a greater abundance of 3,9-OH pterocarpan (**6**) in GYN8. Ultimately, we integrated the *GmCYP93A* expression cassette into GYN8 to construct strain GYN9, which converted more than 92% of 3,9-OH pterocarpan (**6**) into glycinol (**7**). Given the substantial abundance of glycinol (**7**) in yeast, we proceeded to reconstitute the glyceollin module.

**Figure 5:**
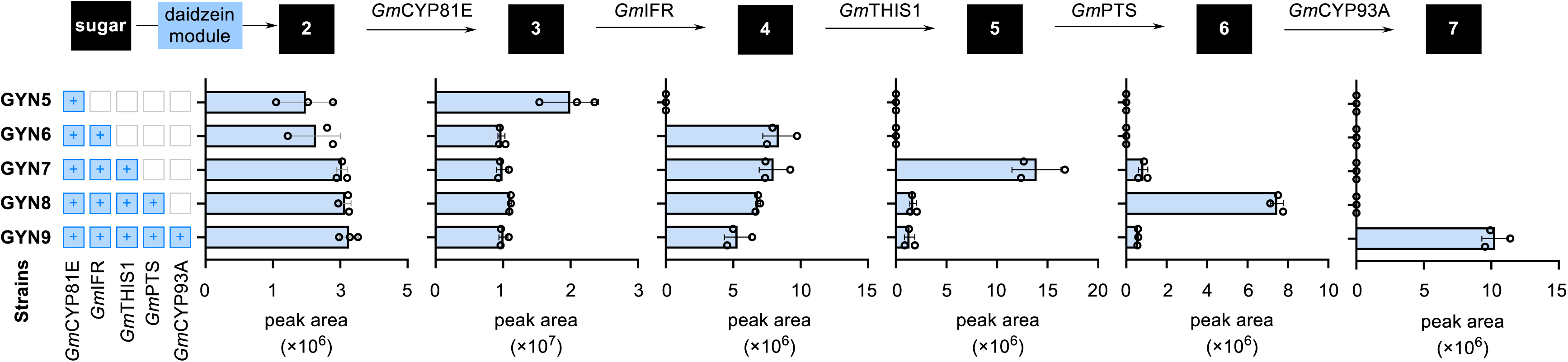
Reconstitution of the glycinol module for *de novo* biosynthesis of glycinol. Scheme of the glycinol module engineering and average LC–MS peak area of the corresponding products produced in yeast strains expressing the indicated enzymes. Data are shown as mean value ± standard deviations (n = 3 biologically independent samples).

### *De novo* biosynthesis of glyceollins in yeast

The glyceollin module encompasses two steps: prenylation and cyclization (**Figure 6a**). Akashi et al. and Yoneyama et al. demonstrated that *Gm*G4DT and *Gm*G2DT catalysed the prenylation of glycinol (Akashi et al., 2009; Yoneyama et al., 2016). Unfortunately, when we introduced both prenyltransferases (PTs) in their full lengths to construct strains GYN11 and GYN10 **(Supplementary Table 4)**, we failed to detect the corresponding prenylated products, 4-glyceollidin (**8**) and glyceocarpin (**9**), in either strain (**Supplementary Figure 15**). Previous studies have suggested that the signal peptides of prenyltransferases might hinder their function in yeast (Li et al., 2015; Sayed et al., 2023; Yang et al., 2024). Therefore, we truncated 44 amino acids from the *N*-terminus of *Gm*G2DT and *Gm*G4DT, leading to the construction of strain GYN12 with truncated *GmG4DT* (*tGmG4DT*) and GYN13 with truncated *GmG2DT* (*tGmG2DT*) (Akashi et al., 2009; Yoneyama et al., 2016). However, prenylated products remained undetectable in these strains (**Supplementary Figure 15**), indicating that prenylation is a major rate-limiting step in our engineered yeast strains.

**Figure 6:**
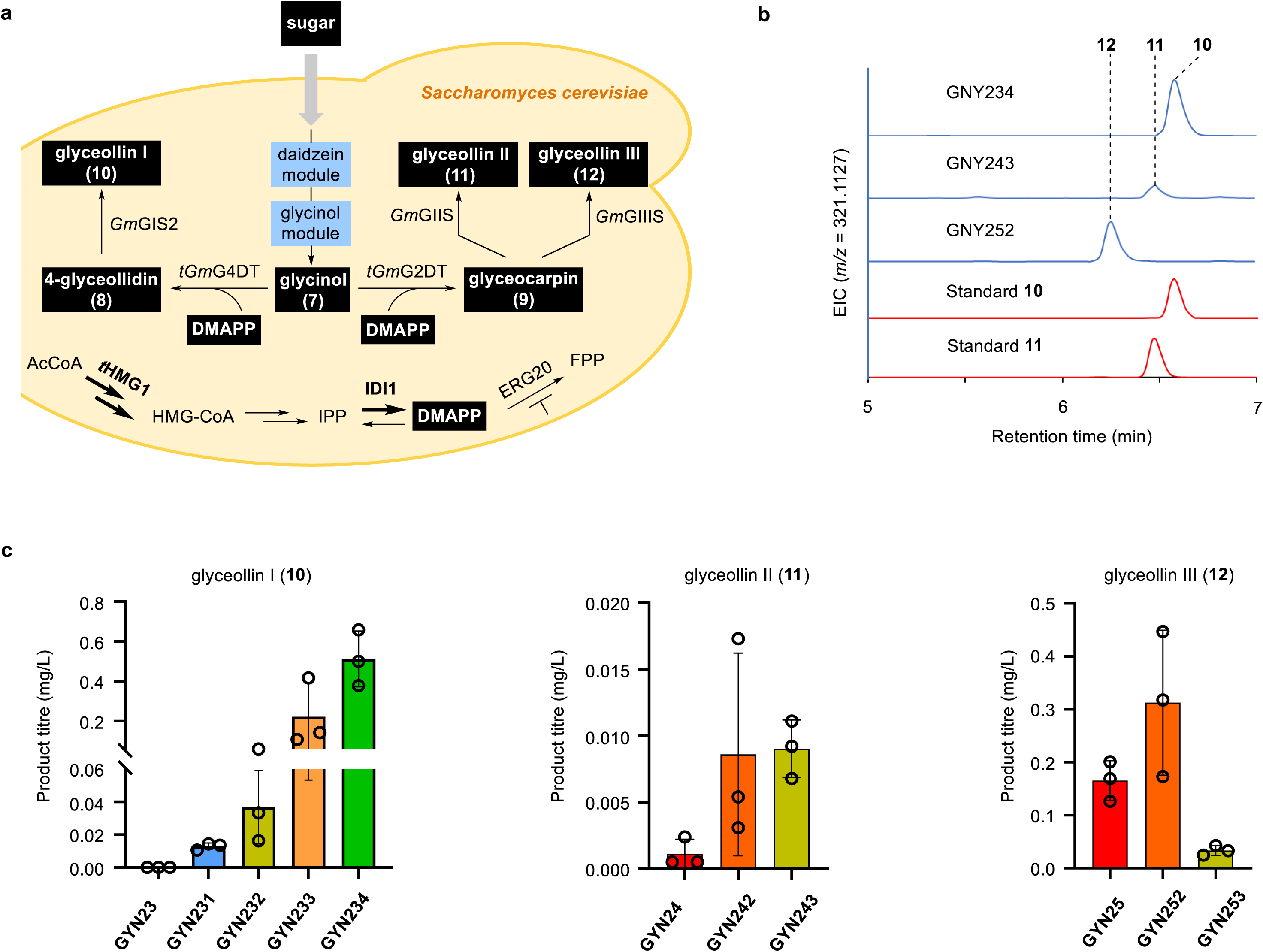
Reconstitution of the glyceollin module for *de novo* biosynthesis of glyceollins. (a) Scheme of the glyceollin module engineering. (b) LC-MS profiles of glyceollins I, II, and III-producing strains GYN234, GYN243, and GYN252. EICs for **10**, **11**, and **12** ([M-H_2_O+ H]^+^ = *m/z* 321.1127) are presented. (c) Changes in the titre of **10**, **11**, and **12** in corresponding glyceollin-producing strains. Data are shown as mean value ± standard deviations (n = 3 biologically independent samples).

To discern whether the instability of prenylated products in yeast contributes to their undetectability, we introduced the last genes, *GmGIS1*, *GmGIS2*, *GmGIIS*, and *GmGIIIS*, resulting in the construction of strains GYN14 to GYN17 (**Supplementary Table 4**), to synthesize the end products (glyceollin I/II/III). However, we could not extract the ion pairs corresponding to any of the end products in these four strains (**Supplementary Figure 16**). When we fed 4-glyceollidin (**8**) or glyceocarpin (**9**) at a final concentration of 1 mg/L, we detected all three types of glyceollins by LC MS, except for the strain containing *Gm*GIS1, despite the titre of glyceollin I catalysed by *Gm*GIS2 remaining low (**Supplementary Figure 17**). These results suggested that *Gm*GIS2, *Gm*GIIS, and *Gm*GIIIS are functional in yeast and that the ultralow titre of the prenylated products is the primary issue.

In the mevalonate pathway, ERG20 is a crucial node that preferentially catalyses dimethylallyl diphosphate (DMAPP) and isopentenyl pyrophosphate (IPP) to form farnesyl pyrophosphate (FPP) in *S. cerevisiae* (**Figure 6a**). It has been reported that decreasing ERG20 expression increases the biosynthesis of prenylated products (Fischer et al., 2011; Li et al., 2015; Zhao et al., 2017; Luo et al., 2019; Yang et al., 2024). To evaluate whether native ERG20 in yeast impaired the prenylation of glycinol (**7**), we transformed both PTs and their truncated versions into strain AM3-1 (Liu et al., 2021), in which the native promoter of *ERG20* (*ERG20p*) was replaced with the dynamically regulated promoter *HXT1p* **(Supplementary Table 4)**. When we fed 0.5 mg/L glycinol (**7**) to AM3-1-expressing PTs, we were able to detect 4-glyceollidin (**8**) and glyceocarpin (**9**), with the truncated PTs demonstrating superior efficiency (**Supplementary Figure 18**). Thus, we replaced the native *ERG20p* in GYN12 and GYN13 with *HXT1p*, generating the yeast strains GYN18 and GYN19, respectively (**Supplementary Table 4**). Consequently, we observed a notable chromatographic peak, with the retention time matching that of the authentic standards for 4-glyceollidin (**8**) and glyceocarpin (**9**), although the ion abundance remained relatively low (**Supplementary Figure 19**).

With respect to cyclization, when we integrated the final genes into GYN18 and GYN19 to construct strains GYN20 with *GmGIS2*, GYN21 with *GmGIIS*, and GYN22 with *GmGIIIS* (**Supplementary Table 4**), we again failed to detect all three glyceollins (**Supplementary Figure 20**). In a previous study, Yang et al. demonstrated that fusing IDI1 with PT significantly increased the titres of the prenylated products (Yang et al., 2024). Accordingly, we constructed strains GYN23, GYN24, and GYN25 by increasing the number of copies of *t*HMG1 and fusing IDI1 with PTs in the relevant strains (**Supplementary Table 4**). As a result, the titres of 4-glyceollidin (**8**) and glyceocarpin (**9**) markedly increased by approximately 22- and 9-fold, respectively (**Supplementary Figure 19**). In the meanwhile, we successfully detected glyceollin III with an estimated titre of 0.06-0.17 mg/L (based on synthetic standards of glyceollins I and II) in strain GYN25 (**Figure 6c, Supplementary Figure 19, 21**). In strain GYN24, we also detected glyceollin II, although its titre (∼1.1 µg/L) was considerably lower than that of glyceollin III (**Figure 6c, Supplementary Figure 19, 22**). In strain GYN23, the titre of glyceollin I was significantly lower (∼0.04 µg/L) than that of glyceollin II and glyceollin III (**Figure 6c, Supplementary Figure 19, 23**), which was consistent with the results of the feeding experiments (**Supplementary Figure 17**). These results indicated that the three P450s were also rate-limiting steps for the *de novo* biosynthesis of glyceollins, especially for *Gm*GIS2 and *Gm*GIIS.

We then explored various metabolic engineering strategies to further enhance the titre of glyceollin, including N-terminal signal peptide engineering and gene copy number engineering. Given that *Gm*GIS, *Gm*GIIS, and *Gm*GIIIS exhibited comparable activities in planta while *Gm*GIIIS exhibited superior catalytic activity in yeast, we proposed the N-terminal signal peptide of *Gm*GIIIS to be more suitable for functional expression of glyceollin synthases (cytochrome P450 enzymes) in yeast. Therefore, we opted to replace the N-terminal signal peptide of *Gm*GIS and *Gm*GIIS with that of GmGIIIS. Based on the signal peptide prediction tool TargetP 2.0 (https://services.healthtech.dtu.dk/services/TargetP-2.0/), *Gm*GIS1 and *Gm*GIS2 possessed identical signal peptide lengths (1-24 amino acids), whereas *Gm*GIIS and *Gm*GIIIS had slightly longer signal peptides (1-35 and 1-29 amino acids, respectively).

As for the biosynthesis of glyceollin I, we substituted the N-terminal signal peptides of *Gm*GIS1 and *Gm*GIS2 with that of *Gm*GIIIS, named as *Gm*GIS1* and *Gm*GIS2*, respectively. The introduction of *Gm*GIS1*, *Gm*GIS2, and *Gm*GIS2* expression cassettes into the GYN23 strain resulted in the construction of strains GYN231, GYN232, and GYN233, respectively (**Supplementary Table 4**). Yeast fermentation demonstrated a significant increase in glyceollin I titre in strains GYN232 and GYN233, with GYN233 achieving a notable titre of up to 0.22 mg/L. Building on this success, we further constructed strain GYN234 by integrating an additional copy of *Gm*GIS2* expression cassette, which further increased the titre of glyceollin I to 0.51 mg/L (**Fig. 6b, 6c**).

For the biosynthesis of glyceollin II, we employed a similar approach. While the replacement of the N-terminal signal peptide failed to yield positive results, the introduction of two additional copies of the original *Gm*GIIS expression cassette significantly enhanced the glyceollin II titre to 0.0090 mg/L (strain GYN243) (**Fig. 6b**, **6c**, **Supplementary Table 4**). Similarly, the integration of an extra copy of *Gm*GIIIS led to an increase in glyceollin III titre, reaching up to 0.31 mg/L (strain GYN252) (**Fig. 6b, 6c, Supplementary Table 4**). Unfortunately, further integration of extra copies of *Gm*GIIIS expression cassette demonstrated an inhibitory effect on glyceollin III biosynthesis (strain GYN253) (**Fig. 6c, Supplementary Table 4**).

Through modular reconstitution and optimization of the glyceollin biosynthetic pathway, we achieved *de novo* biosynthesis of glyceollins I, II, and III from simple carbon sources for the first time. Nevertheless, the titres of glyceollins are still unsatisfactory, particularly those of glyceollin I and glyceollin II, indicating the need for further pathway optimization and metabolic engineering. Considering the accumulation of prenylated products in our engineered yeast strain and the variation in the titre of glyceollins, the last step enzymes (*Gm*GIS2/*Gm*GIIS/*Gm*GIIIS) have become the major bottleneck for efficient biosynthesis. Therefore, protein engineering and chassis engineering should be performed to increase the expression, folding, and/or activity of P450 enzymes (Jiang et al., 2020). For example, *de novo* protein design and amino acid coevolutionary information could be combined for the molecular engineering of P450 enzymes via a structural and data-driven approach (Li et al., 2019). In addition, the microenvironment for P450 activity could be systematically optimized, including overexpressing redox partner genes, increasing the NADPH supply, expanding the endoplasmic reticulum, and regulating iron and haem homeostasis (Li et al., 2019; Cheng et al., 2024). The successful reconstitution and potential further optimization of the glyceollin pathway in yeast demonstrate the feasibility of bulk production of these compounds for functional assays and biotechnological applications.

In summary, using a combination of transient expression in tobacco and *in vitro* enzymatic assays, we successfully elucidated the complete biosynthetic pathway for glyceollins I, II, and III, resolving long-standing gaps in the understanding of these critical soybean phytoalexins. Our discoveries expand the understanding of the precise molecular basis of glyceollin biosynthesis through the identification and characterization of two THI synthases and multiple cytochrome P450 enzymes. Furthermore, by establishing a *de novo* production platform in *S. cerevisiae*, this work represents a significant step towards the sustainable and scalable production of these bioactive compounds. Overall, our findings provide a comprehensive framework for glyceollin biosynthesis, bridging gaps in pathway elucidation and translating these insights into synthetic biology, thereby revealing new opportunities for their application in agriculture, biology, and biotechnology.

## Methods

### Plant materials

Soybean plants were grown in a climate-controlled greenhouse at 24 °C during the day and 18 °C during night, with natural light. *Nicotiana benthamiana* plants were grown in a growth room maintained at 23 ± 2 °C with 16-h day/8-h night light regime.

### Chemicals and molecular biological kits

The solvents used for extraction and chemical synthesis were of HPLC grade, whereas those used for UPLC/HRMS analysis were of MS grade. All solvents were procured from Fisher Scientific, and the chemicals were obtained from Sigma-Aldrich. Information regarding the synthesis of compounds not commercially available can be found in the ‘Synthesis of Compounds’ section. Carbenicillin, gentamycin, spectinomycin, and rifampicin were sourced from Sangon Biotech. Gene and fragment amplifications were conducted using Phanta UniFi Master Mix (Vazyme), whereas colony PCR was performed using Rapid Taq Plus Master Mix (Vazyme). PCR products were purified using the Zymo Research PCR Clean-Up Kit, and plasmid isolation was performed using Promega Wizard Miniprep Kits. RNA was extracted using the FastPure Universal Plant Total RNA Isolation Kit (Vazyme), followed by cDNA synthesis using the Maxima H Minus Kit with dsDNase (Thermo Fisher Scientific). All the restriction enzymes were obtained from NEB.

*Escherichia coli* Trellef® 5α (Tsingke Biotech) or Top10 (Takara) were used (Beijing, China) for the construction of recombinant plasmids. *E. coli* BL21(DE3) was used as the host for heterologous gene expression and recombinant protein purification. LB medium (5 g/L yeast extract, 10 g/L tryptone, 10 g/L NaCl, and 20 g/L agar for solid plates) supplemented with 100 mg/L ampicillin was used for the selection of recombinant *E. coli* strains. *Saccharomyces cerevisiae* WAT11 (*MATa leu2-3, 112 trp1-1 can1-100 ura3-1 ade2-1 his3-11,15*, *AtCPR1*) was used for yeast feeding experiment. *S. cerevisiae* CenA1 containing *Cas9*, *CrCYB5*, and *AtCPR1* was used as the parent strain for the reconstitution of glyceollin biosynthetic pathway. Yeast strains were routinely cultured in YP medium (10 g/L yeast extract, 20 g/L peptone A (meat), and 20 g/L agar for solid plates) supplemented with 20 g/L glucose (YPD) for strain growth or 20 g/L galactose (YPG) for glyceollin fermentation. Drop-out synthetic complete medium (SED-His, SED-Trp, SED-Ura, and SED-His-Ura) containing 1.7 g/L yeast nitrogen base without ammonium and amino acids, 0.6 g/L CSM missing the corresponding nutrients, 1.11 g/L monosodium glutamate monohydrate, 20 g/L glucose, and 20 g/L agar for solid plates was used for the selection of engineered yeast strains.

### Gene cloning

The full-length sequences of the target genes were amplified from *G. max* cDNA using primers listed in Supplementary Table 5. For transient expression in *N. benthamiana*, the PCR products were purified from agarose gel and ligated into a modified 3Ω1 vector, pre-digested with *Bsa*I, using the In-Fusion Kit (Takara). The CCDB resistance marker present in the standard 3Ω1 vector was replaced by candidate genes in the modified version to facilitate easier selection of transformations and decrease false-positive background. Recombinant colonies were selected on LB agar plates containing spectinomycin (100 μg/mL). Positive clones were identified via colony PCR and DNA sequencing.

For heterologous expression in *E. coli* BL21(DE3), the coding sequences (CDS) of *GmTHISs* were amplified from the 3Ω1 constructs. PCR products were purified from agarose gels and ligated into the pOPINF vector, which had been digested with *Hind*III and *Kpn*I, using the In-Fusion Kit (Takara). Recombinant colonies were selected on LB agar plates supplemented with carbenicillin (100 μg/mL). Positive clones were verified by colony PCR and DNA sequencing. Verified plasmids were subsequently transformed into BL21(DE3) competent cells for heterologous expression.

For *de novo* biosynthesis, the daidzein pathway genes (*GmCHIB2*, *Ge2-HIS*, *AmIFS*, and *GmHID*) were codon-optimized for expression in yeast and chemically synthesized by GenScript (Nanjing, China). The glyceollin pathway genes (*GmCYP81E*, *GmIFR*, *GmTHIS1*, *GmPTS*, *GmCYP93A*, *GmG2DT*, *GmG4DT*, *GmGIS1*, *GmGIS2*, *GmGIIS*, and *GmGIIIS*) were amplified from the cDNA of *G. max* and cloned into the linearized pESC-HIS plasmid using Gibson assembly. The sgRNA expression plasmids were constructed in our previous study (Liu, T. et al. 2021).

The synthesis of all primers and DNA sequencing was conducted by Tsingke or Sangon Biotech. The sequences of all primers used in this study are listed in Supplementary Table 5, and the corresponding details on genes and gRNA are provided in Supplementary Tables 6 and 7. All information regarding the plasmids constructed and the sequences of codon-optimized genes are shown in Supplementary Tables 8 and 9.

### Transient expression in *N. benthamiana* and sample harvest

*Agrobacterium* strains carrying the gene constructs were cultured in 8 mL of LB medium supplemented with antibiotics at the following concentrations: 50 μg/mL rifampicin, 50 μg/mL gentamycin, and 250 μg/mL spectinomycin. Cultures were incubated at 28 °C for 24 hours. Cells were harvested by centrifugation at 3000g for 10 minutes, and the resulting cell pellet was resuspended in 5 mL of infiltration buffer (50 mM MES, 2 mM Na PO , 27.8 mM glucose, 10 mM MgCl , and 100 μM acetosyringone), which was centrifuged again under the same conditions and resuspended in 2 mL of fresh infiltration buffer. For individual strain testing, the *Agrobacterium* suspensions were adjusted to an optical density (OD□□□) of 0.6. After a 2-hour incubation at room temperature, the suspensions were infiltrated into the abaxial side of 4- to 6-week-old *N. benthamiana* leaves using a 1 mL syringe without a needle. At 3 days, 200 µM substrates in water with 1% DMSO was infiltrated into underside side of previously Agrobacterium-infiltrated leaves with a needleless 1-ml syringe. Leaves were harvested 12 hours post-infiltration. Each experiment was tested 3 times. To compare the reactivity of the reductases, leaves were harvested 2 hours after infiltration with substrate **4**.

Leaves from *N. benthamiana* transient expression experiments were weighed and snap frozen in liquid N2 and homogenized on a TissueLyser II (Qiagen) using 2-mm-diameter stainless steel beads, with shaking at 25 Hz for 2 min. MeOH (5 μl per mg) was added and were sonicated for 15 min at room temperature, centrifuged at 15,000 rpm for 5 min to pellet plant debris, and the remaining solvent was filtered through 0.22-μm PTFE filters before analysis by high-resolution LC–MS. Metabolites were identified by comparing the retention times and mass fragments of standard compounds.

### Yeast strain construction

Plasmids were transformed into yeast strains (WAT11 or CenA1) using the PEG/LiAc method. To integrate the glyceollin pathway genes into the yeast genome, heterologous gene expression cassettes with 40 bp homologous arms were amplified by PCR. The PCR products were purified (approximately 500 ng) and transformed alongside with the corresponding sgRNA plasmids (approximately 600 ng) into the Cas9 expression yeast strain. Recombinant yeast strains were selected on synthetic dropout agar plates. To ensure the correct integration of gene cassettes into the corresponding genomic sites, several single colonies were picked for diagnostic PCR verification and DNA sequencing.

### Yeast feeding experiments

Recombinant WAT11 clone was selected and inoculated into 10 mL of SD-glucose medium. After incubation for 36 h, 4 mL of culture was centrifuged. The cell pellet was washed with SD-galactose medium, centrifuged again, and the supernatant was removed. The cells were resuspended in 2 mL SD-galactose medium with 50 µM substrate and incubated for 16 h. The cells were lysed using glass beads and 200 µL of MeOH was added. The samples were vortexed for 10 min, centrifuged, filtered with a 0.2 µm filter to remove solids, and then the filtered sample was loaded onto an HPLC for analysis.

### Yeast fermentation

Genome-edited yeast strains were grown in 1 mL YPD medium for 24 hours in 24-well cell culture plates at 30 °C and 200 rpm. The cells were collected, washed once with sterile water, and then resuspended in 1 mL YPG medium. The strains were cultivated for an additional 24 hours at 30 °C and 200 rpm to induce the expression of heterologous genes and the production of glyceollins.

### Protein expression and purification

Genes cloned into pOPINF vectors (*Gm*THIS) were transformed into BL21(DE3) strain for protein expression and carbenicillin (100 µg/mL) was used for selection. Single colonies grown in LB agar media were picked and grown overnight in 10 mL LB media at 37 °C. 1 mL of the overnight cultures was used to inoculate 100 mL of 2xYT media. The fresh cultures were grown at 37 °C until OD600 reaching 0.6-0.8, followed by induction with 250 µM IPTG and incubation at 18 °C for 16-18 h. For purification, the cells were harvested by centrifugation (10 min at 3200g), resuspended in 10 mL of binding buffer (50 mM tris-HCl pH 8, 50 mM glycine, 500 mM sodium chloride, 20 mM imidazole, 5% v/v glycerol, pH 8) containing 0.2 mg/mL Lysozyme and EDTA free protease inhibitor cocktail (Roche cOmpleteTM) and incubated for 30 min on ice. Cells were lysed by sonication using a Sonics Vibra Cell at 40% amplitude, 2s ON, 2s OFF, 6 min total. The crude lysates were centrifuged at 25,000g for 20 min and the cleared lysates were incubated with 250 μL Ni-NTA agarose beads (Qiagen) for 60 min at 4 °C. Next, the beads were sedimented by centrifugation at 1000g for 2 min and washed 3 times with binding buffer before eluting the proteins with elution buffer (50 mM tris-HCl pH 8, 50 mM glycine, 500 mM Sodium Chloride, 250 mM imidazole, 5% v/v glycerol, pH 8). Dialysis and buffer exchange was performed using Buffer A4 (20 mM HEPES pH 7.5; 150 mM NaCl) in centrifugal concentrators with size exclusion of 10 or 30 KDa depending on the protein size. Proteins were aliquoted in 30 µL, snap-frozen and stored at -80.

### *In vitro* enzyme assays

*Gm*THISs were tested using 2’-OH dihydrodaidzein (**4**) as the substrate. Reaction mixtures consisted of *Gm*THISs (2 μM), substrate (50 μM), and NADPH (500 μM) in a total volume of 100 μL TE buffer (50 mM, pH 7.5). Reactions were incubated at 30 for 1 hours. Negative controls consisted of boiled *Gm*THISs (95 , 10 min) in the reaction mixture. After incubation, the reactions were quenched by addition of 1 volume of MeOH, filtered through 0.22-μm PTFE filters and analyzed by untargeted LC-MS.

### LC-MS analysis

The collection of analytical LCMS data was conducted using a Waters LC-MS system, which consisted of a Waters 2767 autosampler, a Waters 2545 pump, a Phenomenex Kinetex column (2.6 μm, C18, 100 Å, 4.6 × 100 mm) equipped with a Phenomenex Security Guard precolumn (Luna, C5, 300 Å), and with a solvent flow of 0.4 mL·min^−1^. For detection purposes, two instruments were utilized: a Waters ZQ mass detector capable of operating in both ES^+^ and ES^−^ modes within a mass range of 100 to 1000 m/z, and a 996 Diode Array detector with a wavelength range of 210 to 600 nm. In this study, two HPLC solvents were utilized: Acetonitrile (B) with a concentration of 0.045 % formic acid, and water (A) with an additional 0.05 % formic acid. The HR-ESI-MS studies utilized the Agilent 1200 Infinity Series High Resolution Electrospray Ionization Mass Spectrometry (MS) instrument. The mass spectrometer utilized in this study was the maXis ESI TOF instrument from Bruker Daltonics, Agilent Technologies. In the stationary phase, a Waters Acquity UPLC BEH C18 column (Milford, USA; 2.1 x 50 mm, 1.7 μm) was employed. The ultraviolet/visible spectra were recorded within the wavelength range of 200 to 600 nm.

For the detection of metabolites in yeast, 800 µL of the culture was extracted using 600 µL of ethyl acetate via a high-throughput tissue grinder SCIENTZ-48 (SCIENTZ, Ningbo, China) at 40 Hz for 2 min. For prenylated products, extra glass beads (0.1 mm in diameter) were added to the culture, and the cells were disrupted by the tissue grinder at 60 Hz for 4 min before extraction with ethyl acetate. The organic layer was collected by centrifugation at 12,000 g for 5 minutes and subsequently filtered through a 0.22 μm membrane before analysis by the SHIMADZU LC-MS 8045 equipped with an Accucore™ C18 column (2.6 μm, 50 mm x 2.1 mm). For the LC conditions, the mobile phase consisted of phase A (water with 0.1% formic acid) and phase B (methanol with 0.1% formic acid); the gradient was as follows: 0 min, 20% B; 6 min, 90% B; 8 min, 90% B; 8.01 min, 20% B; 12 min, stop. The flow rate was set to 0.3 mL/min, and the column temperature was maintained at 30 °C. The MS parameters included positive mode and multiple-reaction monitoring (MRM) mode. The desolvation line (DL) temperature and heating block temperature were set to 250 °C and 400 °C, respectively; the atomizing gas flow rate was 3 L/min, and the collision-induced dissociation (CID) gas pressure was set to 230 kPa. LabSolutions software (version 5.91) from SHIMADZU was used for data analysis. The ion pairs and collision energy (CE) used for detecting metabolites in MRM mode are detailed in Supplementary Table 6, with three biological replicates performed for each experiment.

### Confocal microscopy analysis

To determine the subcellular localization via confocal microscopy, GFP and mCherry were fused to the C-terminus of the target proteins and EMC1 (an endogenous ER localization protein), respectively. The GFP fusion protein encoding genes were cloned into pESC-HIS, while the mCherry fusion protein encoding gene was cloned into pESC-LEU. The plasmid bearing yeast strains were cultured in 1 mL of SED-His-Leu medium for 12 h and then transferred to 1 mL of SEG-His-Leu medium (containing 2% galactose) for an additional 12 h of cultivation. Cells were harvested by centrifugation and resuspended in 100 µL of phosphate-buffered saline (1x), 5 µL of which was immediately imaged using an Olympus FV3000 microscope.

### NMR analysis

NMR spectra were recorded on 500 MHz Bruker Advance III HD spectrometers (Bruker Biospin GmbH, Rheinstetten, Germany). *d_4_*-methenol and *d_6_*-acetone were used as solvents. The chemical shifts (δ) were referenced to the residual solvent signals at δH 3.31 and δC 49.0 ppm for *d_4_*-methenol, and δH 2.05 and δC 29.8 ppm for *d_6_*-acetone. Spectrometer control and data processing were performed using Bruker TopSpin ver. 3.6.1 software.

## Data availability

Data supporting the findings of this work are available within the paper and its Supplemental Information files. The sequence of genes characterized in this article will be deposited in the NCBI database prior to publication. Correspondence and requests for materials should be addressed to B.H. or J.L.

## Acknowledgements

This research was supported by Zhejiang Provincial Natural Science Foundation of China under Grant No. XHD24B0701 (to B.H.) and National Natural Science Foundation of China under Grant No. 22278361 (to J.Z.). We are grateful for financial support from Westlake University startup, Research Center for Industries of the Future (RCIF) at Westlake University, and Zhejiang Key Laboratory Construction Project (to B. H.). We thank Yuechen Bai (Fudan University) for providing soybean seeds. We thank Westlake University Instrumentation and Service Center for Molecular Sciences (ISCMS) for the facility support and technical assistance. We thank Dr. Yinjuan Chen for assistance with mass spectrometry, Dr. Xingyu Lu and Dr. Xiaohuo Shi for assistance with NMR analysis.

## Author Contributions

Y.S. performed the compounds synthesis, cloning, transient expression, protein-expression and protein-assay experiments; C.C. carried out the metabolic engineering in yeast, C.L. performed the NMR experiments and structural assignments. H.Z. constructed the CYP mutants. J.L. and B.H. designed and directed the work. Y.S., C.C., J.L. and B.H. wrote the manuscript with contributions from all authors. All authors discussed the results and commented on the paper.

