## supplementary information for "Elucidation and *de novo* Reconstitution of Glyceollin Biosynthesis"

### 1. Supplementary Figures and Tables

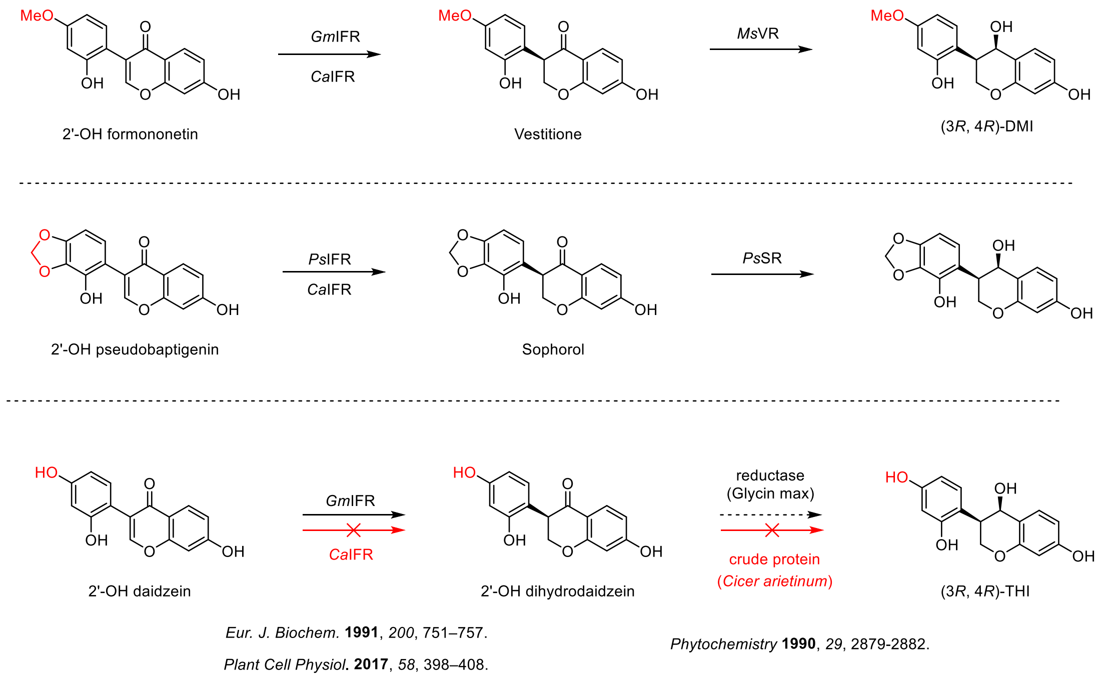

**Supplementary Fig. 1** Reported IFR and VR from legume species including pea (*Pisum sativum*), chickpea (*Cicer arietinum*), alfalfa (*Medicago sativa* L).and soybean (*Glycine max*) (Dixon, R. A. 1993).

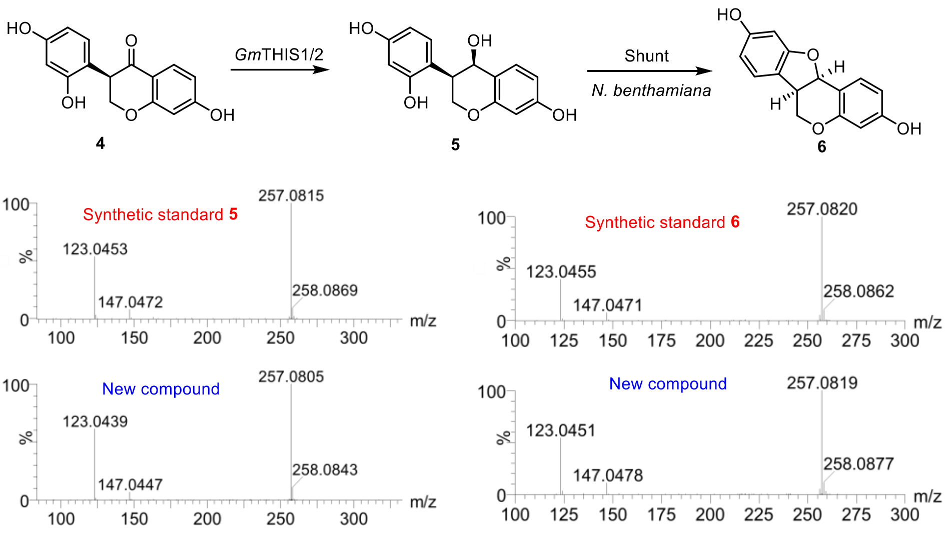

**Supplementary Fig. 2** MS/MS (20 to 50 eV) spectra of generated THI (**5**) and 3,9-OH pterocarpan (**6**) compared to synthetic standards.

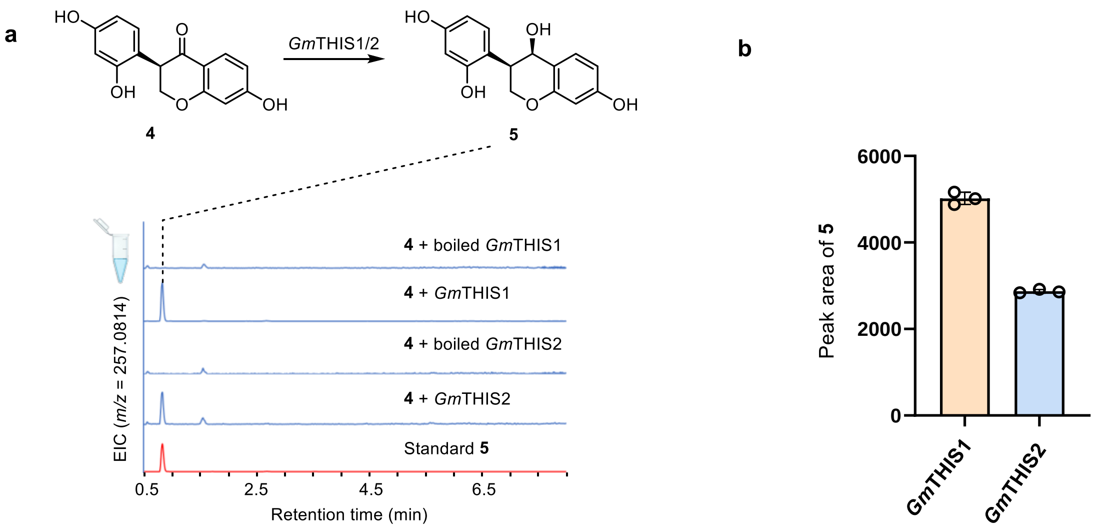

**Supplementary Fig. 3** *In vitro* assays of *Gm*THIS1 and *Gm*THIS2. (**a**) Extracted ion chromatograms (EICs) for **5** ([M-$H_2O$+ H]$^+$ = *m/z* 257.0814) are displayed. (**b**) Peak areas for the formation of **5** after *in vitro* enzyme experiments for *Gm*THIS1 and *Gm*THIS2. Data are shown as mean value ± standard deviations (n = 3 biologically independent samples).

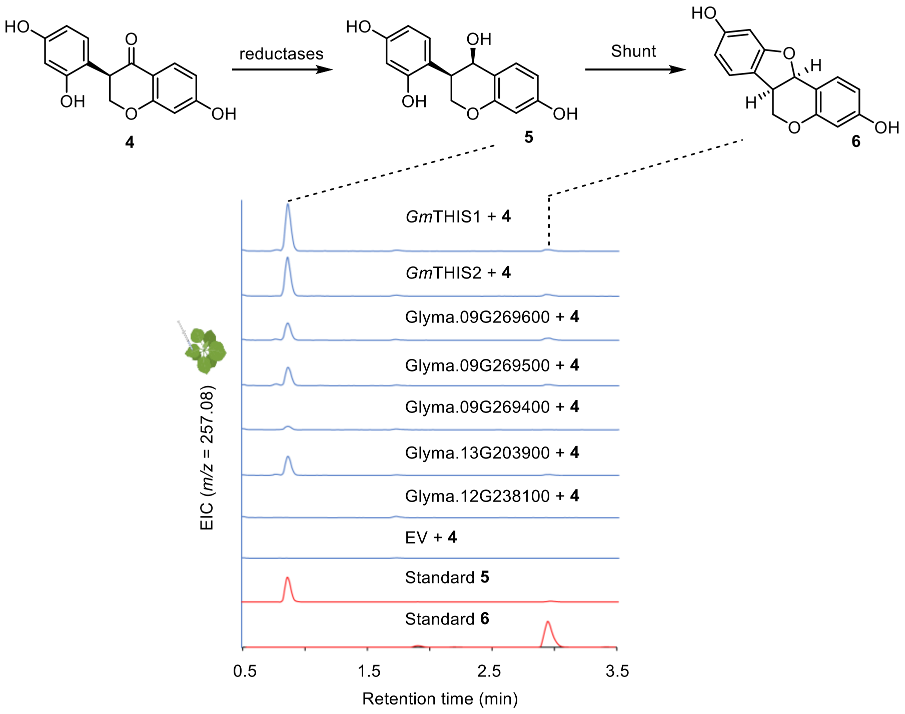

**Supplementary Fig. 4** Functional characterization of reductases activity in tobacco. Substrate **4** was infiltrated into *Nicotiana benthamiana* leaves, and the leaves were harvested after 2 hours and analyzed by LC-MS. Extracted ion chromatograms (EICs) for **5** ($[M\text{-}H_2O+H]^+$ = *m/z* 257.0814) and **6** ($[M+H]^+$ = *m/z* 257.0814) are displayed.

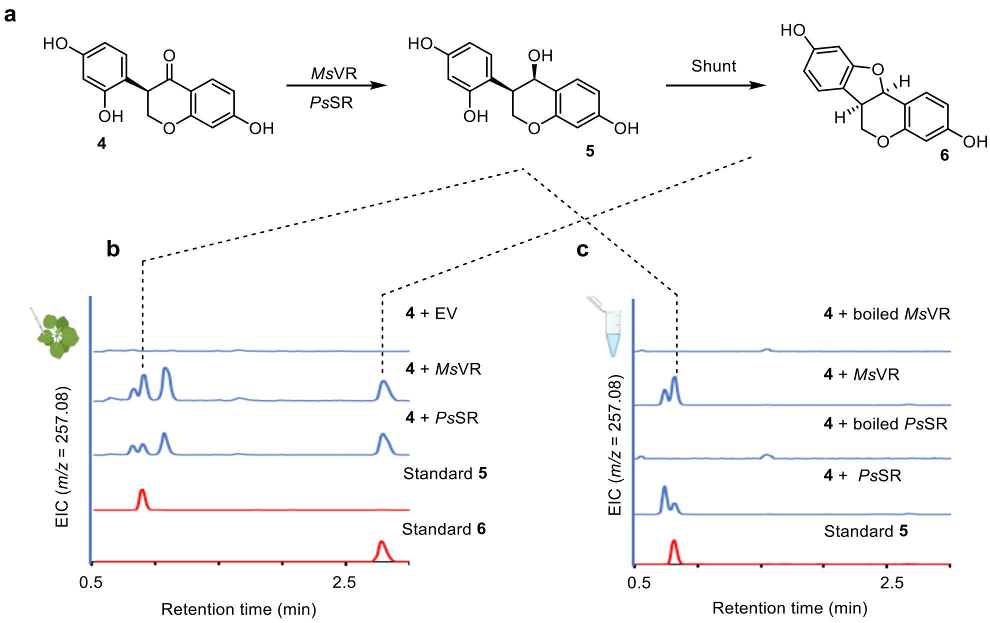

**Supplementary Fig. 5** Transient expression (**b**) and *in vitro* experiment of *Ms*VR and *Ps*SR (**c**) in the presence of substrate **4**. Extracted ion chromatograms (EICs) for **5** ($[M-H_2O+H]^+$ = *m/z* 257.0814) and **6** ([M + H]+ = *m/z* 257.0814) are displayed.

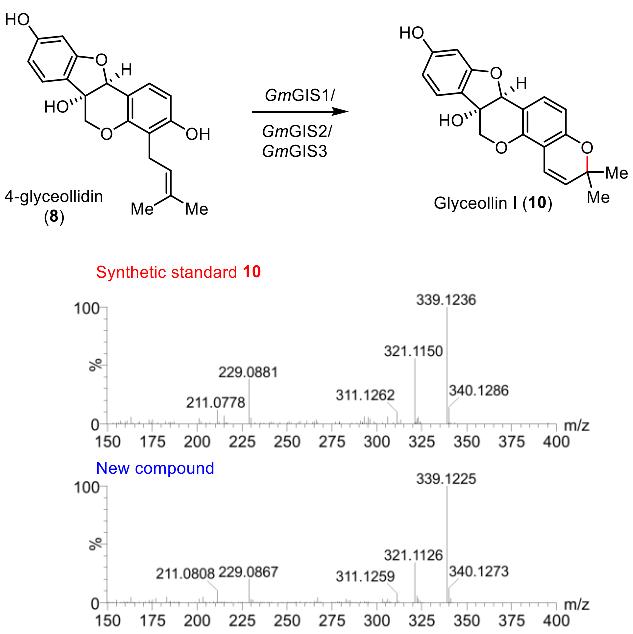

**Supplementary Fig. 6** MS/MS (20 to 50 eV) spectra of generated glyceollin I (**10**) compared to synthetic standard.

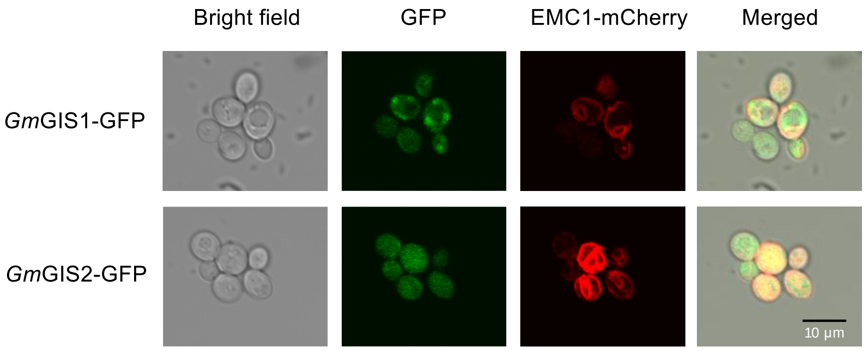

**Supplementary Fig. 7** Subcellular localizations of *Gm*GIS1 and *Gm*GIS2. GFP tag was fused to the C-terminal of *Gm*GIS1 (top) and *Gm*GIS2 (bottom) and co-expressed with EMC1 (ER marker) in yeast, respectively. Left to right: bright field, fluorescent image of GFP, mCherry, merged image of GFP, mCherry and bright field.

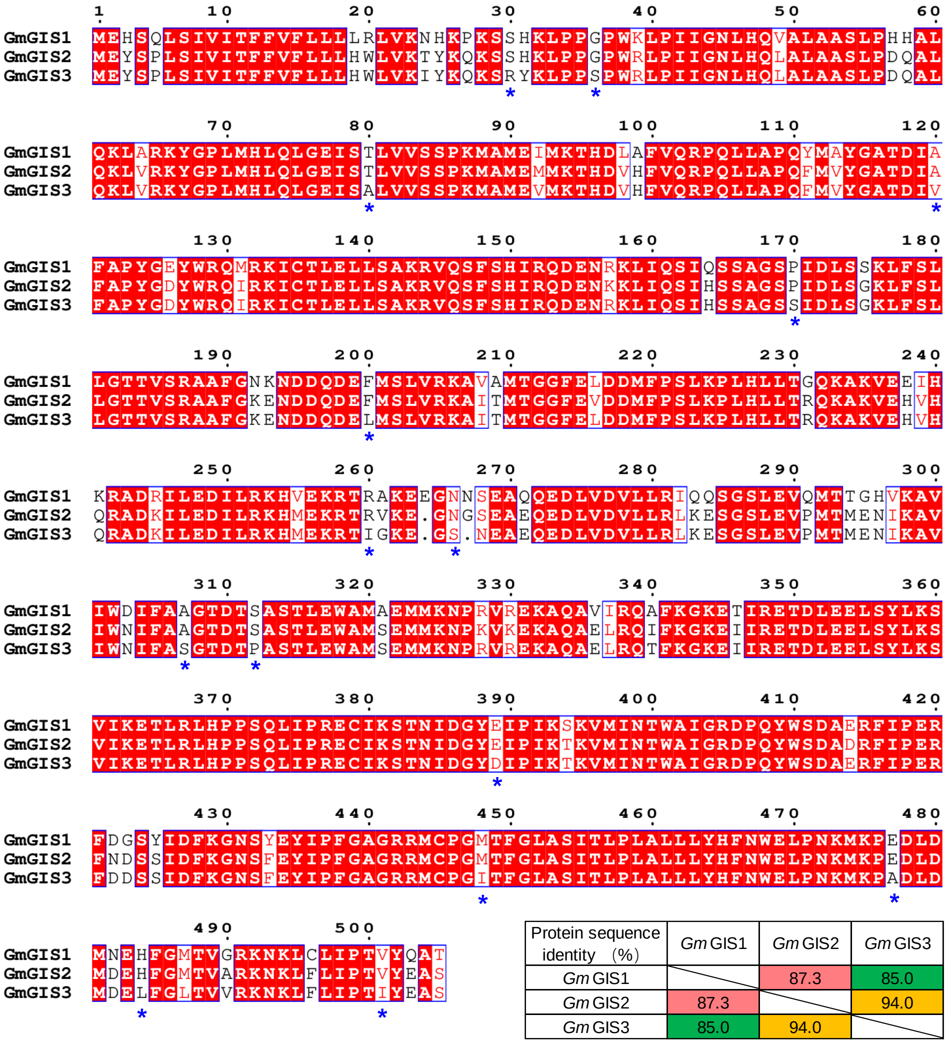

| Protein sequence identity （%） | *Gm* GIS1 | *Gm* GIS2 | *Gm* GIS3 |
|---|---|---|---|
| *Gm* GIS1 | | 87.3 | 85.0 |
| *Gm* GIS2 | 87.3 | | 94.0 |
| *Gm* GIS3 | 85.0 | 94.0 | |

**Supplementary Fig. 8** Protein sequences alignment of *Gm*GIS1, *Gm*GIS2, and *Gm*GIS3.

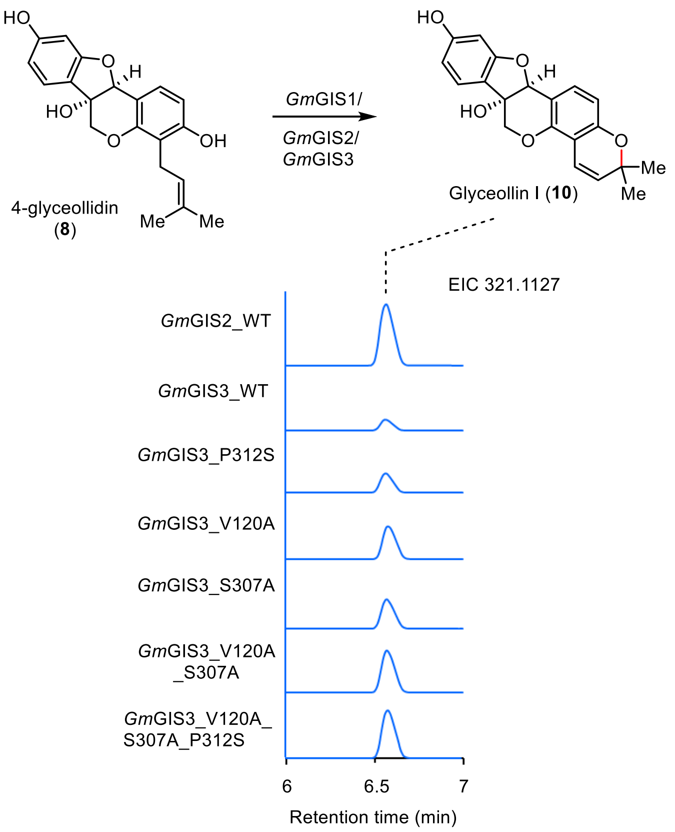

**Supplementary Fig. 9** Functional characterization of *Gm*GIS3 mutants. Transient expression of *Gm*GIS3 mutants in *N. benthamiana* with co-infiltration of 4-glyceollidin (**8**). Extracted ion chromatograms for Glyceollin I (**10**) ($[M-H_2O+H]^+$ = *m/z* 321.1127) are displayed.

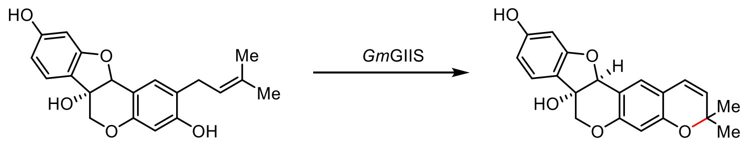

glyceocarpin (**9**) Glyceollin II (**11**)

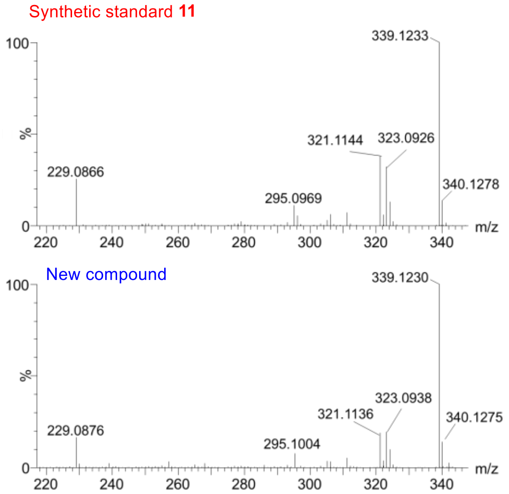

**Supplementary Fig. 10** MS/MS (20 to 50 eV) spectra of generated glyceollin II (**11**) compared to synthetic standard.

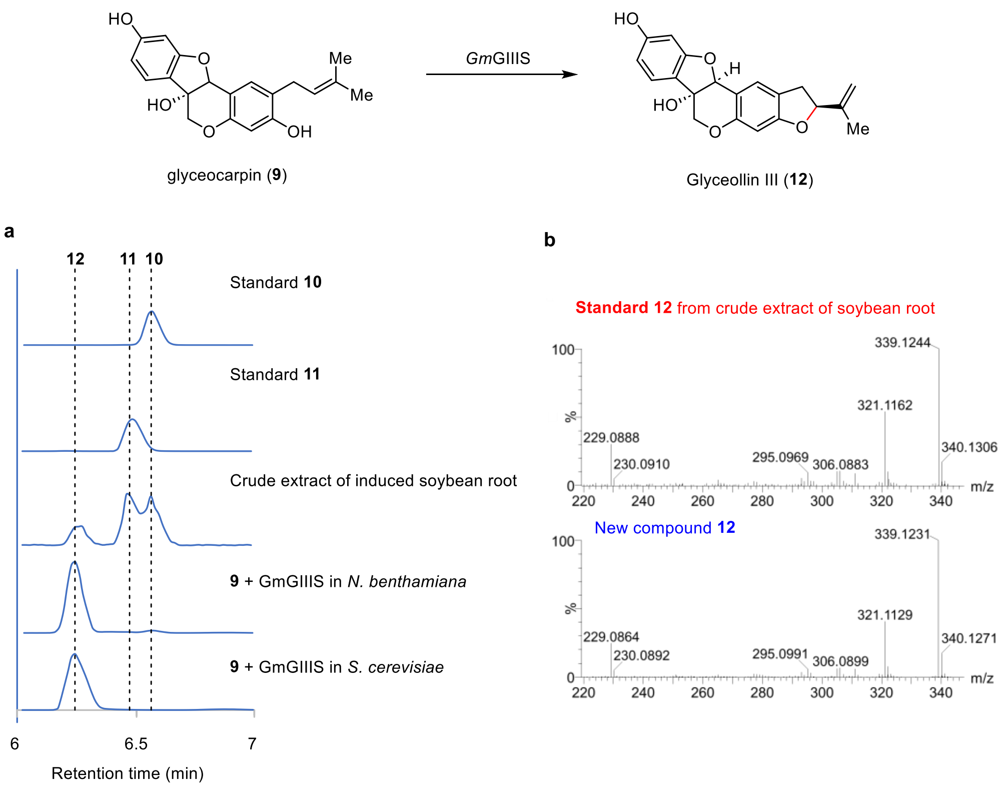

**Supplementary Fig. 11** Chromatograms and MSMS of compound **12**: **a**. Comparison between *Gm*GIIIS experiments including transient expression and feeding experiment and crude extract from Triton-100 induced soybean roots. Extracted ion chromatograms ($[M\text{-}H_2O+H]^+$ = *m/z* 321.1127) for **10**, **11**, and **12** are displayed. **b**. MS/MS (20 to 50 eV) spectra of generated glyceollin III (**12**) compared to standard obtained from crude extract of Triton-100 induced soybean roots.

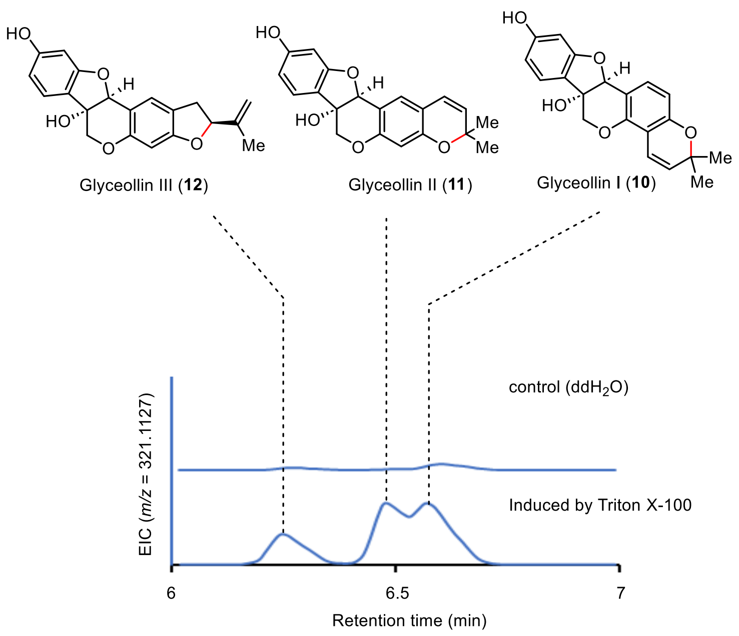

**Supplementary Fig. 12** Induced soybean root by triton X-100 and the control ($ddH_2O$).

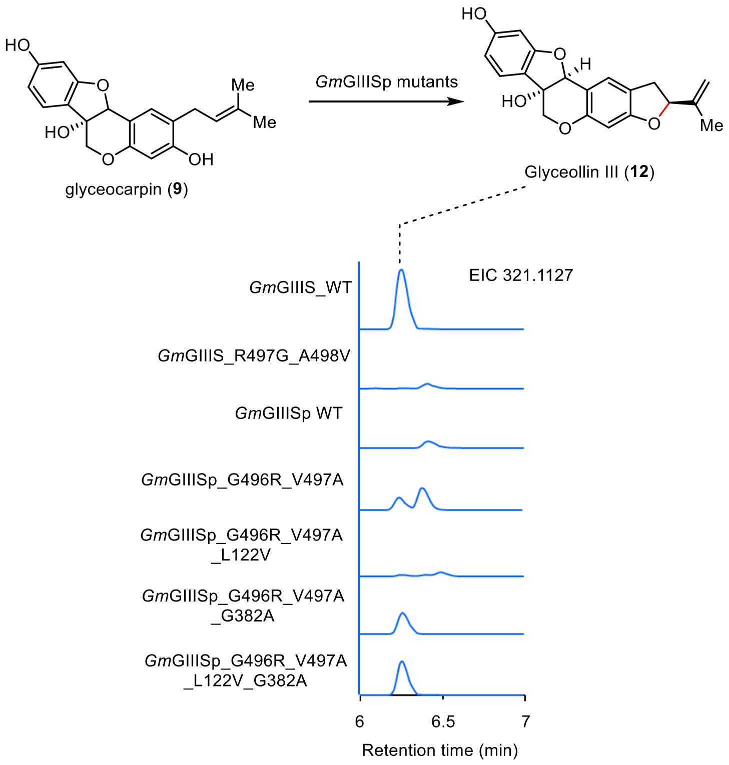

**Supplementary Fig. 13** Functional characterization of *Gm*GIIISp mutants. Transient expression of *Gm*GIIISp mutants in *N. benthamiana* with co-infiltration of glyceocarpin (**9**). Extracted ion chromatograms for Glyceollin III (**12**) ([M-$H_2O$+ H]$^+$ = *m/z* 321.1127) are displayed.

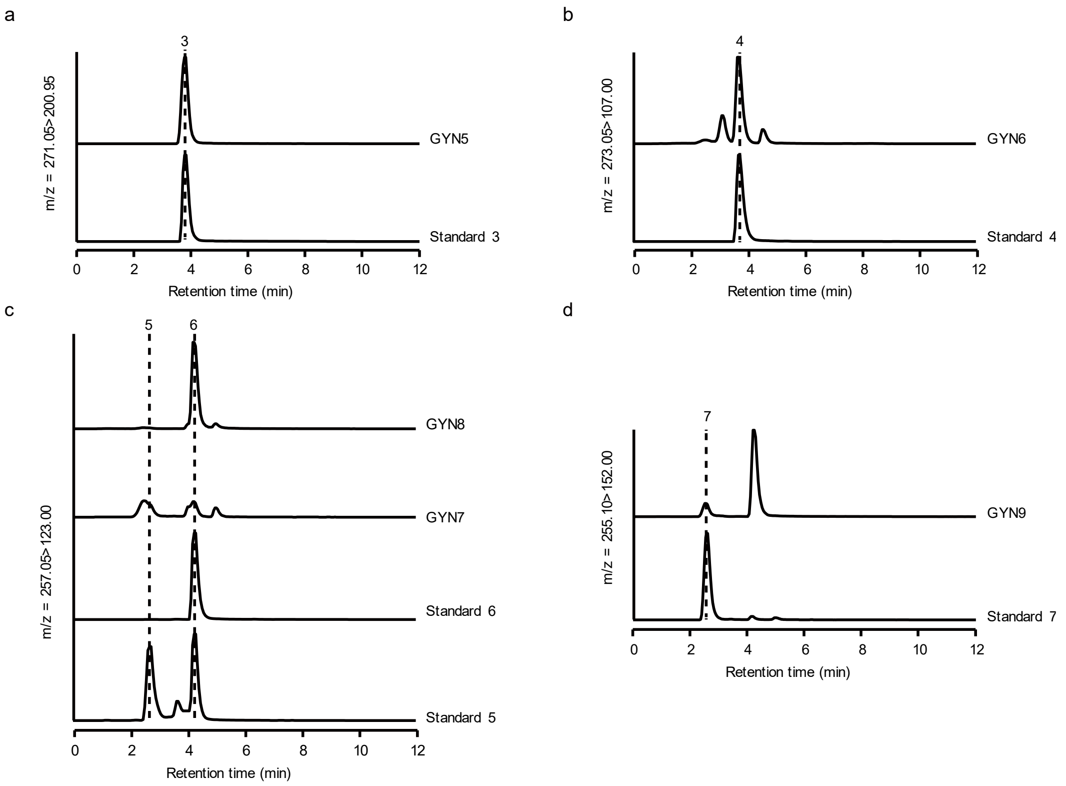

**Supplementary Fig. 14** LC-MS analysis of **3** to **7** in strains GYN5 to GYN9. The MRM spectra for the selected ion pairs are displayed for the standards of **3** to **7**, as well as for the fermented products from strains GYN5 to GYN9.

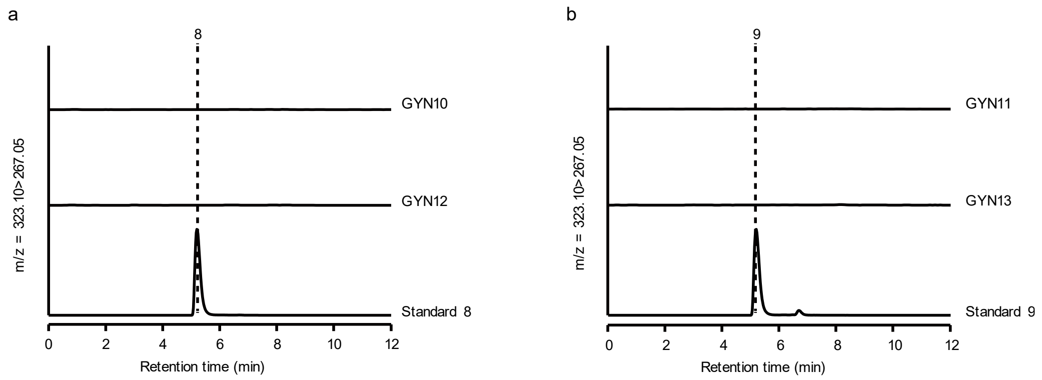

**Supplementary Fig. 15** LC-MS analysis of **8** and **9** in strains GYN10 to GYN13. The MRM spectra for the selected ion pair of *m/z* = 323.10 > 267.05 are shown for the standards of compounds **8** (a) and **9** (b), as well as for the fermented products from strains GYN10 and GYN12 (a), GYN11 and GYN13 (b). Strains GYN10 to GYN13 express *Gm*G4DT, *Gm*G2DT, *tGm*G4DT, and *tGm*G2DT individually.

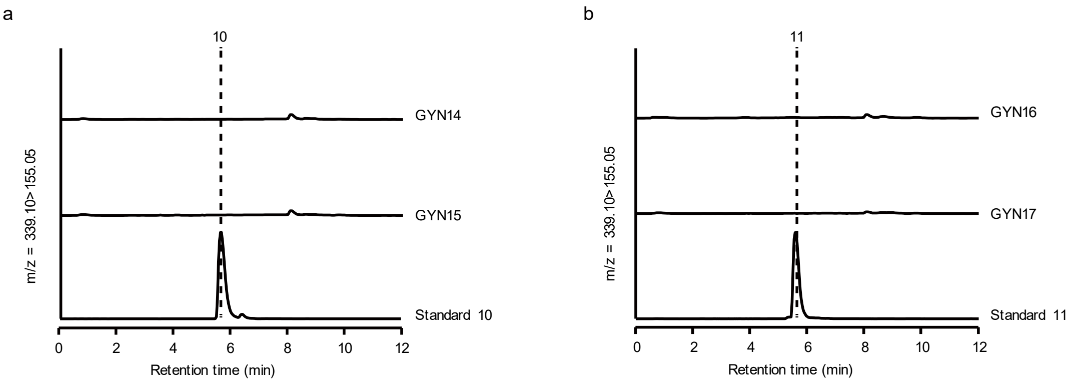

**Supplementary Fig. 16** LC-MS analysis of glyceollin I (**10**), glyceollin II (**11**), and glyceollin III (**12**) in strains GYN14 to GYN17. The MRM spectra for the selected ion pair of *m/z* = 339.10 > 155.05 are displayed for the standards of **10** and **11**, as well as for the fermented products from strains GYN14 to GYN17. These strains express *Gm*GIS1, *Gm*GIS2, *Gm*GIIS, and *Gm*GIIIS individually.

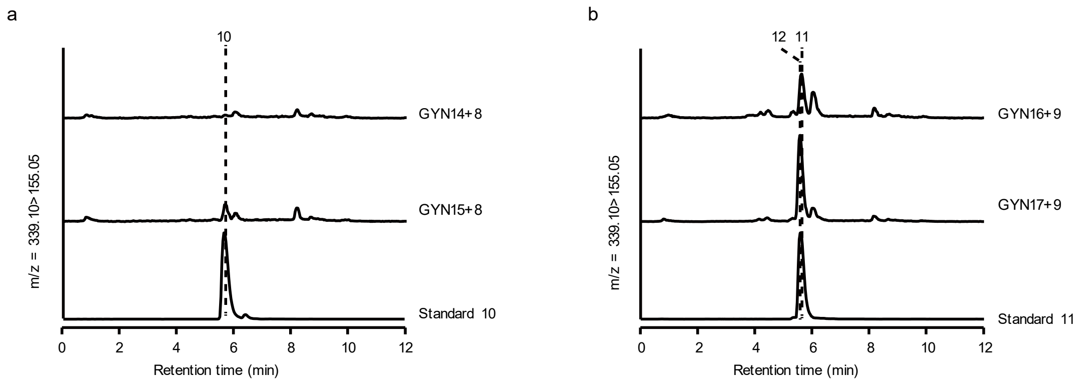

**Supplementary Fig. 17** LC-MS analysis of the fermented products from strains GYN14 to GYN17 following the feeding of 4-glyceollidin (**8**) and glyceocarpin (**9**), respectively. The MRM spectra for the selected ion pair of *m/z* 339.10 > 155.05 are presented for the standards of glyceollin I (**10**) and glyceollin II (**11**), along with the metabolites from strains GYN14 to GYN17. These strains express *Gm*GIS1, *Gm*GIS2, *Gm*GIIS, and *Gm*GIIIS individually.

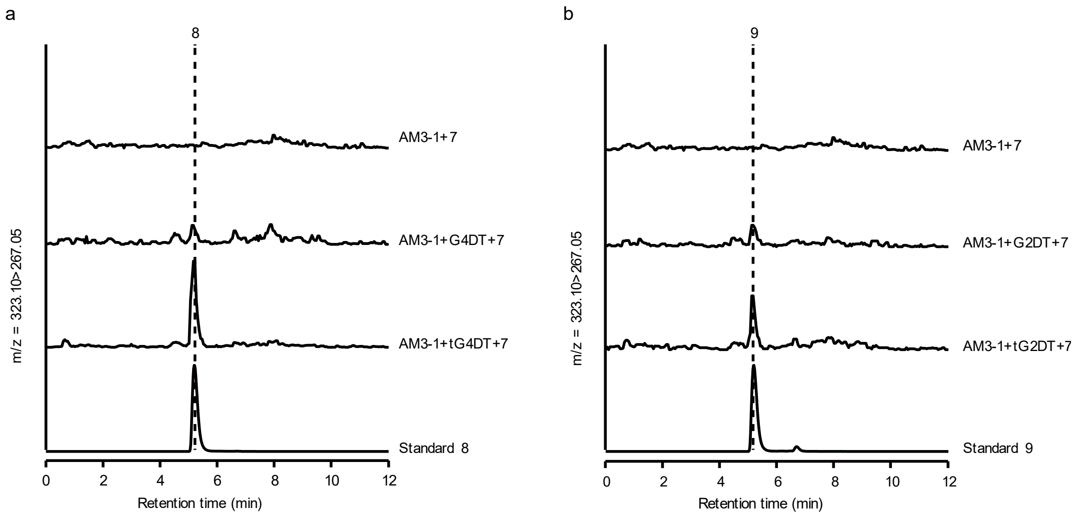

**Supplementary Fig. 18** Feeding experiment in strain AM3-1 where the native ERG20p was replaced with a dynamically regulated promoter HXT1p, demonstrating the conversion of **7** to **8** (a) and **9** (b).

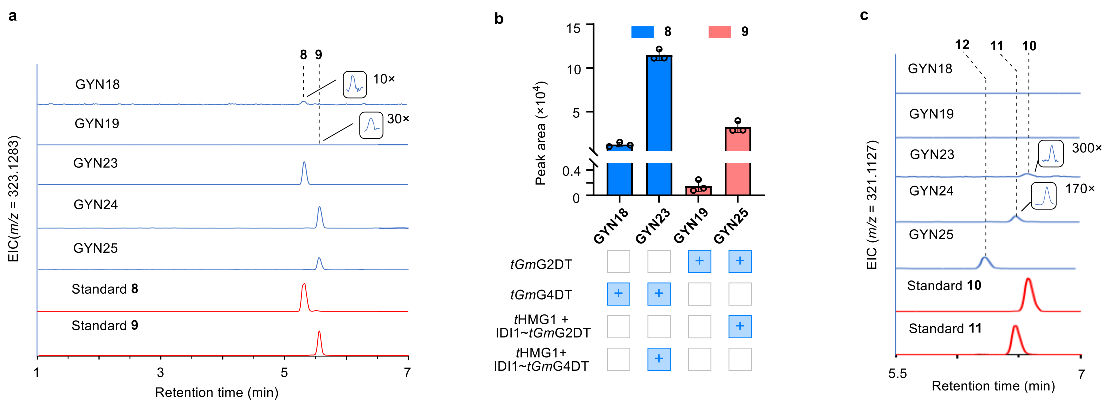

**Supplementary Fig. 19** Reconstitution of the glyceollin module for de novo biosynthesis of glyceollins. (a) Fusion expression of IDI1 with tGmG2DT/tGmG4DT, enhancing the titer of 8 and 9. EICs for 8 and 9 ([M-H2O+ H]+ = m/z 323.1283) are presented. (b) Changes in the titre of 8 and 9 in strains GYN18, GYN23, GYN19, and GYN25. (c) LC-MS profiles of glyceollins I, II, and III-producing strains. EICs for 10, 11, and 12 ([M-H2O+ H]+ = m/z 321.1127) are presented. Data are shown as mean value ± standard deviations (n = 3 biologically independent samples).

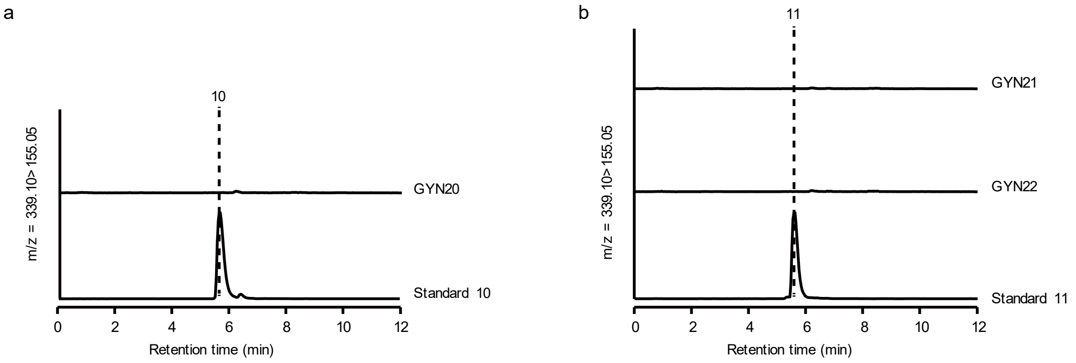

**Supplementary Fig. 20** LC-MS analysis of the fermented products from strains GYN20 to GYN22. The MRM spectra for the selected ion pair of *m/z* 339.10 > 155.05 are presented for the standards of glyceollin I (**10**) and glyceollin II (**11**), as well as for the metabolites from the three strains. Strains GYN20 to GYN22 express *Gm*GIS2, *Gm*GIIS, and *Gm*GIIIS individually.

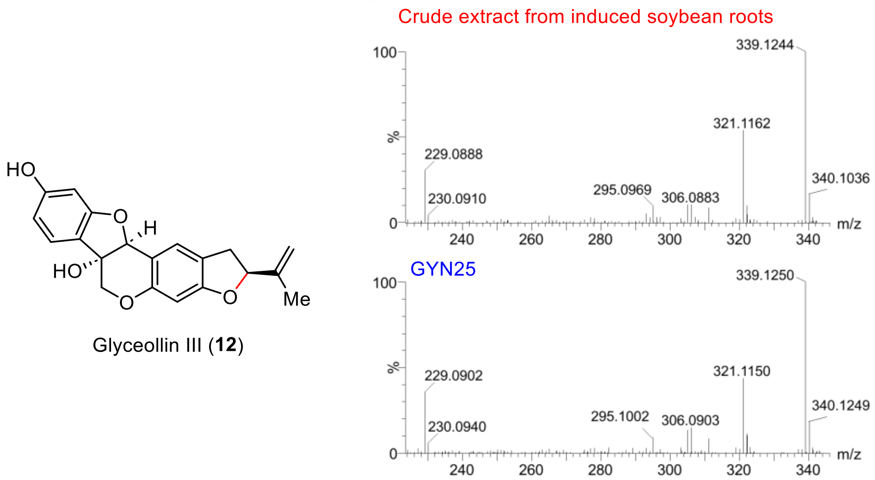

**Supplementary Fig. 21** MS/MS (20 to 50 eV) spectra of generated glyceollin III (**12**) from yeast strain GYN25 compared to standard **12** from crude extract from induced soybean roots.

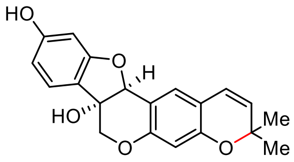

Glyceollin II (**11**)

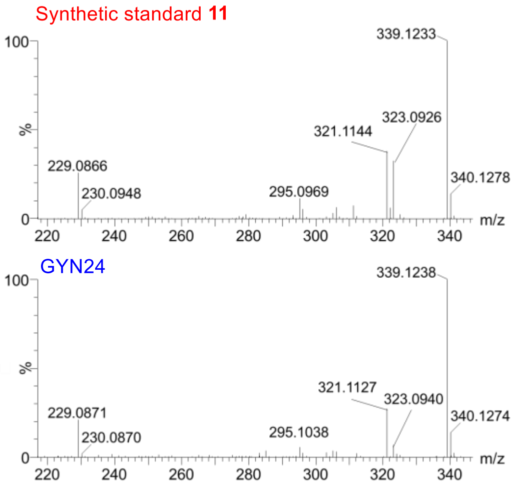

**Supplementary Fig. 22** MS/MS (20 to 50 eV) spectra of generated glyceollin II (**11**) from yeast strain GYN24 compared to synthetic standard **11**.

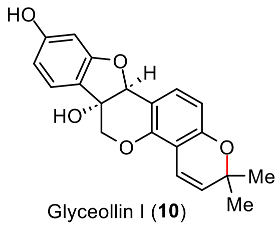

Glyceollin I (**10**)

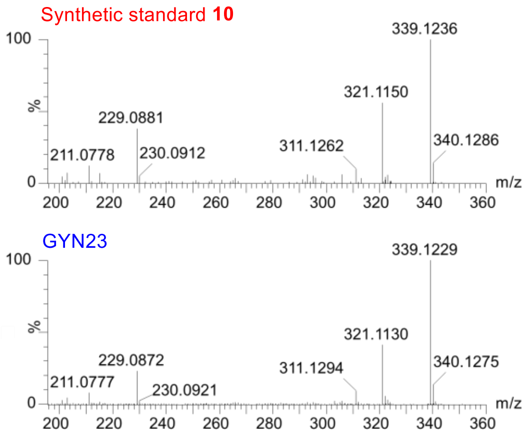

**Supplementary Fig. 23** MS/MS (20 to 50 eV) spectra of generated glyceollin I (**10**) from yeast strain GYN23 compared to synthetic standard **10**.

**Supplementary Table 1.** Candidate genes encoding THIS from *Glycine max*

| **Gene** | **% Identity (*Ms*VR)** | **% Identity (*Ps*SR)** |
|---|---|---|
| Glyma.18G220500 (***Gm*THIS1**) | 79.2 | 74.4 |
| Glyma.09G269600 (***Gm*THIS2**) | 78.9 | 75.0 |
| Glyma.18G220600 | 77.4 | 73.8 |
| Glyma.09G269500 | 76.5 | 72.9 |
| Glyma.09G269400 | 69.9 | 69.1 |
| Glyma.13G203900 | 64.5 | 64.6 |
| Glyma.12G238100 | 63.6 | 64.3 |

**Supplementary Table 2**. Expression levels of genes in the glyceollin biosynthetic pathway after treatment with WGE and water

| Name | Gene | Expression level (FPKM) | | | | | | | |
|---|---|---|---|---|---|---|---|---|---|
| | | WGE1 | WGE2 | WGE3 | WGE4 | Water1 | Water2 | Water3 | Water4 |
| *Gm*PAL | Glyma.10G058200 | 347.0 | 514.9 | 533.9 | 392.0 | 298.6 | 247.6 | 335.3 | 247.2 |
| *Gm*C4H | Glyma.02G236500 | 194.1 | 229.3 | 195.6 | 184.7 | 208.2 | 203.0 | 218.8 | 200.9 |
| *Gm*4CL1 | Glyma.17G064600 | 57.2 | 73.0 | 68.8 | 53.6 | 57.4 | 57.3 | 113.2 | 74.6 |
| *Gm*CHR5 | Glyma.18G285800 | 874.0 | 1160.2 | 990.4 | 801.4 | 698.7 | 710.9 | 557.3 | 702.1 |
| *Gm*CHI1B2 | Glyma.10G292200 | 72.7 | 102.2 | 86.7 | 65.8 | 46.3 | 56.0 | 56.6 | 36.3 |
| *Gm*2-HIS | Glyma.13G173500 | 390.2 | 628.0 | 553.3 | 415.5 | 168.3 | 164.9 | 154.4 | 150.4 |
| *Gm*HID | Glyma.01G239600 | 341.1 | 468.0 | 375.0 | 313.1 | 248.3 | 263.4 | 287.9 | 242.0 |
| *Gm*CYP81E | Glyma.09G049100 | 352.9 | 456.5 | 346.7 | 329.4 | 305.1 | 231.4 | 377.7 | 261.6 |
| *Gm*IFR | Glyma.11G070500 | 48.3 | 73.5 | 56.3 | 49.6 | 17.7 | 16.2 | 17.3 | 15.2 |
| *Gm*THIS1 | Glyma.18G220500 | 102.5 | 129.3 | 92.0 | 94.8 | 88.3 | 68.6 | 90.7 | 83.3 |
| *Gm*THIS2 | Glyma.09G269600 | 74.4 | 131.1 | 91.0 | 70.5 | 23.8 | 26.2 | 37.9 | 30.5 |
| | Glyma.18G220600 | 78.0 | 99.5 | 70.7 | 80.6 | 53.0 | 57.1 | 54.3 | 56.2 |
| | Glyma.09G269500 | 35.2 | 47.2 | 38.1 | 38.2 | 18.8 | 18.8 | 24.9 | 19.6 |
| reductase | Glyma.09G269400 | 1.6 | 12.8 | 19.9 | 25.2 | 0.4 | 0.0 | 0.2 | 0.1 |
| | Glyma.13G203900 | 0.0 | 0.0 | 0.0 | 0.0 | 0.0 | 0.0 | 0.0 | 0.0 |
| | Glyma.12G238100 | 0.0 | 0.0 | 0.0 | 0.0 | 0.0 | 0.0 | 0.0 | 0.0 |
| *Gm*PTS | Glyma.03G147700 | 214.4 | 388.3 | 248.8 | 208.9 | 92.5 | 59.3 | 122.4 | 89.9 |
| *Gm*CYP93A | Glyma.03G143700 | 363.2 | 450.6 | 396.1 | 318.9 | 153.2 | 164.6 | 183.8 | 179.4 |
| *Gm*G4DT | Glyma.10G295300 | 153.1 | 160.1 | 163.5 | 171.5 | 29.9 | 30.2 | 23.8 | 33.9 |
| *Gm*G2DT | Glyma.20G245100 | 21.1 | 32.9 | 27.9 | 20.2 | 18.0 | 13.3 | 25.3 | 20.9 |
| *Gm*GIIS | Glyma.01G135200 | 85.1 | 141.0 | 133.8 | 66.7 | 1.3 | 1.5 | 2.5 | 1.0 |
| *Gm*GIS1 | Glyma.11G062500 | 42.8 | 93.8 | 66.2 | 31.6 | 3.5 | 2.4 | 4.8 | 2.6 |
| *Gm*GIS2 | Glyma.11G062600 | 170.6 | 331.7 | 311.9 | 205.3 | 32.2 | 24.9 | 33.4 | 21.4 |
| *Gm*GIIIS | Glyma.13G285300 | 47.8 | 96.5 | 67.4 | 38.8 | 3.8 | 2.8 | 5.0 | 3.5 |
| *Gm*GIIISp | Glyma.15G203500 | 0.7 | 6.8 | 2.0 | 0.8 | 1.4 | 0.3 | 0.1 | 0.0 |
| *Gm*GIS3 | Glyma.01G179700 | 0.1 | 0.1 | 0.5 | 0.1 | 0.0 | 0.0 | 0.0 | 0.0 |

**Supplementary Table 3**. Upregulated CYP genes with expression levels at least twice as high as those in the control group

| **Transcript ID** | **P450 Family** | **Fold Change (WGE treatment over water_24h)** | **Up or Down regulation** |
|---|---|---|---|
| **Glyma.01G135200 (*Gm*GIIS)** | CYP82 | 67.7 | Up |
| Glyma.09G049300 | CYP81 | 19.6 | Up |
| Glyma.16G195600 | CYP71 | 19.2 | Up |
| Glyma.09G048900 | CYP81 | 17.3 | Up |
| **Glyma.11G062500 (*Gm*GIS1)** | CYP71 | 17.6 | Up |
| Glyma.09G049200 | CYP81 | 14.7 | Up |
| **Glyma.13G285300 (*Gm*GIIIS)** | CYP82 | 16.6 | Up |
| Glyma.13G265300 | CYP76 | 12.3 | Up |
| Glyma.18G080400 | CYP71 | 10.9 | Up |
| Glyma.11G051800 | CYP81 | 7.5 | Up |
| **Glyma.11G062600 (*Gm*GIS2)** | CYP71 | 9.1 | Up |
| Glyma.05G022100 | CYP75 | 7.4 | Up |
| Glyma.05G042800 | CYP71 | 7.0 | Up |
| Glyma.15G156100 | CYP81 | 6.3 | Up |
| Glyma.20G188000 | CYP77 | 5.4 | Up |
| Glyma.19G146800 | CYP93 | 5.1 | Up |
| Glyma.13G068800 | CYP82 | 4.0 | Up |
| **Glyma.15G203500 (*Gm*GIIISp)** | CYP82 | 5.7 | Up |
| Glyma.08G140600 | CYP736 | 3.4 | Up |
| Glyma.08G104100 | CYP78 | 3.4 | Up |
| Glyma.10G115500 | CYP71 | 2.9 | Up |
| Glyma.13G173500 | CYP93 | 2.8 | Up |
| Glyma.10G203500 | CYP76 | 2.7 | Up |
| Glyma.05G010200 | CYP734 | 2.5 | Up |
| Glyma.14G015100 | CYP71 | 2.3 | Up |
| Glyma.03G143700 | CYP93 | 2.2 | Up |
| Glyma.03G160100 | CYP94 | 2.2 | Up |
| Glyma.20G114200 | CYP73 | 2.0 | Up |

**Supplementary Table 4.** List of yeast strains constructed in this study

| Name | Genotype | Reference |
|---|---|---|
| AM3-1 | CEN.PK2-Δ*ERG20p*::*HXT1p* | (Liu et al., 2022) |
| CenA1 | CEN.PK2-*CrCYB5-AtCPR1* | (Liu et al., 2021) |
| CenA1ΔA10 | CenA1-Δ*ARO10*::*ARO4-ARO7* | This study |
| GYN1 | CenA1ΔA10-IntL7::*FjTAL-Pc4CL*; IntL8::*PhCHS~GmCHR5-PhCHI*; IntL9::Gm*CHR5* | This study |
| GYN2 | GYN1-IntZ6::op*GmCHIB2* | This study |
| GYN3 | GYN2-IntZ15::op*Ge2-HIS*-op*GmHID*; IntZ6::op*GmCHIB2* | This study |
| GYN4 | GYN2-IntZ15::op*AmIFS*-op*GmHID*; IntZ6::op*GmCHIB2* | This study |
| GYN5 | GYN3-IntZ14::*GmCYP81E* | This study |
| GYN6 | GYN3-IntZ14::*GmCYP81E-GmIFR* | This study |
| GYN7 | GYN3-IntZ14::*GmCYP81E-GmIFR*; IntZ13::*GmTHIS1* | This study |
| GYN8 | GYN3-IntZ14::*GmCYP81E-GmIFR*; IntZ13::*GmTHIS1-GmPTS* | This study |
| GYN9 | GYN8-IntZ10::*GmCYP93A* | This study |
| GYN10 | GYN8-IntZ10::*GmCYP93A-GmG4DT* | This study |
| GYN11 | GYN8-IntZ10::*GmCYP93A-GmG2DT* | This study |
| GYN12 | GYN8-IntZ10::*GmCYP93A-tGmG4DT* | This study |
| GYN13 | GYN8-IntZ10::*GmCYP93A-tGmG2DT* | This study |
| GYN14 | GYN10-IntZ11:: *GmGIS1* | This study |
| GYN15 | GYN10-IntZ11::*GmGIS2* | This study |
| GYN16 | GYN11-IntZ11::*GmGIIS* | This study |
| GYN17 | GYN11-IntZ11::*GmGIIIS* | This study |
| GYN18 | GYN12-Δ*ERG20p*::*HXT1p* | This study |
| GYN19 | GYN13-Δ*ERG20p*::*HXT1p* | This study |
| GYN20 | GYN18-IntZ11::*GmGIS2* | This study |
| GYN21 | GYN19-IntZ11::*GmGIIS* | This study |
| GYN22 | GYN19-IntZ11::*GmGIIIS* | This study |
| GYN23 | GYN20-IntZ4::*SctHMG1-ScIDI1~tGmG4DT* | This study |
| GYN24 | GYN21-IntZ4::*SctHMG1-ScIDI1~tGmG2DT* | This study |
| GYN25 | GYN22-IntZ4::*SctHMG1-ScIDI1~tGmG2DT* | This study |
| GYN231 | GYN23-IntZ29::*GmGIS1** | This study |
| GYN232 | GYN23-IntZ9::*GmGIS2* | This study |
| GYN233 | GYN23-IntZ9::*GmGIS2** | This study |
| GYN234 | GYN233-IntZ29::*GmGIS2** | This study |
| GYN242 | GYN24-IntZ9::*GmGIIS* | This study |
| GYN243 | GYN242-IntZ35::*GmGIIS* | This study |
| GYN252 | GYN25-IntZ9::*GmGIIIS* | This study |

| | | |
|---|---|---|
| GYN253 | GYN252-IntZ35::*GmGIIIS* | This study |

Note: "-" indicates that two gene expression cassettes locate in the same integration site. "~" indicates gene fusion. *GmGIS1**, *GmGIS2**, and *GmGIS2** represent that these genes' N-terminal signal peptide are replaced with that of *GmGIIIS*.

**Supplementary Table 5.** Primers used in this study

| Name | Sequence (5'-3') |
|---|---|
| pOPINF-THIS1-F | AAGTTCTGTTTCAGGGTACCATGGCAGAGGGAAAAGGTAG |
| pOPINF-THIS1-R | CAAACTGGTCTAGAAAGCTTTTAAAGATAACCCTTTTCCT |
| pOPINF-THIS2-F | AAGTTCTGTTTCAGGGTACCATGGCAGAGGGAAAAGGTAG |
| pOPINF-THIS2-R | CAAACTGGTCTAGAAAGCTTTTAAAGATAACCCTTTTCCT |
| pOPINF-seqF | CTCTAGAGCCTCTGCTAACCATGTT |
| pOPINF-seqR | TTATTAGCCAGAAGTCAGATGCTCA |
| pOPINF-MsVR-F | AAGTTCTGTTTCAGGGTACCATGGCTGAAGGAAAAGGAAGA |
| pOPINF-MsVR-R | CAAACTGGTCTAGAAAGCTTTTAGAGATAGCCTTTCTCCTT |
| pOPINF-PsSR-F | AAGTTCTGTTTCAGGGTACCATGGCAGAGGGGAAAGGAA |
| pOPINF-PsSR-R | AAGTTCTGTTTCAGGGTACCTTAGAGATATCCTTTTTCCTTGC |
| 3Ω-MsVR-F | TTTATGAATTTTGCAGCTCGATGGCTGAAGGAAAAGGAAGA |
| 3Ω-MsVR-R | GACAACCACAACAAGCACCGTTAGAGATAGCCTTTCTCCTT |
| 3Ω-PsSR-F | TTTATGAATTTTGCAGCTCGATGGCAGAGGGGAAAGGAA |
| 3Ω-PsSR-R | GACAACCACAACAAGCACCGTTAGAGATATCCTTTTTCCTTGC |
| 3Ω-GmGIS1-F | TTTATGAATTTTGCAGCTCGATGGAACATTCTCAACTGTCC |
| 3Ω-GmGIS1-R | GACAACCACAACAAGCACCGTCATGTAGCTTGGTAAACGGT |
| 3Ω-GmGIS2-F | TTTATGAATTTTGCAGCTCGATGGAATATTCTCCATTGTCCA |
| 3Ω-GmGIS2-R | GACAACCACAACAAGCACCGTCATGAAGCTTCATAAACAGTG |
| 3Ω-GmGIS3-F | TTTATGAATTTTGCAGCTCGATGGAATATTCTCCACTGTCC |
| 3Ω-GmGIS3-R | GACAACCACAACAAGCACCGTCATGAAGCTTCATAAATAGTGG |
| 3Ω-GmGIIS-F | TTTATGAATTTTGCAGCTCGATGGAATTAGTTCTACATTTCCTAA |
| 3Ω-GmGIIS-R | GACAACCACAACAAGCACCGTCACATACTTTTGTAACAACTTGG |
| 3Ω-GmGIIIS-F | TTTATGAATTTTGCAGCTCGATGGAGTTAGTTCTAAACAGCA |
| 3Ω-GmGIIIS-R | GACAACCACAACAAGCACCGTTAGATACTTTCATAACAACTAGG |
| 3Ω-GmGIIISp-F | TTTATGAATTTTGCAGCTCGATGGACTTAGTTCTAAACACC |
| 3Ω-GmGIIISp-R | GACAACCACAACAAGCACCGTTACATACTTTCATAACAACTAGGAG |
| 3Ω-GmTHIS1-F | TTTATGAATTTTGCAGCTCGATGGCAGAGGGAAAAGGTAG |
| 3Ω-GmTHIS1-R | GACAACCACAACAAGCACCGTTAAAGATAACCCTTTTCCT |
| 3Ω-GmTHIS2-F | TTTATGAATTTTGCAGCTCGATGGCAGAGGGAAAAGGTAG |
| 3Ω-GmTHIS2-R | GACAACCACAACAAGCACCGTTAAAGATAACCCTTTTCCT |
| 3Ω-seq-F | TCGGTTTAATGAGAAGGCCTAAAAT |
| 3Ω-seq-R | CTGTAAGTGGTATAGTAAACGGAAG |
| pESC- GmGIS1-F | GAGAAAAAACCCCGGATCCATGATGGAACATTCTCAACTGTCC |
| pESC- GmGIS1-R | ACTTCTGTTCCATGTCGACTCATGTAGCTTGGTAAACGGT |
| pESC- GmGIS2-F | GAGAAAAAACCCCGGATCCATGATGGAATATTCTCCATTGTCCA |
| pESC- GmGIS2-R | ACTTCTGTTCCATGTCGACTCATGAAGCTTCATAAACAGTG |
| pESC- GmGIS3-F | GAGAAAAAACCCCGGATCCATGATGGAATATTCTCCACTGTCC |
| pESC- GmGIS3-R | ACTTCTGTTCCATGTCGACTCATGAAGCTTCATAAATAGTGG |
| pESC- GmGIIS-F | GAGAAAAAACCCCGGATCCATGATGGAATTAGTTCTACATTTCCTAA |
| pESC- GmGIIS-R | ACTTCTGTTCCATGTCGACTCACATACTTTTGTAACAACTTGG |
| pESC- GmGIIIS-F | GAGAAAAAACCCCGGATCCATGATGGAGTTAGTTCTAAACAGCA |
| pESC- GmGIIIS-R | ACTTCTGTTCCATGTCGACTTAGATACTTTCATAACAACTAGG |

**Supplementary Table 5**. Primers used in this study (continued)

| | |
|---|---|
| GmGIIISp_gv-ra-F | AGAGCAACCAACTCTAAAGCCACTTC |
| GmGIIISp_gv-ra-R | TTAGAGTTGGTTGCTCTAAAGACTTCAGTCATATCAAGAG |
| GmGIIISp_ l-v-F | GTCGTAGCACCCTACGGTCCCTAT |
| GmGIIISp _l-v-R | CGTAGGGTGCTACGACTATCATGGATCGGTTGTAGC |
| GmGIIISp_ g-a-F | GCTCCTCTCTCAAGACCTCGT |
| GmGIIISp _g-a-R | GTCTTGAGAGAGGAGCTGGAGGGTACAATCTTAAAG |
| GmGIS3_p-s-F | TCAGCATCAACATTGGAGTGGG |
| GmGIS3_p-s-R | CAATGTTGATGCTGAGGTATCAGTTCCAGAAGCAA |
| GmGIS3_v-a-F | GCTTTTGCTCCATACGGTGATTA |
| GmGIS3_v-a-R | CGTATGGAGCAAAAGCAATATCTGTTGCTCCATATACC |
| GmGIS3_s-a-F | GCTGGAACTGATACCCCGGCATC |
| GmGIS3_s-a-R | GGTATCAGTTCCAGCAGCAAATATGTTCCATATCAC |
| ScARO4 (K229L)-F | ATGAGTGAATCTCCAATGTTCG |
| ScARO4 (K229L)-R | CTATTTCTTGTTAACTTCTCTTCTTTGTC |
| ScARO7 (G141S)-F | ATGGATTTCACAAAACCAGAA |
| ScARO7 (G141S)-R | TTACTCTTCCAACCTTCTTAG |
| FjTAL-F | ATGAACACCATTAACGAATAC |
| FjTAL-R | TTAGTTGTTAATTAAATGATC |
| Pc4CL2-F | ATGGGTGATTGTGTTGCTCCA |
| Pc4CL2-R | TTATTTTGGCAAATCACCAGA |
| PhCHS-F | ATGGTTACCGTTGAAGAATAT |
| PhCHS-R | TTAAGTAGCAACACTATGCAA |
| GmCHR5-F | ATGGCTGCCACCACCTTAGTC |
| GmCHR5-R | TCATTCTTCATCCCATAGATC |
| PhCHI-F | ATGTCACCACCAGTTTCTGTT |
| PhCHI-R | TAAACACCAATAACTGGAATT |
| GmCHIB2-F | ATGGCCACCCCTGCTTCTATT |
| GmCHIB2-R | TCAATTTTCAATGTTTGGGTT |
| Ge2-HIS-F | ATGTTGGTCGAACTGGCGATT |
| Ge2-HIS-R | TTATGAGGAGAATAACTTTGG |
| GmHID-F | ATGGCCAAGGAAATTGTTAAG |
| GmHID-R | TTAGACCAAGAAAGAAGCTAA |
| AmIFS-F | ATGTTGTTAGAACTTGCAGTA |
| AmIFS-R | TCAAGAGGAAAGGAGTTTAGC |
| GmCYP81E-F | GTCAAGGAGAAAAAACCCCGGATCCATGACAGTAATAACAATGCCT |
| GmCYP81E-R | AGCCGCGGTACCAAGCTTACTCGAGTTAATAACTTCCAACTTTGCT |
| GmIFR-F | TCGAATTCAACCCTCACTAAAGGGCATGGCTGGGAAAGATAGAATC |
| GmIFR-R | CCTTGTAATCCATCGATACTAGTGCTCAGACAAACTGATTCAAATA |
| GmTHIS1-F | GTCAAGGAGAAAAAACCCCGGATCCATGGCAGAGGGAAAAGGTAGA |
| GmTHIS1-R | AGCCGCGGTACCAAGCTTACTCGAGTTAAAGATAACCCTTTTCCTT |
| GmPTS-F | TCGAATTCAACCCTCACTAAAGGGCATGGCCAAATCCACTTTCTTT |
| GmPTS-R | CCTTGTAATCCATCGATACTAGTGCTCAATAATGGTTCACGAATAC |
| GmCYP93A-F | GTCAAGGAGAAAAAACCCCGGATCCATGGCTTATCAAGTGTTGCTA |

**Supplementary Table 5**. Primers used in this study (continued)

| | |
|---|---|
| GmCYP93A-R | AGCCGCGGTACCAAGCTTACTCGAGTCAAATAGTAGGGAATGGGTT |
| GmG4DT-F | TCGAATTCAACCCTCACTAAAGGGCATGGATTGGGGGCTTGCTATA |
| GmG4DT-R | CCTTGTAATCCATCGATACTAGTGCTCATCTAATTAATGCCATGAG |
| GmG2DT-F | TCGAATTCAACCCTCACTAAAGGGCATGGATTCGGGGTCTGTTATA |
| GmG2DT-R | CCTTGTAATCCATCGATACTAGTGCTTATCTAATTAAAGCCATGAG |
| GmGIS1-F | TCGAATTCAACCCTCACTAAAGGGCATGGAACATTCTCAACTGTCC |
| GmGIS1-R | CCTTGTAATCCATCGATACTAGTGCTCATGTAGCTTGGTAAACGGT |
| GmGIS2-F | TCGAATTCAACCCTCACTAAAGGGCATGGAATATTCTCCATTGTCC |
| GmGIS2-R | CCTTGTAATCCATCGATACTAGTGCTCATGAAGCTTCATAAACAGT |
| GmGIIS-F | TCGAATTCAACCCTCACTAAAGGGCATGGAATTAGTTCTACATTTC |
| GmGIIS -R | CCTTGTAATCCATCGATACTAGTGCTCACATACTTTTGTAACAACT |
| GmGIIIS -F | TCGAATTCAACCCTCACTAAAGGGCATGGAGTTAGTTCTAAACAGC |
| GmGIIIS -R | CCTTGTAATCCATCGATACTAGTGCTTAGATACTTTCATAACAACT |
| tGmG4DT-F | TCGAATTCAACCCTCACTAAAGGGCATGCACAAAAGGGAAACTCAA |
| tGmG2DT-F | TCGAATTCAACCCTCACTAAAGGGCATGTACAAAAAGAAAACTCAA |
| tScHMG1-F | GTCAAGGAGAAAAAACCCCGGATCCATGGCTGCAGACCAATTGGTG |
| tScHMG1-R | AGCCGCGGTACCAAGCTTACTCGAGTTAGGATTTAATGCAGGTGAC |
| ScIDI1-F | TCGAATTCAACCCTCACTAAAGGGCATGACTGCCGACAACAATAGT |
| ScIDI1-R-linker | ACCTCCGCCACCACTACCACCACCGCCTAGCATTCTATGAATTTG |
| Linker-tGmG4DT-F | GGTAGTGGTGGCGGAGGTTCTGGTGGAGGAGGTAGCCACAAAAGGGAAACTCAA |
| Linker-tGmG2DT-F | GGTAGTGGTGGCGGAGGTTCTGGTGGAGGAGGTAGCTACAAAAAGAAAACTCAA |

Note: Sequence with red marker indicates homology arm.

**Supplementary Table 6**. List of genes used for the biosynthesis of glyceollins

| Name | Annotation | Source | Accession in NCBI |
|---|---|---|---|
| *At*CPR1 | P450 reductase 1 | *Arabidopsis thaliana* | NP_001190823 |
| *Cr*CYB5 | Cytochrome b5 | *Catharanthus roseus* | AJO70762 |
| *Sc*ARO4 | 3-deoxy-7-phosphoheptulonate synthase | *Saccharomyces cerevisiae* | NP_009808 |
| *Sc*ARO7 | Chorismate mutase | *Saccharomyces cerevisiae* | NP_015385 |
| *Fj*TAL | Histidine ammonia-lyase | *Flavobacterium johnsoniae* | WP_012023194 |
| *Pc*4CL2 | 4-coumaroyl-CoA synthase 2 | *Petroselinum crispum* | P14913 |
| *Ph*CHS | Chalcone synthase | *Petunia x hybrida* | AAF60297 |
| *Gm*CHR5 | Chalcone reductase CHR5 | *Glycine max* | NP_001353935 |
| *Ph*CHI | Chalcone flavanone isomerase | *Petunia x hybrida* | 1807331A |
| *Gm*CHIB2 | Chalcone--flavanone isomerase 1B-2 | *Glycine max* | NP_001236097 |
| *Ge*2-HIS | 2-hydroxyisoflavanone synthase | *Glycyrrhiza echinata* | Q9SXS3 |
| *Gm*HID | 2-hydroxyisoflavanone dehydratase | *Glycine max* | NP_001237228 |
| *Am*IFS | Isoflavone synthase | *Astragalus membranaceus* | AEH68209 |
| *tSc*HMG1 | Hydroxymethylglutaryl-CoA reductase | *Saccharomyces cerevisiae* | KAJ1536229 |
| *Sc*IDI1 | Isopentenyl-diphosphate delta-isomerase | *Saccharomyces cerevisiae* | NP_015208 |
| *Sc*EMC1 | ER membrane protein complex subunit 1 | *Saccharomyces cerevisiae* | NP_009884 |

**Supplementary Table 7**. List of integration sites used in this study

| Name | Spacer sequences (5'-3') | Chromosomal locus |
|---|---|---|
| IntL7 | AATCCGAACAACAGAGCATA | ChrXVI: 776,883-776,902 |
| IntL8 | GGTTTTCATACTGGGGCCGC | ChrVI: 237073-237092 |
| IntL9 | GCGCCACAGTTTCAAGGGTC | ChrXIV: 280,250-280,269 |
| IntZ4 | CCTGGCGCTATGATGATGAG | ChrVII: 478,898-478,917 |
| IntZ6 | AAGATAGTCGACCCTTACAC | ChrVII: 318707-318688 |
| IntZ9 | GAGATGGTTCTATCGGACCA | ChrVII: 609985-609966 |
| IntZ10 | CACGCAATCCAACAAATCGG | ChrVII: 678274-678255 |
| IntZ11 | AATCTGGAGATAAGGCAACG | ChrVII: 857174-857155 |
| IntZ13 | GAGCATTTACTGACACCTGG | ChrVII: 1011327-1011308 |
| IntZ14 | AATGGATAAAAAATACAACG | ChrVIII: 90487-90506 |
| IntZ15 | GAGGACAGCGTGAATCACAA | ChrVIII: 116848-116867 |
| IntZ9 | GAGATGGTTCTATCGGACCA | ChrVII: 609985-609966 |
| IntZ29 | GAGTCACGCTGAACACGCGG | ChrXV: 1019467-1019486 |
| IntZ35 | ATTACCCCAGAGCTGCTACG | ChrXVI: 114581-114562 |

**Supplementary Table 8**. List of plasmids constructed in this study

| Plasmid | Description |
|---|---|
| pOPINF-THIS1 | pOPINF-Amp |
| pOPINF-THIS2 | pOPINF- Amp |
| pESC- GmGIS1 | pESC-Kana |
| pESC- GmGIS2 | pESC- Kana |
| pESC- GmGIIS | pESC- Kana |
| pESC- GmGIIIS | pESC- Kana |
| 3Ω-GmTHIS1 | 3Ω-Ccdb-Spe |
| 3Ω-GmTHIS2 | 3Ω-Ccdb-Spe |
| 3Ω-GmGIS1 | 3Ω-Ccdb-Spe |
| 3Ω-GmGIS2 | 3Ω-Ccdb-Spe |
| 3Ω-GmGIS3 | 3Ω-Ccdb-Spe |
| 3Ω-GmGIIS | 3Ω-Ccdb-Spe |
| 3Ω-GmGIIIS | 3Ω-Ccdb-Spe |
| pESC-HIS-ScARO4-ARO7 | 2μ; HIS3; AmpR; GAL1p-ScARO4-CYC1t; GAL10p-ARO7-ADH1t |
| pESC-HIS-FjTAL-Pc4CL2 | 2μ; HIS3; AmpR; GAL1p-FjTAL-CYC1t; GAL10p-Pc4CL2-ADH1t |
| pESC-LEU2d-PhCHS-GmCHR5 | 2μ; LEU2d; AmpR; GAL1p-PhCHS-CYC1t; GAL10p-GmCHR5-ADH1t |
| pESC-LEU2d-PhCHI | 2μ; LEU2d; AmpR; GAL10p-PhCHI-ADH1t |
| pESC-HIS-opGmCHIB2 | 2μ; HIS3; AmpR; GAL1p-opGmCHIB2-CYC1t |
| pESC-HIS-opGmHID | 2μ; HIS3; AmpR; GAL10p-opGmHID-CYC1t |
| pESC-HIS-opGe2-HIS-opGmHID | 2μ; HIS3; AmpR; GAL1p-opGe2-HIS-CYC1t; GAL10p-opGmHID-ADH1t |
| pESC-HIS-opAmIFS-opGmHID | 2μ; HIS3; AmpR; GAL1p-opAmIFS-CYC1t; GAL10p-opGmHID-ADH1t |
| pESC-HIS-GmCYP81E | 2μ; HIS3; AmpR; GAL1p-GmCYP81E-CYC1t |
| pESC-HIS-GmCYP81E-GmIFR | 2μ; HIS3; AmpR; GAL1p-GmCYP81E-CYC1t; GAL10p-GmIFR-ADH1t |
| pESC-HIS-GmTHIS1 | 2μ; HIS3; AmpR; GAL1p-GmTHIS1-CYC1t |
| pESC-HIS-GmTHIS1-GmPTS | 2μ; HIS3; AmpR; GAL1p-GmTHIS1-CYC1t; GAL10p-GmPTS-ADH1t |
| pESC-HIS-GmCYP93A | 2μ; HIS3; AmpR; GAL1p-GmCYP93A-CYC1t |
| pESC-HIS-GmCYP93A-GmG2DT | 2μ; HIS3; AmpR; GAL1p-GmCYP93A-CYC1t; GAL10p-GmG2DT-ADH1t |
| pESC-HIS-GmCYP93A-GmG4DT | 2μ; HIS3; AmpR; GAL1p-GmCYP93A-CYC1t; GAL10p-GmG4DT-ADH1t |
| pESC-HIS-GmCYP93A-tGmG2DT | 2μ; HIS3; AmpR; GAL1p-GmCYP93A-CYC1t; GAL10p-tGmG2DT-ADH1t |
| pESC-HIS-GmCYP93A-tGmG4DT | 2μ; HIS3; AmpR; GAL1p-GmCYP93A-CYC1t; GAL10p-tGmG4DT-ADH1t |
| pESC-HIS-SctHMG1 | 2μ; HIS3; AmpR; GAL1p-tSctHMG1-CYC1t |
| pESC-HIS-SctHMG1-IDI1-tGmG2DT | 2μ; HIS3; AmpR; GAL1p-tSctHMG1-CYC1t; GAL10p-IDI1-tGmG2DT-ADH1t |
| pESC-HIS-SctHMG1-IDI1-tGmG4DT | 2μ; HIS3; AmpR; GAL1p-tSctHMG1-CYC1t; GAL10p-IDI1-tGmG4DT-ADH1t |
| pESC-HIS- GmGIIS | 2μ; HIS3; AmpR; GAL1p- GmGIIS -CYC1t |
| pESC-HIS- GmGIS1 | 2μ; HIS3; AmpR; GAL1p- GmGIS1-CYC1t |
| pESC-HIS- GmGIIIS | 2μ; HIS3; AmpR; GAL1p- GmGIIIS -CYC1t |
| pESC-HIS- GmGIS2 | 2μ; HIS3; AmpR; GAL1p- GmGIS2-CYC1t |

**Supplementary Table 9.** Codon-optimized genes for yeast expression used in this study

| Gene | Description |
|---|---|
| opGmCHIB2 | ATGGCCACCCCTGCTTCTATTACCAACGTCACTGTCGAATTCTTGCAATTCCCAGCTTTGGTTACTCCAC<br>CTGCTTCCACCAAGTCATACTTTTTAGGTGGTGCCGGTGTCCGTGGTTTGAACATCCAAGAAGAATTT<br>GTTAAGTTCACAGGTATCGGTGTTTACTTGGAAGACAAAGCTGTTTCCTCCCTTGGTGCAAAATGGAA<br>GGGCAAGTCTGCTGCTGAATTGTTGGACTCTTTGGACTTTTATAGAGATATTATCAAGGGTCCATTCGA<br>AAAGTTGATCAGAGGTTCCAAGTTGAGAACTCTAGATGGTAGAGAATACGTTAGAAAGGTTAGCGAA<br>AACTGTGTTGCTCACATGGAGTCCGTCGGTACTTACTCTGAAGCCGAAGAAAAAGCTATCGAAGAATT<br>CAGAAACGCTTTCAAGGACCAAAACTTCCCACCAGGTTCTACTGTTTTCTACAAGCAATCTCCAACCG<br>GTACTTTGGGTTTATCTTTCTCTAAGGATGAAACTATTCCAGAACACGAACACGCTGTCATTGACAACA<br>AGCCATTGAGTGAAGCCGTCTTGGAAACCATGATTGGTGAAATCCCAGTTTCCCCAGCTTTGAAGGA<br>ATCTTTGGCTACCAGATTCCATCAATTCTTCAAGGAATTAGAAGCTAACCCAAACATTGAAAATTGA |
| opGe2-HIS | ATGTTGGTCGAACTGGCGATTACCTTGTTGGTTATTGCTTTGTTCATCCATTTGCGTCCAACTCTATCTG<br>CCAAGTCCAAGTCTCTTAGACACTTGCCAAATCCACCATCTCCAAAGCCAAGATTGCCATTTGTCGGTC<br>ACCTGCATCTTTTGGACAAGCCTTTGTTGCACTACTCTTTAATCGACTTATCGAAGAGATACGGTCCATT<br>GTATTCTTTGTACTTCGGTTCCATGCCAACCGTCGTCGCTTCCACTCCAGAATTGTTCAAGTTGTTTTTG<br>CAAACCCACGAAGCCAGCTCTTTTAACACCAGATTCCAAACTTCTGCTATCAGAAGATTGACTTACGAC<br>AACTCCGTTGCTATGGTTCCATTCGGCCCATACTGGAAGTTCATTAGAAAGCTAATCATGAACGACTTG<br>TTAAACGCTACCACTGTTAACAAGTTAAGACCATTGAGATCCCAAGAAATTAGAAAGGTTTTGAGAGT<br>TATGGCTCAATCCGCTGAATCTCAAGTTCCATTGAACGTTACTGAAGAATTGCTAAAATGGACCAATTC<br>TACTATTAGTAGAATGATGTTGGGTGAAGCTGAAGAGATCAGAGACATTGCCAGAGATGTCTTGAAGA<br>TTTTTGGTGAATACTCTTTGACCGATTTCATCTGGCCATTGAAGAAGTTGAAGGTTGGTCAATACGAAA<br>AGAGAATTGATGACATCTTCAACAGATTCGATCCAGTCATTGAAAGAGTCATCAAAAAAAGGCAAGA<br>AATTAGAAAGAAGAGAAAAGAAAGAAACGGTGAGATAGAAGAAGGTGAACAATCTGTCGTCTTCTT<br>GGACACTTTGTTAGATTTCGCTGAAGATGAAACCATGGAAATTAAAATCACCAAGGAACAAATCAAG<br>GGTTTAGTTGTTGATTTCTTCTCTGCTGGTACTGACTCCACTGCTGTTGCTACCGACTGGGCCTTGTCT<br>GAATTGATCAACAACCCAAGAGTTTTCCAAAAGGCTAGAGAAGAAATTGATGCTGTTGTTGGTAAGG<br>ACAGATTAGTTGACGAAGCTGATGTTCAAAACTTACCTTACATCAGATCTATTGTTAAGGAAACTTTCA<br>GAATGCACCCACCACTACCAGTTGTCAAGCGTAAGTGTGTCCAAGAATGTGAAGTTGATGGTTACGTT<br>ATCCCAGAAGGTGCTTTGATTTTGTTTAATGTCTGGGCTGTAGGTAGAGACCCAAAATACTGGGACAG<br>ACCAACCGAATTTCGTCCTGAAAGATTCTTGGAAAACGTCGGTGAAGGTGACCAAGCTGTTGACTTG<br>AGAGGTCAACACTTCCAATTGCTACCTTTCGGTTCTGGTCGTAGAATGTGTCCAGGTGTCAACTTGGC<br>TACTGCTGGTATGGCCACATTATTGGCTTCCGTGATCCAATGCTTCGATTTGTCCGTTGTCGGTCCACAA<br>GGTAAGATCTTGAAGGGTAACGATGCTAAGGTCTCCATGGAAGAACGTGCCGGTTTGACTGTCCCAA<br>GAGCTCACAACTTGATCTGTGTCCCAGTTGCCCGTTCTTCTGCCGTTCCAAAGTTATTCTCCTCATAA |
| opAmIFS | ATGTTGTTGGAATTGGCTGTTACTTTGTTGGTTATTGCTTTGTTTATTCATTTGAGACCAACTCCATCTG<br>CTAAATCCAAGGCTTTGAGACATTTGCCAAATCCACCATCTCCTAAACCAAGATTGCCATTTATTGGTCA<br>TTTGCATTTGTTGGATAAGCCATTGTTACATCAATCCTTGATTAGATTGGGTGAAAGATATGGTCCATTA<br>TATAGTTTGTACTTCGGTTCTATGCCATGTGTTGTTGCTTCTACTCCTGAATTATTTAAGTTGTTTTTGCA<br>AACCCATGAAGCTTCTTCTTTTAATACTAGATTCCAAACTTCCGCTATTAGAAGATTGACTTATGATAACT<br>CCGTTGCTATGGTTCCATTTGGTCCATATTGGAAATTTATTAGAAAGTTGATTATGAACGATTTGTTGAA<br>CGCTACAACTGTTAACAAGTTGAGACCATTGAGATCTCAAGAAATTAGAAAAGTTTTGAACGTTATGG<br>CTAAGTCCGCTGAAGCTCAACAACCATTGAACGTTACTGAAGAATTGTTGAAATGGACTAATTCAACA<br>ATTTCCAGAATGATGTTGGGTGAAGCAGAAGAAATTAGAGATATTGCTAGAGATGTTTTGAAGATTTT<br>CGGTGAATATTCATTGACTGATTTCATTTGGCCATTGAAAAAGTTTAAGGTTGGTCAATACGAAAAGA<br>GAATTGATGATATTTTCAACAGATTCGATCCAGTTATTGAAAAGGTTATTAAGAAGAGACAAGAAATTA<br>TTAAGAGAAGAAAGGAAAGAAACGGTGAATTGGAAGAAGGTGAACAAAGTGTTGTTTTTTTGGATA<br>CATTGTTGCAATACGCTGAAGATGAAACTATGGAAATTAAAATTACCAAGGAACAAATTAAGGGTTTG<br>GTTGTTGATTTTTTCAGTGCTGGTACTGATAGTACTGCTGTTGCTACTGATTATGCTTTAGCTGAATTGA<br>TTAACAACCCAAAGGTTTTGAGAAAGGCTAGAGAAGAAGTTGATACTGTTGTTGGTAAGGATAGATT<br>GGTTGATGAAAGTGATGTTCAACATTTGCATTATATTAGAGCTATTGTTAAGGAAACCTTCAGAATGCA<br>TCCACCATTGCCAGTTGTTAAAAGAAAATGTACTCAAGATTGTGAAATTGATGGTTTTGTTATTCCAGA<br>AGGTGCTTTAATTTTGTTTAACGTTTGGGCAGTTGGTAGAGATCCAAAATATTGGGATAGACCATCTGA<br>ATTTTTGCCTGAAAGATTTTTGGAAAAGGCTGGTGGTGAAGGTGAAGTTGGTCCAATTGATTTGAGA<br>GGTCAACATTTTCAATTGTTGCCATTTGGTAGTGGTAGAAGAATGTGTCCAGGTGTTAACTTGGCAAC<br>TGCTGGTATGGCTACTTTGTTGGCTTCTGTTATTCAAACCTTTGATTTGCAAGTTCCAGGTCCTCAAGG<br>TCAAATTTTGAAAGGTGATGAAGCTAAAGTTTCTATGGAAGAAAGAGCAGGTTTGACAGTTCCAAGA<br>GCTCATAATTTGATTTGTGTTCCTTTGGCTAGAGCTGGTGTTGCTGCTAAGTTATTGTCTAGTTAA |
| opGmHID | ATGGCCAAGGAAATTGTTAAGGAATTATTGCCATTGATCAGAGTTTACAAGGATGGTTCCGTTGAAAG<br>ATTGTTGTCCTCTGAAAACGTTGCTGCCAGTCCAGAAGACCCACAAACTGGTGTCTCTTCCAAGGACA<br>TTGTCATTGCTGATAATCCATACGTTTCCGCTAGAATCTTCTTGCCAAAGTCTCACCACACCAACAACAA |

| | |
|---|---|
| | GTTGCCAATTTTTTTGTACTTCCACGGTGGTGCTTTCTGTGTTGAATCTGCTTTCTCTTTCTTTGTTCAC<br>AGATACTTGAACATCCTAGCATCTGAAGCCAACATCATCGCCATTTCTGTTGATTTCCGTCTGTTACCAC<br>ACCATCCAATCCCAGCTGCTTACGAAGACGGTTGGACTACATTGAAATGGATTGCTTCCCACGCTAATA<br>ACACCAACACTACTAACCCAGAACCATGGTTATTAAACCACGCTGACTTCACCAAGGTCTACGTCGGT<br>GGTGAAACTTCTGGTGCTAACATTGCTCACAACTTGTTGTTGAGAGCTGGTAACGAATCCTTGCCTGG<br>TGACTTGAAGATCTTGGGTGGTTTGTTGTGTTGTCCATTCTTCTGGGGTTCTAAACCTATTGGTTCCGA<br>AGCTGTCGAAGGTCACGAACAATCTTTGGCTATGAAGGTTTGGAACTTTGCTTGCCCAGACGCTCCA<br>GGCGGTATTGACAACCCATGGATCAACCCATGTGTCCCAGGTGCCCCATCTTTGGCTACCTTGGCTTGT<br>TCAAAATTGTTGGTTACTATCACCGGTAAGGACGAATTCAGAGACCGTGATATCTTATATCATCACACTG<br>TCGAGCAATCCGGTTGGCAAGGTGAATTGCAATTGTTCGATGCCGGTGATGAAGAACACGCCTTCCA<br>ATTATTCAAGCCAGAAACCCATTTGGCCAAGGCTATGATCAAGAGATTAGCTTCTTTCTTGGTCTAA |

**Supplementary Table 10**. List of ion pairs used for metabolite detection by MRM

| Metabolite | Ion mode | Collision energy CE (eV) | Ion pair (*m/z*) |
|---|---|---|---|
| Isoliquiritigenin | Negative | 25 | 255.15>119.10 |
| Liquiritigenin | Positive | 25 | 257.00>136.95 |
| Naringenin | Negative | 25 | 271.00>119.15 |
| Daidzein | Positive | 35 | 254.95>198.95 |
| 2’-OH daidzein | Positive | 35 | 271.05>200.95 |
| 2’-OH dihydrodaidzein | Positive | 35 | 273.05>107.00 |
| 7,2’,4’-trihydroxyisoflavanol | Positive | 35 | 257.05>123.00 |
| 3, 9-OH peterocarpan | Positive | 35 | 257.05>123.00 |
| Glycinol | Positive | 35 | 255.10>152.00 |
| 4-Glyceollidin | Positive | 35 | 323.10>267.05 |
| Glyceocarpin | Positive | 35 | 323.10>267.05 |
| Glyceollin I | Positive | 35 | 339.10>155.05 |
| Glyceollin II | Positive | 35 | 339.10>155.05 |
| Glyceollin III | Positive | 35 | 339.10>155.05 |

#### 2. Synthesis of compounds

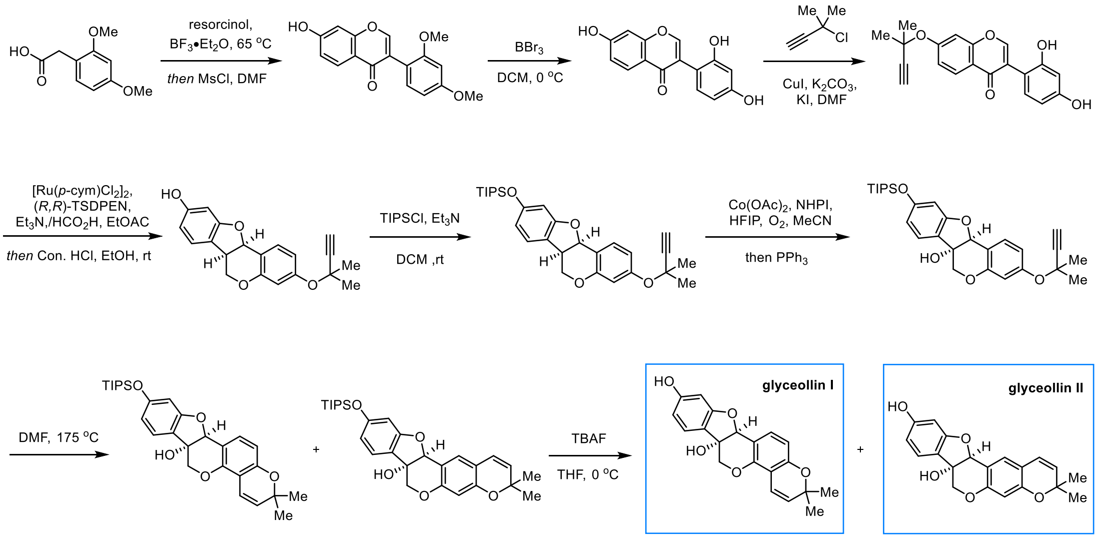

Approximately 30 mg **glyceollin I** (**10**) and 8 mg **glyceollin II** (**11**) were synthesized according to the reported method (Ciesielski and Metz, 2020) .

Data of glyceollin I (**10**): **$^{1}$H-NMR** (500 MHz, Methanol-d4) δ 7.21 (d, *J* = 8.4 Hz, 1H), 7.17 (d, *J* = 8.2 Hz, 1H), 6.60 (d, *J* = 10.0 Hz, 1H), 6.47 (dd, *J* = 8.4, 0.6 Hz, 1H), 6.40 (dd, *J* = 8.1, 2.1 Hz, 1H), 6.23 (d, *J* = 2.1 Hz, 1H), 5.61 (d, *J* = 10.0 Hz, 1H), 5.17 (s, 1H), 4.17 (dd, *J* = 11.4, 1.0 Hz, 1H), 3.94 (d, *J* = 11.4 Hz, 1H), 1.38 (d, *J* = 5.3 Hz, 6H);
**$^{13}$C-NMR** (126 MHz, Methanol-d4) δ 162.2, 161.2, 155.3, 151.9, 132.2, 130.3, 125.2, 121.2, 117.5, 114.2, 111.6, 111.3, 109.4, 99.0, 85.9, 77.2, 77.1, 71.1, 28.1;
**HRMS (ESI)** $[M+H]^+$ calculated for $C_{20}H_{19}O_5$ :339.1233, found 339.1236.

Data of glyceollin II (**11**): **$^{1}$H-NMR** (500 MHz, Acetone-d6) δ 8.48 (s, 1H), 7.20 (d, J = 8.1 Hz, 1H), 7.14 (s, 1H), 6.49 – 6.37 (m, 2H), 6.25 (d, J = 2.1 Hz, 1H), 6.21 (s, 1H), 5.65 (d, J = 9.9 Hz, 1H), 5.25 (s, 1H), 4.96 (s, 1H), 4.13 (dd, J = 11.4, 0.8 Hz, 1H), 4.04 (d, J = 11.4 Hz, 1H), 1.39 (s, 4H), 1.36 (s, 3H);
**$^{13}$C-NMR** (126 MHz, Acetone-d6) δ 162.0, 160.8, 156.8, 155.3, 129.9, 129.8, 125.2, 122.3, 121.4, 117.2, 114.5, 109.0, 104.8, 98.7, 85.8, 77.2, 76.7, 70.6, 28.3, 28.3;
**HRMS (ESI)** $[M+H]^+$ calculated for $C_{20}H_{19}O_5$ :339.1233, found 339.1233.

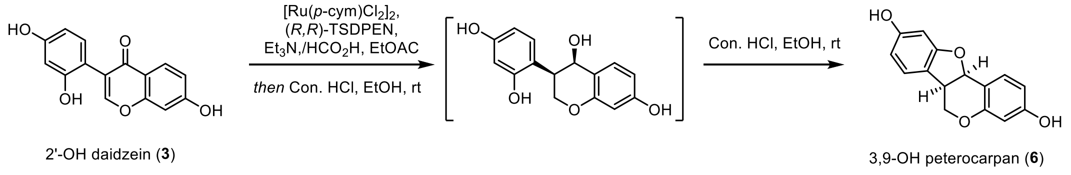

3,9-OH peterocarpan (**6**) were synthesized according to the reported method (Ciesielski and Metz, 2020).

Catalyst solution: $[Ru(p\text{-cym})Cl_2]_2$ (45.2 mg, 73.8 μmol) and (*R,R*)-TsDPEN (60.8 mg, 162.4 μmol) were dissolved in DMSO (0.88 mL) in a 5 mL round-bottomed flask. Another flask containing TEA (3.6 mL) was cooled to 0 °C, and $HCO_2H$ (1.2 mL) was added. The mixture was stirred vigorously for 5 min at room temperature. An aliquot of this mixture (2.4 mL) was added to the ruthenium catalyst, and the resulting solution was stirred for another 3 min.

To a stirred solution of 2'-OH daidzein (**3**) 10 (542.6 mg, 2.0 mmol) in DMSO (2.5 mL) and water (0.36 mL) was added the catalyst solution (2.2 mL). The reaction mixture was stirred at 45 °C for 19.5 h, cooled to room temperature, and hydrochloric acid (37 %, 1.04 mL, 12.4 mmol, 6.2 equiv.) was added. The mixture was stirred for 10 min. The reaction was quenched by addition of saturated aqueous $NH_4Cl$ solution, and the mixture was extracted three times with EtOAc. The combined organic layers were dried over $Na_2SO_4$, the solvents were removed in vacuo, and the residue was purified by flash chromatography (PE : EA=3:1 v/v) to afford 3,9-OH peterocarpan (**6**) (281.6 mg, 55%) as a white solid.

Data of 3,9-OH peterocarpan (**6**): **$^{1}$H-NMR** (500 MHz, acetone-*d6*) δ 8.55 (s, 1H), 8.31 (s, 1H), 7.31 (d, *J* = 8.4 Hz, 1H), 7.13 (d, *J* = 8.0 Hz, 1H), 6.55 (dd, *J* = 8.4, 2.4 Hz, 1H), 6.48 – 6.33 (m, 2H), 6.28 (d, *J* = 2.2 Hz, 1H), 5.47 (d, *J* = 6.4 Hz, 1H), 4.42 – 4.17 (m, 1H), 3.70 – 3.38 (m, 2H); **$^{13}$C-NMR** (126 MHz, acetone-*d6*) δ 161.8, 159.7, 159.6, 157.7, 133.1, 125.9, 119.2, 112.91, 110.5, 108.3, 103.9, 98.5, 79.3, 67.2, 40.3;

**HRMS (ESI)** $[M+H]^+$ calculated for $C_{15}H_{13}O_4$ :257.0814, found 257.0820.

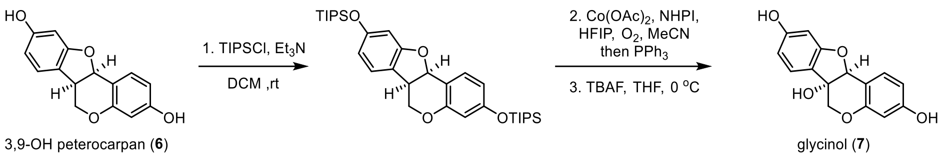

To a solution of 3,9-OH peterocarpan (**6**) (34 mg, 0.13 mmol) in DCM (2 mL) DMAP (3.3 mg, 0.027 mmol, 0.2 equiv.), triethylamine (0.11 mL, 0.789 mmol, 6.0 equiv.) and TIPSCl (0.17 mL, 0.789 mmol, 6 equiv.) were added subsequently. The reaction was stirred for 19 h at room temperature and quenched by addition of saturated aqueous $NH_4Cl$ solution. The mixture was extracted three times with DCM, the combined organic layers were dried over $Na_2SO_4$, the solvents were removed in vacuo, and the residue was used directly in the next step.

The above residue was dissolved in MeCN (5 mL), then N-hydroxyphthalimide (21.2 mg, 0.13 mmol, 1.0 equiv.), cobalt(II) acetate (11.5 mg, 0.065 mmol, 0.5 equiv.) and 1,1,1,3,3,3-hexafluoropropan-2-ol (0.07 mL, 0.59 mmol, 4.5 equiv.) were added. The argon atmosphere was replaced with air (balloon), and the reaction mixture was stirred for overnight at room temperature. The air atmosphere was replaced with argon, and triphenylphosphine (34.1 mg, 0.13 mmol, 1.0 equiv.) was added. After additional stirring for 45 min, the reaction was quenched by addition of saturated aqueous $NH_4Cl$ solution. The mixture was extracted three times with EtOAc, the combined organic layers were dried over $Na_2SO_4$, the solvents were removed in vacuo, and the residue was redissolved in THF (5 mL,) and TBAF (0.13 mL, 0.13 mol, 1M in THF) was added at 0 $^{o}$C. After stirring at 0 ºC for 1 h, the reaction was quenched by sat. $NH_4Cl$ (5 mL) and the resulting mixture was extracted with ethyl acetate (3 x 5 mL). The combined organic phases were washed with brine (5 mL), dried over $Na_2SO_4$, filtered and concentrated *in vacuo*. The residue was purified on silica gel chromatography (Petroleum ether/Ethyl acetate = 1/1) to provide glycinol (**7**) 10.2 mg, 29%) as a colorless oil.

Data of glycinol (**7**): **$^{1}$H-NMR** (500 MHz, acetone-*d*$_6$) δ 8.56 (s, 1H), 8.48 (s, 1H), 7.30 (dd, J = 8.4, 0.6 Hz, 1H), 7.20 (d, J = 8.1 Hz, 1H), 6.55 (dd, J = 8.4, 2.4 Hz, 1H), 6.42 (dd, J = 8.1, 2.1 Hz, 1H), 6.32 (d, J = 2.4 Hz, 1H), 6.24 (d, J = 2.1 Hz, 1H), 5.26 (s, 1H), 4.94 (s, 1H), 4.12 (dd, J = 11.3, 0.7 Hz, 1H), 4.02 (d, J = 11.4 Hz, 1H);

**$^{13}$C-NMR** (126 MHz, acetone-*d*$_6$) δ 162.0, 160.7, 159.6, 157.1, 133.2, 125.1, 121.5, 113.3, 110.7, 108.8, 103.8, 98.6, 85.9, 76.7, 70.6;

**HRMS (ESI)** $[M+H]^+$ calculated for $C_{15}H_{13}O_5$ :273.0763, found 273.0762.

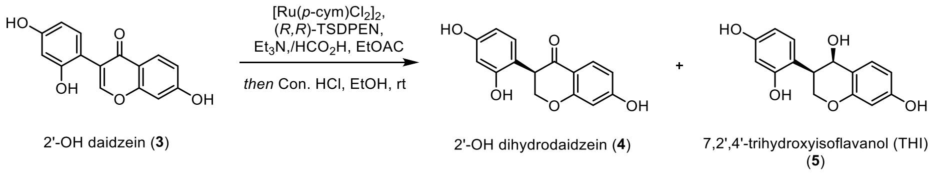

2'-OH dihydrodaidzein (**4**) and 7,2',4'-trihydroxyisoflavanol (THI) (**5**) were synthesized according to the reported method (Ciesielski and Metz, 2020).

Catalyst solution: [Ru(*p*-cym)$Cl_2$]$_2$ (45.2 mg, 73.8 µmol) and (*R,R*)-TsDPEN (60.8 mg, 162.4 µmol) were dissolved in DMSO (0.88) in a 5 mL round-bottomed flask. Another flask containing TEA (3.6 mL) was cooled to 0 °C, and $HCO_2H$ (1.2 mL) was added. The mixture was stirred vigorously for 5 min at room temperature. An aliquot of this mixture (2.4 mL) was added to the ruthenium catalyst, and the resulting solution was stirred for another 3 min.
To a stirred solution of 2'-OH daidzein (**3**) (542.6 mg, 2.0 mmol) in DMSO (2.5 mL) and water (0.36 mL) was added the catalyst solution (2.2 mL). The reaction mixture was stirred at 45 °C for 19.5 h, cooled to room temperature. The reaction was quenched by addition of saturated aqueous $NH_4Cl$ solution, and the mixture was extracted three times with EtOAc. The combined organic layers were dried over $Na_2SO_4$, the solvents were removed in vacuo, and the residue was purified by flash chromatography (DCM:EtOAc 3:1 v/v) to afford 2'-OH dihydrodaidzein (**4**) (90.1 mg,) and 7,2',4'-trihydroxyisoflavanol (THI) (**5**) (191.8 mg) as a white solid.

Data of 2'-OH dihydrodaidzein (**4**): **$^1$H-NMR** (500 MHz, acetone-$d_6$) δ 9.40 (s, 1H), 8.51 (s, 1H), 8.19 (s, 1H), 7.77 (d, *J* = 8.6 Hz, 1H), 6.93 (d, *J* = 8.3 Hz, 1H), 6.58 (dd, *J* = 8.7, 2.3 Hz, 1H), 6.42 (dd, *J* = 11.1, 2.4 Hz, 2H), 6.31 (dd, *J* = 8.3, 2.4 Hz, 1H), 4.66 (dd, *J* = 11.1, 9.8 Hz, 1H), 4.53 (dd, *J* = 11.1, 5.2 Hz, 1H), 4.12 (dd, *J* = 9.8, 5.2 Hz, 1H);
**$^{13}$C-NMR** (126 MHz, acetone-$d_6$) δ 191.7, 165.1, 164.7, 158.7, 157.2, 131.2, 130.1, 115.6, 114.6, 111.3, 107.8, 103.8, 103.5, 71.7, 47.7;
**HRMS (ESI)** $[M+H]^+$ calculated for $C_{15}H_{13}O_5$ :273.0763, found 273.0763.

Data of 7,2',4'-trihydroxyisoflavanol (THI) (**5**): **$^1$H-NMR** (500 MHz, acetone-$d_6$) δ 8.75 (s, 1H), 8.31 (s, 1H), 8.10 (s, 1H), 7.07 (d, *J* = 8.3 Hz, 1H), 7.01 (d, *J* = 8.3 Hz, 1H), 6.41 – 6.36 (m, 2H), 6.34 – 6.28 (m, 2H), 4.79 (d, *J* = 3.4 Hz, 1H), 4.60 (dd, *J* = 12.3, 10.2 Hz, 1H), 4.50 (d, *J* = 4.2 Hz, 1H), 4.08 (ddd, *J* = 10.2, 3.7, 1.3 Hz, 1H), 3.44 (dt, *J* = 12.2, 3.3 Hz, 1H);
**$^{13}$C-NMR** (126 MHz, acetone-$d_6$) δ 159.3, 158.2, 157.3, 156.1, 132.3, 131.4, 118.1, 117.2, 108.8, 107.6, 104.1, 103.4, 66.9, 64.8, 40.7;
**HRMS (ESI)** $[M+H-H_2O]^+$ calculated for $C_{15}H_{15}O_6$ :257.0814, found 257.0813.

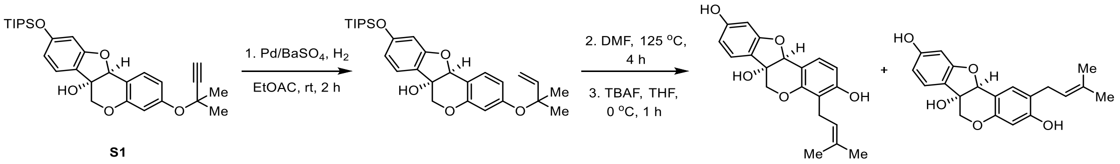

To a stirred solution of **S1** (40 mg, 0.08 mmol) in ethyl acetate (4 mL) was added Lindlar catalyst (4 mg) and quinoline (1 uL, 0.008 mmol, 0.1 eq). This suspension was degassed at -78 °C and backfilled with $H_2$ three times. After 2.5 h, the catalyst was removed by filtration and the filtrate was concentrated and used in the next step directly.

The residue was dissolved in DMF (2 mL) and heat at 125 °C for 4 hours, then the reaction was concentrated in vacuo. The residue was redissolved in THF (4 mL) and TBAF (120 uL, 120 mmol, 1.5 equiv.) was added 0 °C. After stirring at 0 ºC for 1 h, the reaction was quenched by sat. $NH_4Cl$ (5 mL) and the resulting mixture was extracted with ethyl acetate (3 x 5 mL). The combined organic phases were washed with brine (5 mL), dried over $Na_2SO_4$, filtered and concentrated *in vacuo*. The residue was purified on silica gel chromatography (Petroleum ether/Ethyl acetate = 1/1) to provide 4-glyceollidin (**8**) (16 mg, 59%) and glyceocarpin (**9**) (4 mg, 15%) as colorless oil. The spectroscopic data of **8** and **9** are in accordance with the literature reported values (Akashi et al., 2009; Yoneyama et al., 2016).

Data of 4-glyceollidin (**8**): **$^1$H-NMR** (500 MHz, acetone-*d*$_6$) δ 8.42 (s, 1H), 8.38 (s, 1H), 7.21 (d, *J* = 8.1 Hz, 1H), 7.14 (d, *J* = 8.4 Hz, 1H), 6.59 (d, *J* = 8.4 Hz, 1H), 6.42 (dd, *J* = 8.2, 2.1 Hz, 1H), 6.23 (d, *J* = 2.1 Hz, 1H), 5.27 (s, 1H), 5.20 – 5.16 (m, 0H), 4.90 (s, 1H), 4.14 (d, *J* = 11.3 Hz, 1H), 4.07 (s, 0H), 1.72 (s, 2H), 1.59 (s, 2H);

**$^{13}$C-NMR** (126 MHz, acetone-*d*$_6$) δ 162.1, 160.7, 156.8, 155.0, 131.1, 129.8, 125.2, 123.9, 121.7, 116.6, 113.7, 110.2, 108.8, 98.6, 86.7, 76.9, 71.2, 25.9, 22.9, 17.9;

**HRMS (ESI)** $[M+H]^+$ calculated for $C_{20}H_{21}O_5$ :341.1389, found 341.1392.

Data of glyceocarpin (**9**): **$^1$H-NMR** (500 MHz, acetone-*d*$_6$) δ 8.51 (s, 1H), 8.45 (s, 1H), 7.18 (d, *J* = 8.2 Hz, 1H), 7.16 (s, 1H), 6.41 (dd, *J* = 8.2, 2.1 Hz, 1H), 6.34 (s, 1H), 6.24 (d, *J* = 2.1 Hz, 1H), 5.39 – 5.30 (m, 1H), 5.23 (s, 1H), 4.88 (s, 1H), 4.08 (dd, *J* = 11.3, 0.8 Hz, 1H), 3.97 (d, *J* = 11.3 Hz, 1H), 1.73 (q, *J* = 1.4 Hz, 6H);

**$^{13}$C-NMR** (126 MHz, acetone-*d*$_6$) δ 206.1, 161.9, 160.6, 156.9, 155.0, 132.5, 132.2, 125.04, 123.8, 123.1, 121.6, 112.9, 108.8, 103.4, 98.6, 86.0, 76.8, 70.6, 30.2, 30.1, 29.9, 29.8, 29.6, 29.5, 29.3, 28.4, 25.9, 17.8;

**HRMS (ESI)** $[M+H]^+$ calculated for $C_{20}H_{21}O_5$ :341.1389, found 341.1380.

### 3. NMR Spectra of compounds

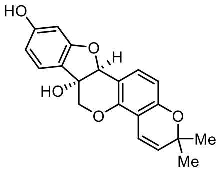

**glyceollin I**

$^{1}$H-NMR, MeOH-*d4*
500 MHz, 298K

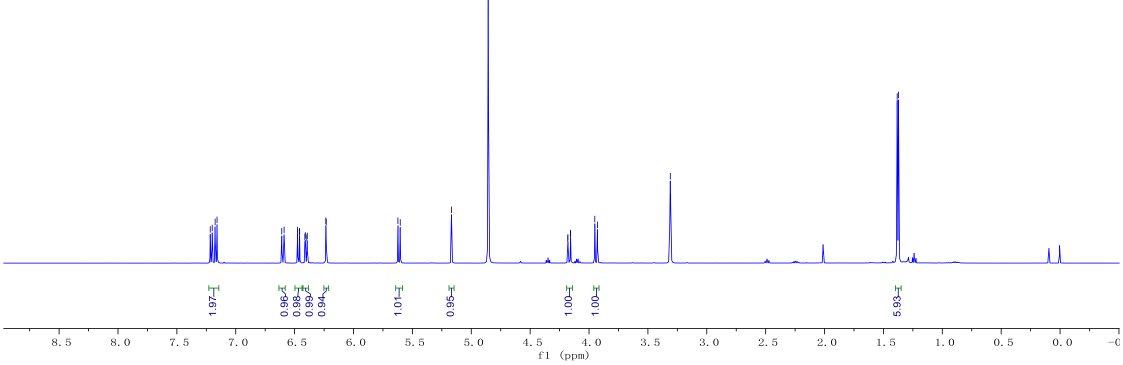

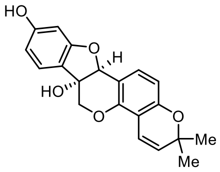

**glyceollin I**

$^{13}$C-NMR, MeOH-*d4*
126 MHz, 298K

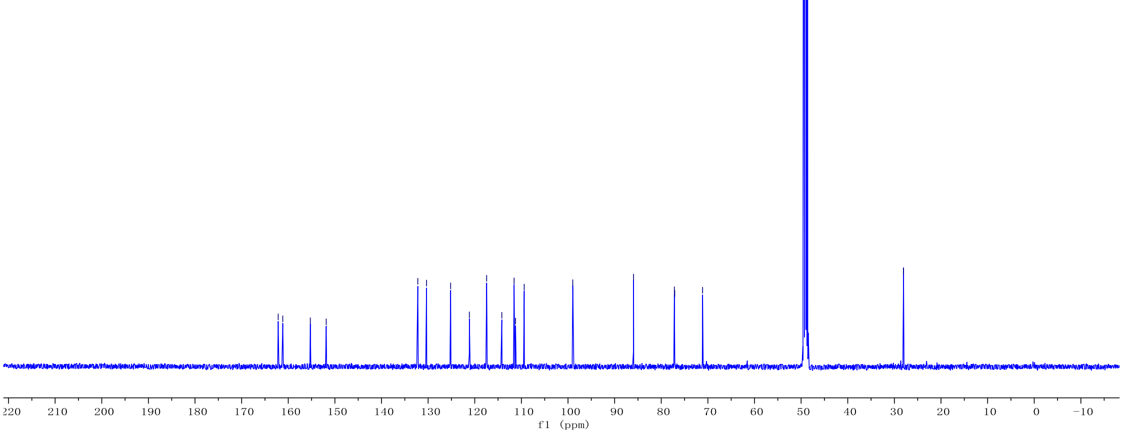

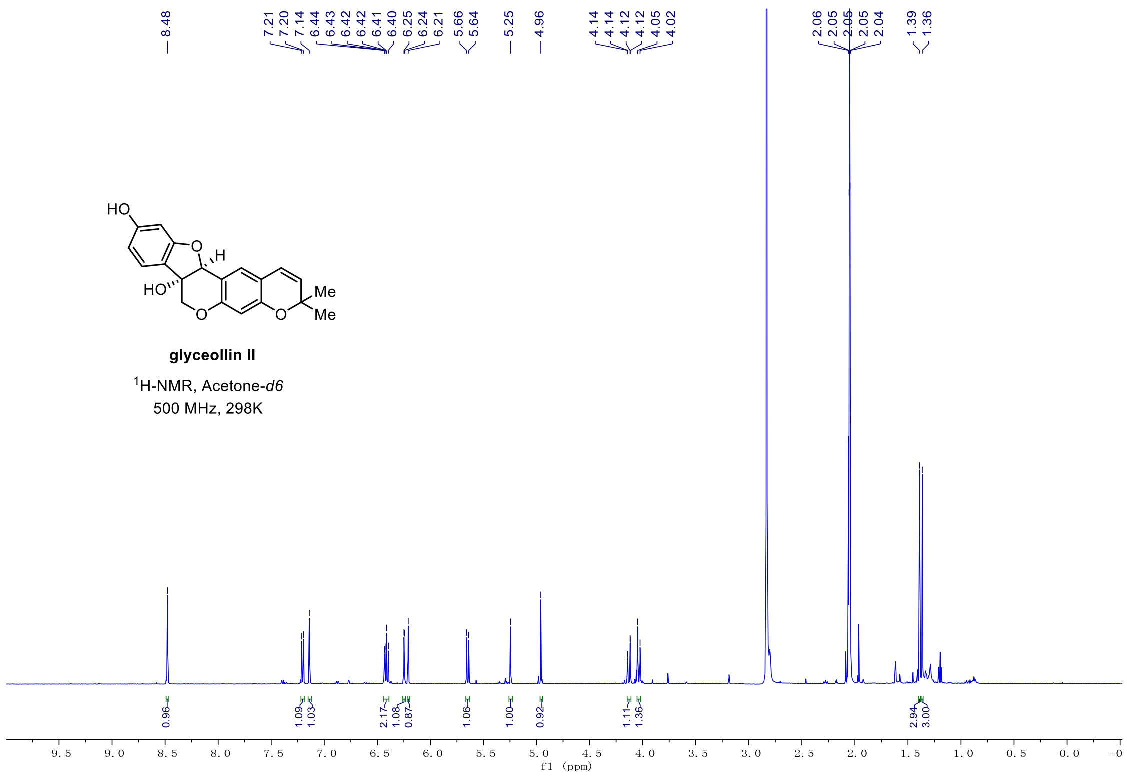
HO
H
HO
Me
Me
O
O
O
glyceollin II
1H-NMR, Acetone-d6
500 MHz, 298K
f1 (ppm)

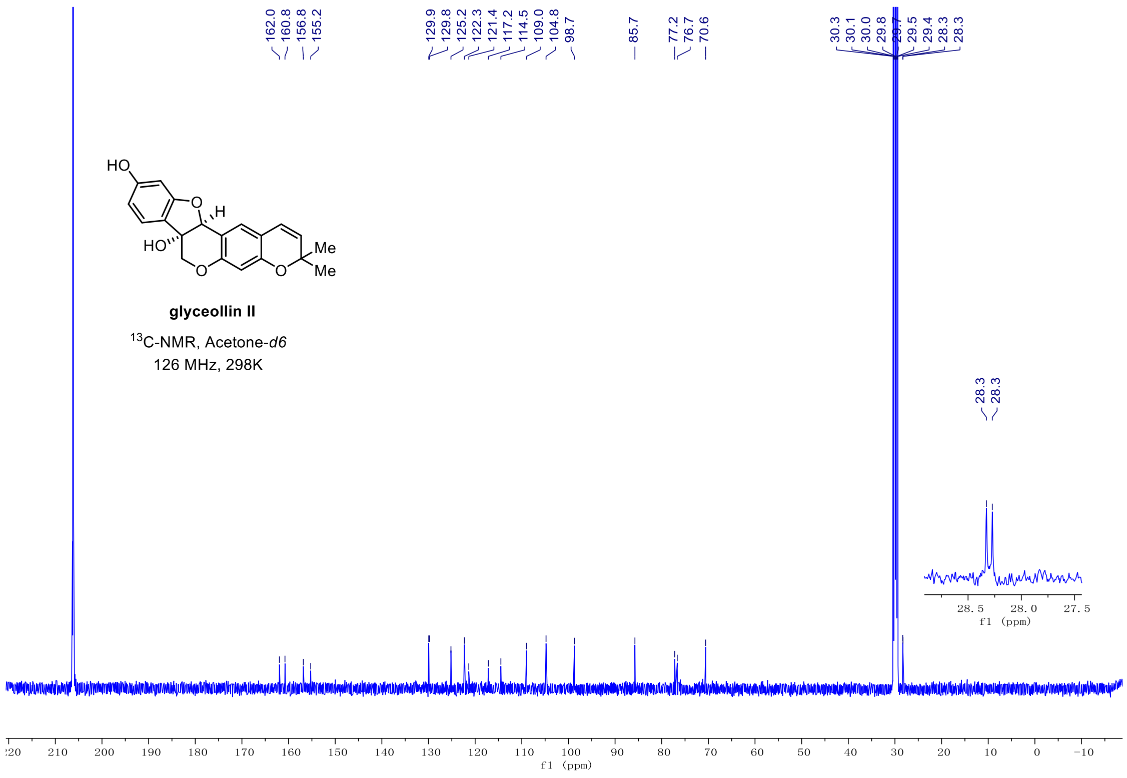
HO
H
HO
Me
Me
O
O
O
glyceollin II
13C-NMR, Acetone-d6
126 MHz, 298K
f1 (ppm)

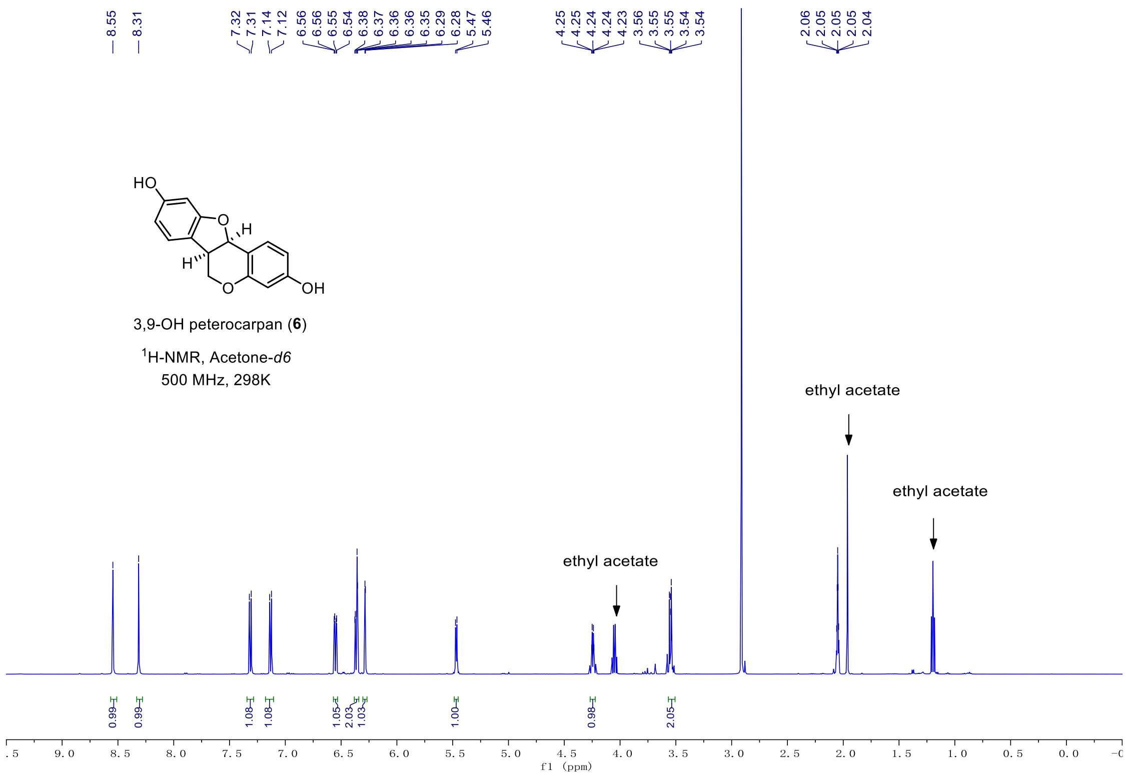
8.55
8.31
7.32
7.31
7.14
7.12
6.56
6.56
6.55
6.54
6.38
6.37
6.36
6.36
6.35
6.29
6.28
5.47
5.46
4.25
4.25
4.24
4.24
4.23
3.56
3.55
3.55
3.54
3.54
2.06
2.05
2.05
2.05
2.04
HO
O
H
H
O
OH
3,9-OH peterocarpan (6)
1H-NMR, Acetone-d6
500 MHz, 298K
ethyl acetate
ethyl acetate
ethyl acetate
0.99
0.99
1.08
1.08
1.05
2.03
1.03
1.00
0.98
2.05
f1 (ppm)

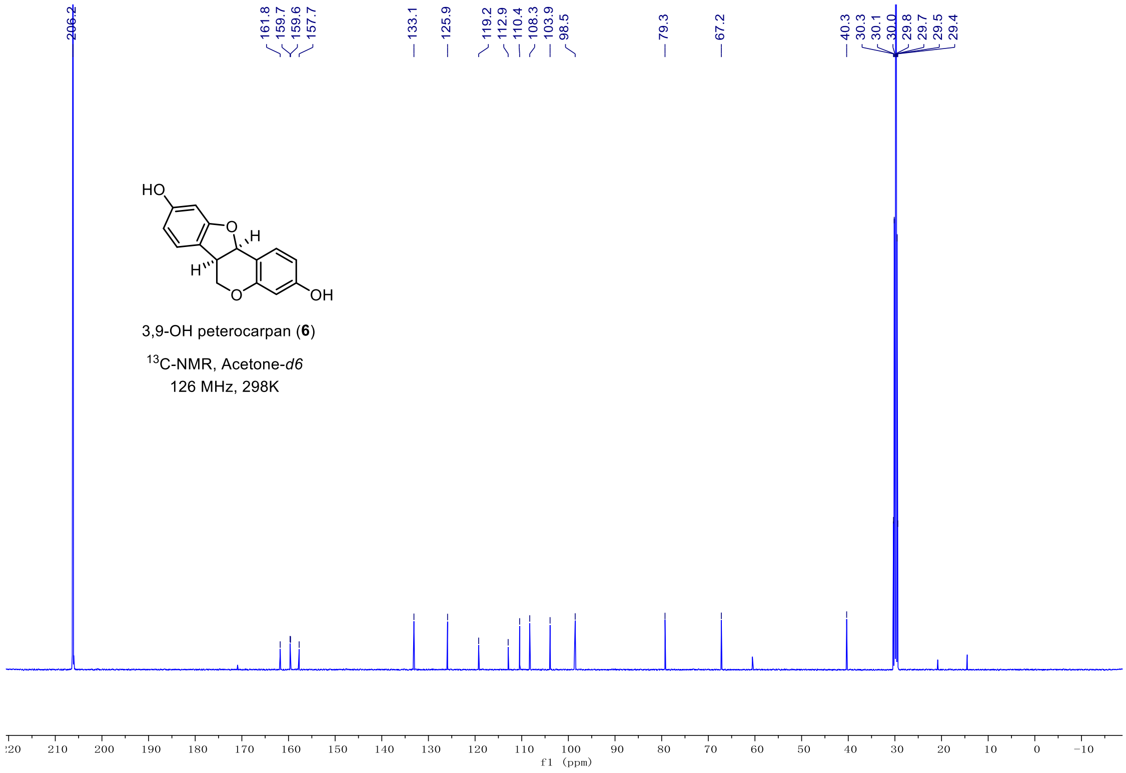
206.2
161.8
159.7
159.6
157.7
133.1
125.9
119.2
112.9
110.4
108.3
103.9
98.5
79.3
67.2
40.3
30.3
30.1
30.0
29.8
29.7
29.5
29.4
HO
O
H
H
O
OH
3,9-OH peterocarpan (6)
13C-NMR, Acetone-d6
126 MHz, 298K
f1 (ppm)

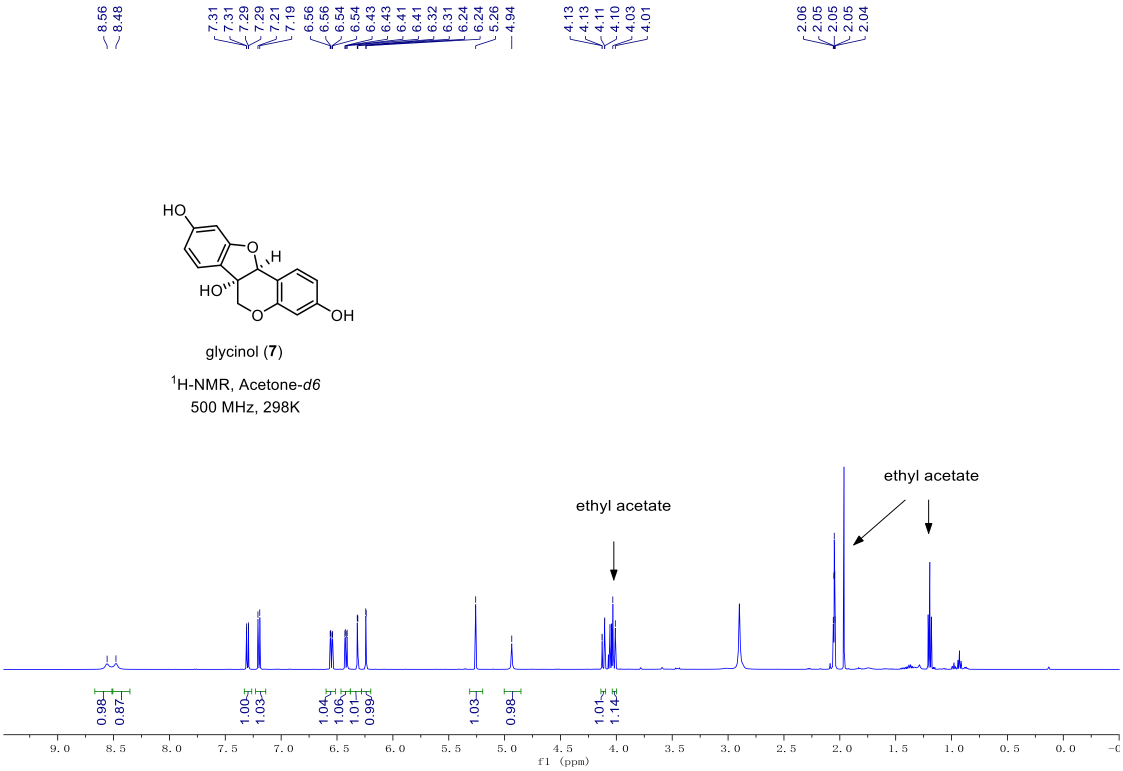

8.56
8.48
7.31
7.31
7.29
7.29
7.21
7.19
6.56
6.56
6.54
6.54
6.43
6.43
6.41
6.41
6.32
6.31
6.24
6.24
5.26
4.94
4.13
4.13
4.11
4.10
4.03
4.01
2.06
2.05
2.05
2.05
2.04
HO
O
H
HO
O
OH
glycinol (7)
1H-NMR, Acetone-d6
500 MHz, 298K
ethyl acetate
ethyl acetate
0.98
0.87
1.00
1.03
1.04
1.06
1.01
0.99
1.03
0.98
1.01
1.14
f1 (ppm)

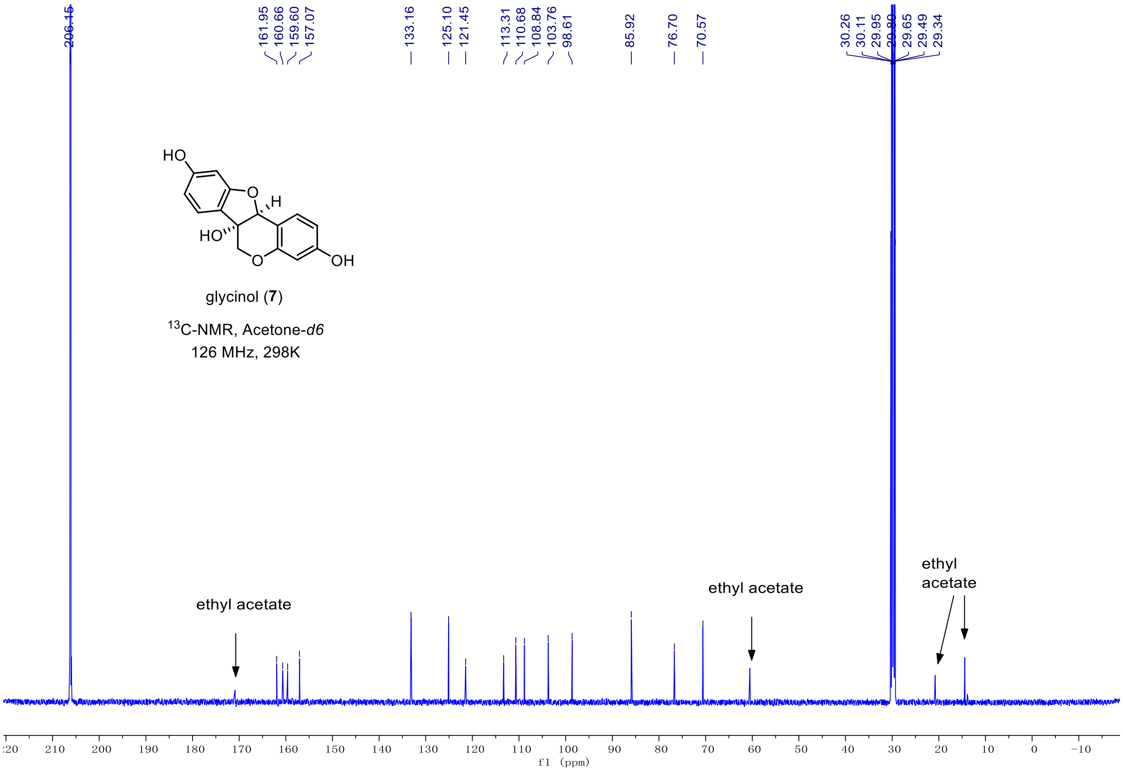

206.15
161.95
160.66
159.60
157.07
133.16
125.10
121.45
113.31
110.68
108.84
103.76
98.61
85.92
76.70
70.57
30.26
30.11
29.95
29.65
29.49
29.34
HO
O
H
HO
O
OH
glycinol (7)
13C-NMR, Acetone-d6
126 MHz, 298K
ethyl acetate
ethyl acetate
ethyl acetate
f1 (ppm)

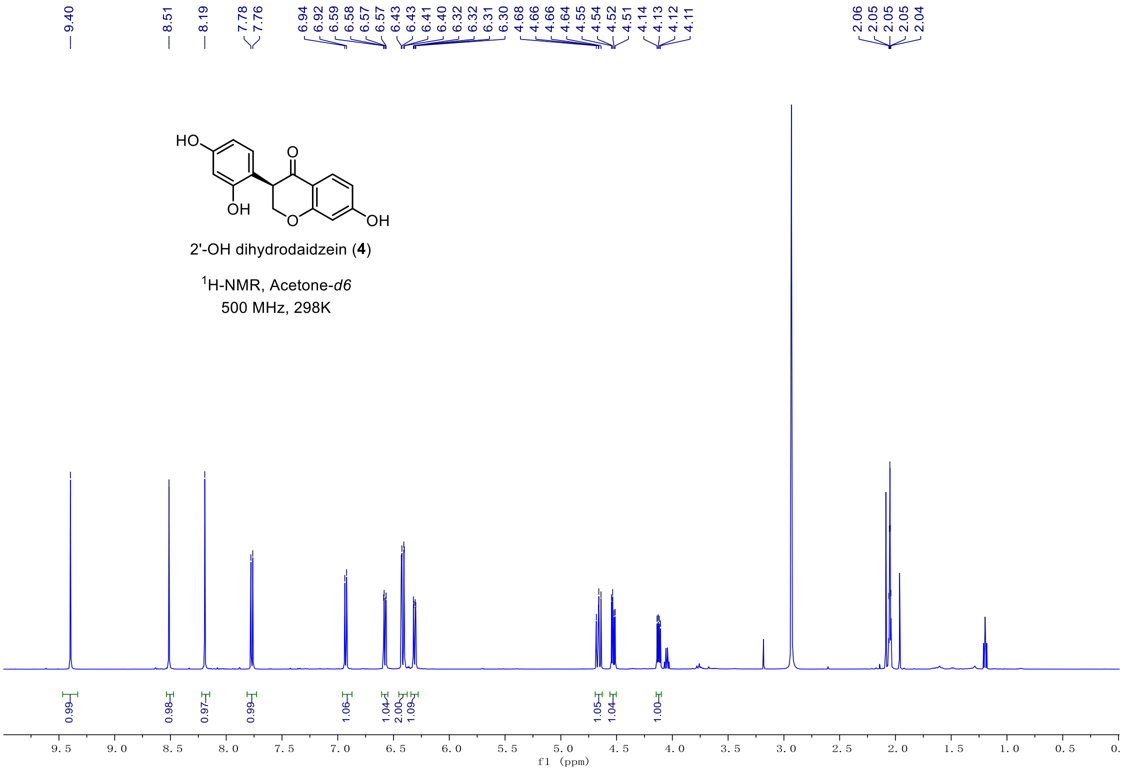
9.40
8.51
8.19
7.78
7.76
6.94
6.92
6.59
6.58
6.57
6.57
6.43
6.43
6.41
6.40
6.32
6.32
6.31
6.30
4.68
4.66
4.64
4.55
4.54
4.52
4.51
4.14
4.13
4.12
4.11
2.06
2.05
2.05
2.05
2.04
HO
O
OH
O
OH
2'-OH dihydrodaidzein (4)
1H-NMR, Acetone-d6
500 MHz, 298K
0.99
0.98
0.97
0.99
1.06
1.04
2.00
1.09
1.05
1.04
1.00
f1 (ppm)

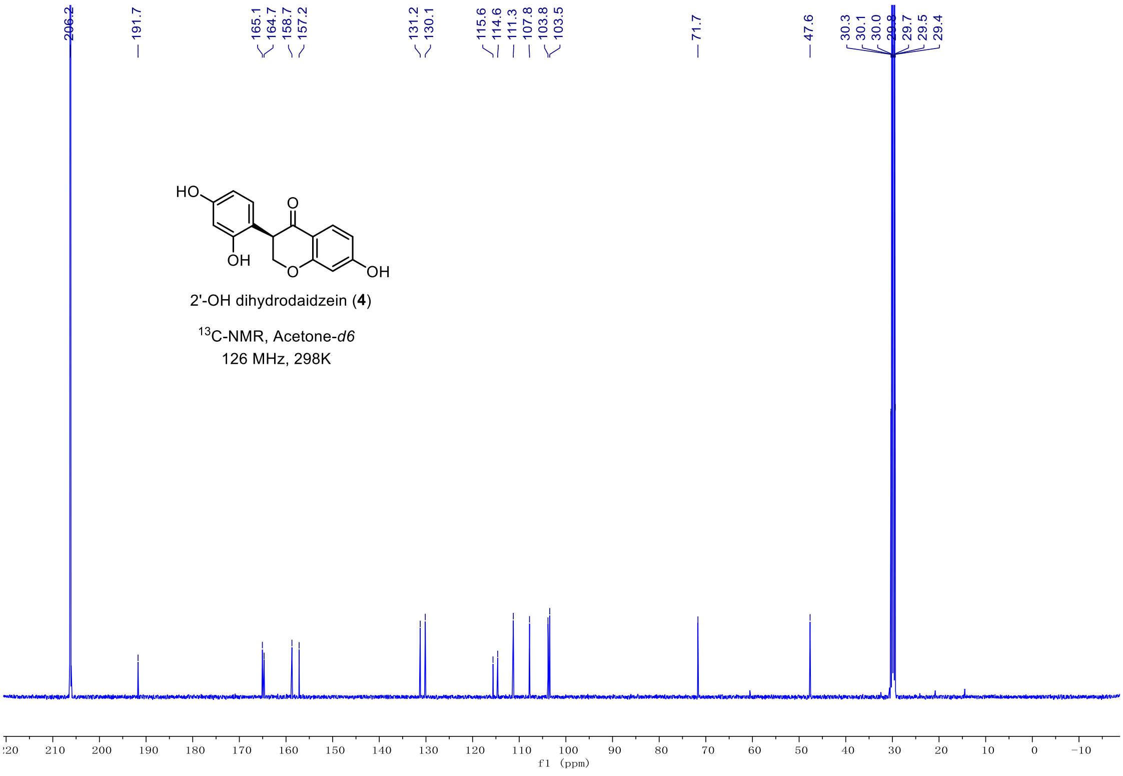
206.2
191.7
165.1
164.7
158.7
157.2
131.2
130.1
115.6
114.6
111.3
107.8
103.8
103.5
71.7
47.6
30.3
30.1
30.0
29.8
29.7
29.5
29.4
HO
O
OH
O
OH
2'-OH dihydrodaidzein (4)
13C-NMR, Acetone-d6
126 MHz, 298K
f1 (ppm)

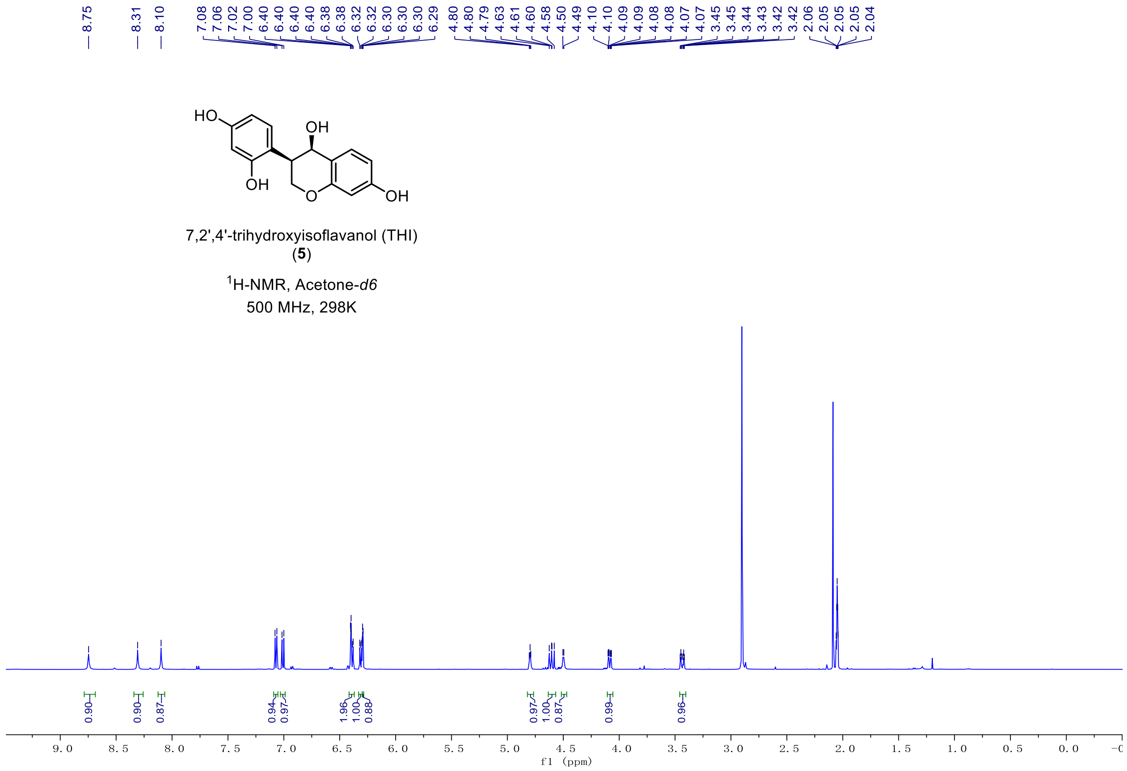
8.75
8.31
8.10
7.08 7.06 7.02 7.00 6.40 6.40 6.40 6.40 6.38 6.38 6.32 6.32 6.30 6.30 6.30 6.29
4.80 4.80 4.79 4.63 4.61 4.60 4.58 4.50 4.49
4.10 4.09 4.09 4.08 4.08 4.07 4.07
3.45 3.45 3.44 3.43 3.43 3.42
2.06 2.05 2.05 2.05 2.04
HO
OH
OH
O
OH
7,2',4'-trihydroxyisoflavanol (THI)
(5)
1H-NMR, Acetone-d6
500 MHz, 298K
0.90 0.90 0.87 0.94 0.97 1.96 1.00 0.88 0.97 1.00 0.87 0.99 0.96
f1 (ppm)

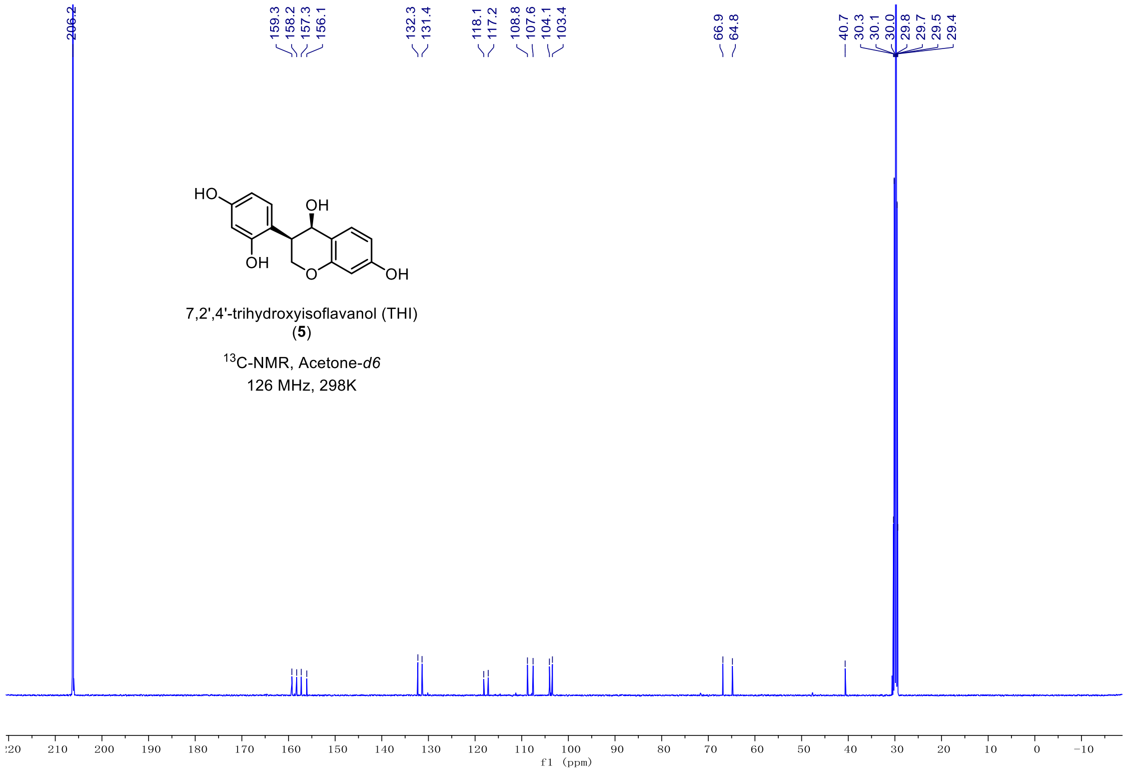
206.2
159.3 158.2 157.3 156.1
132.3 131.4
118.1 117.2
108.8 107.6 104.1 103.4
66.9 64.8
40.7 30.3 30.1 30.0 29.8 29.7 29.5 29.4
HO
OH
OH
O
OH
7,2',4'-trihydroxyisoflavanol (THI)
(5)
13C-NMR, Acetone-d6
126 MHz, 298K
f1 (ppm)

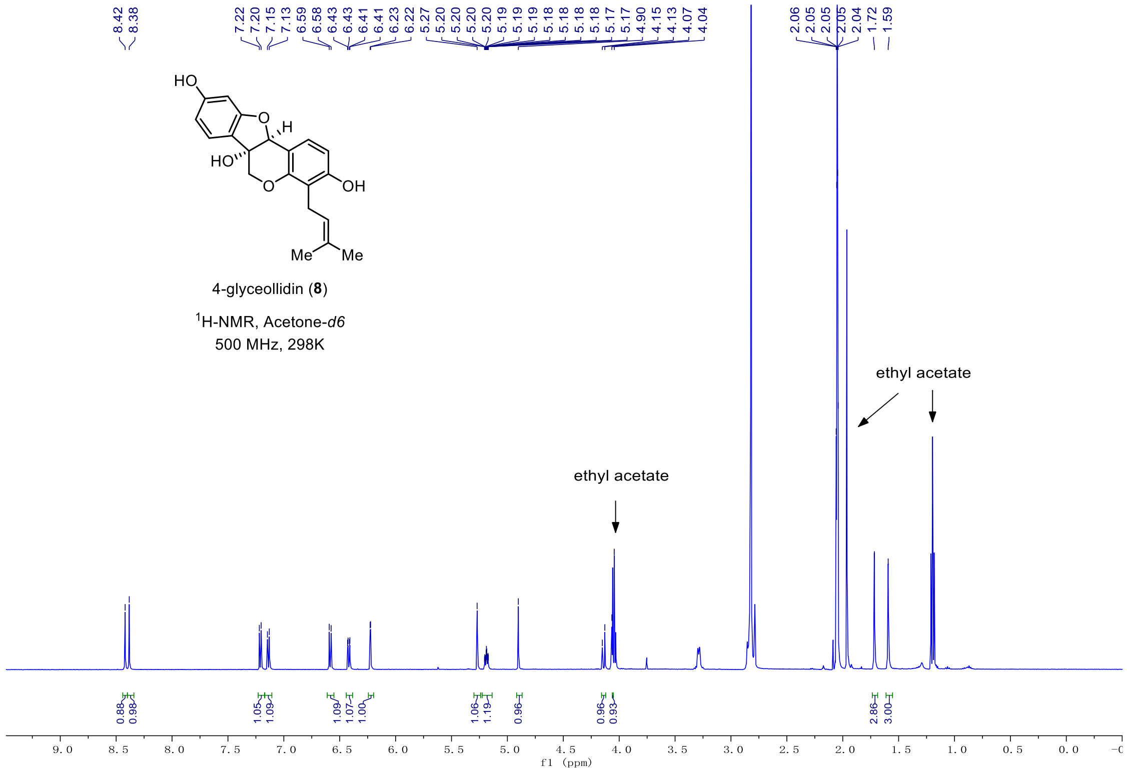
8.42
8.38
7.22
7.20
7.15
7.13
6.59
6.58
6.43
6.43
6.41
6.41
6.23
6.22
5.27
5.20
5.20
5.20
5.20
5.19
5.19
5.19
5.18
5.18
5.18
5.18
5.17
5.17
4.90
4.15
4.13
4.07
4.04
2.06
2.05
2.05
2.05
2.04
1.72
1.59
HO
O
H
HO
O
OH
Me
Me
4-glyceollidin (8)
1H-NMR, Acetone-d6
500 MHz, 298K
ethyl acetate
ethyl acetate
0.88
0.98
1.05
1.09
1.09
1.07
1.00
1.06
1.19
0.96
0.96
0.93
2.86
3.00
f1 (ppm)

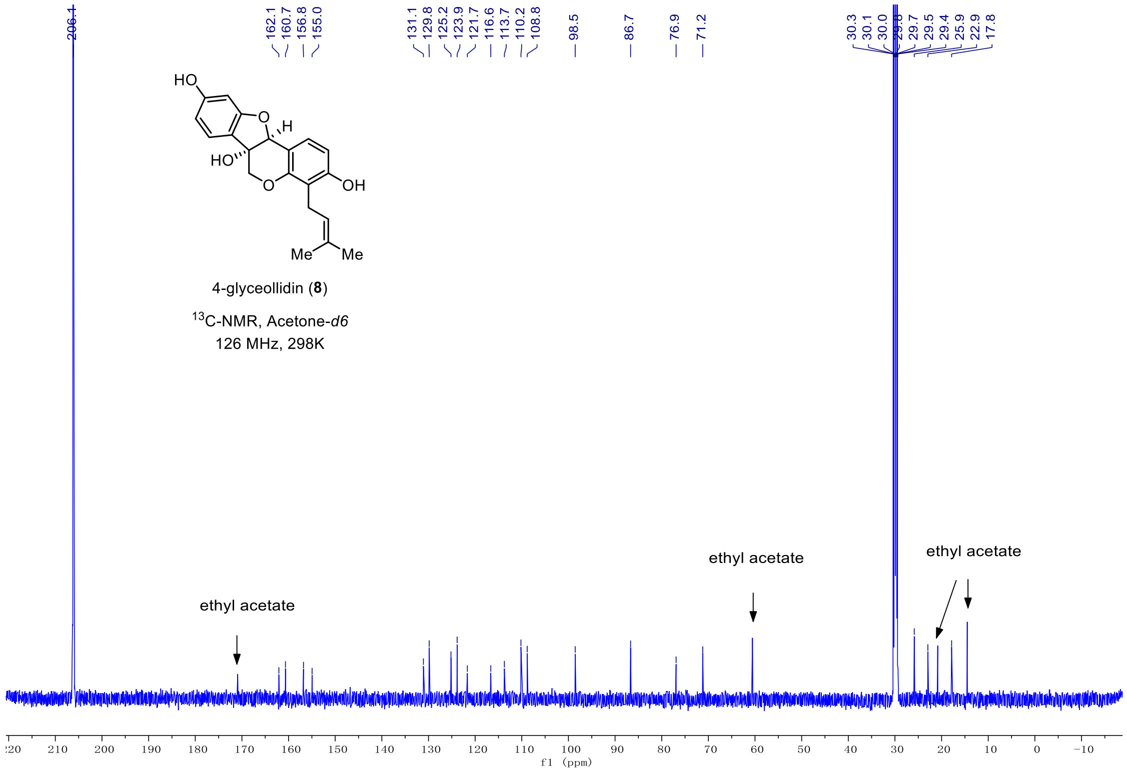
206.1
162.1
160.7
156.8
155.0
131.1
129.8
125.2
123.9
121.7
116.6
113.7
110.2
108.8
98.5
86.7
76.9
71.2
30.3
30.1
30.0
29.8
29.7
29.5
29.4
25.9
22.9
17.8
HO
O
H
HO
O
OH
Me
Me
4-glyceollidin (8)
13C-NMR, Acetone-d6
126 MHz, 298K
ethyl acetate
ethyl acetate
ethyl acetate
f1 (ppm)

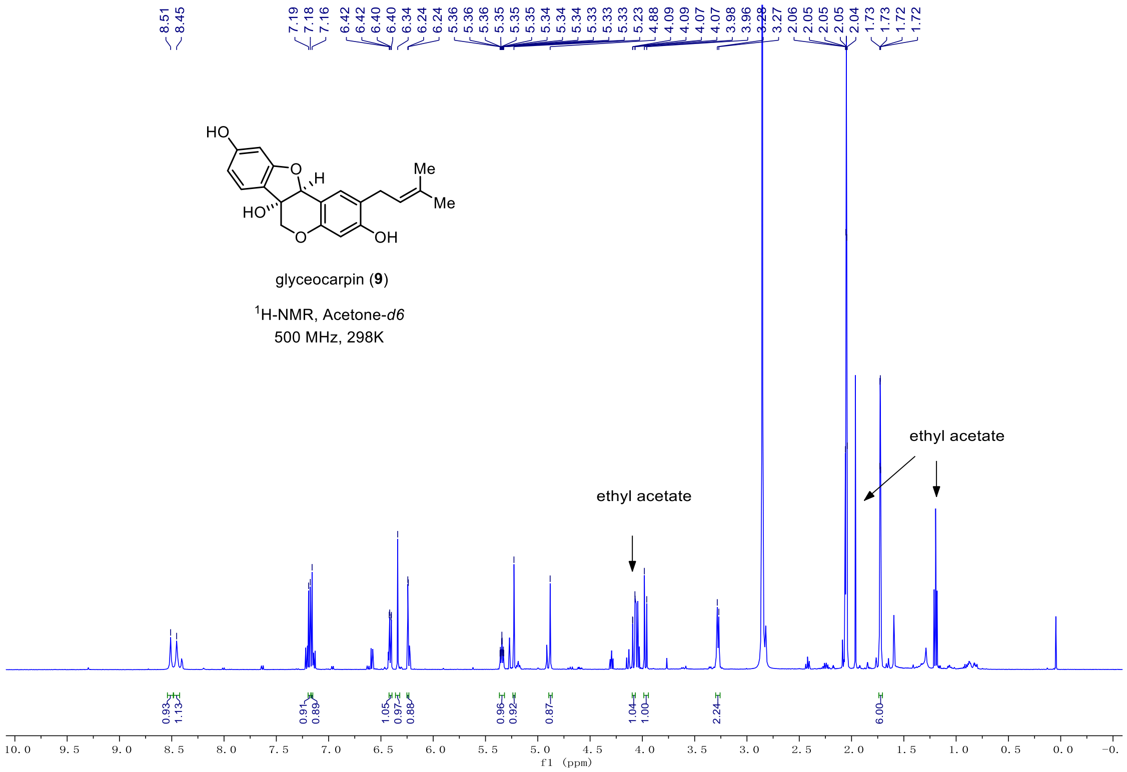
8.51
8.45
7.19
7.18
7.16
6.42
6.42
6.40
6.40
6.34
6.24
6.24
5.36
5.36
5.35
5.35
5.34
5.34
5.34
5.33
5.33
5.33
5.23
4.88
4.09
4.09
4.07
4.07
3.98
3.96
3.28
3.27
2.06
2.05
2.05
2.05
2.04
1.73
1.73
1.72
1.72
HO
O
H
Me
Me
HO
O
OH
glyceocarpin (9)
$^{1}$H-NMR, Acetone-*d6*
500 MHz, 298K
ethyl acetate
ethyl acetate
0.93
1.13
0.91
0.89
1.05
0.97
0.88
0.96
0.92
0.87
1.04
1.00
2.24
6.00
10.0
9.5
9.0
8.5
8.0
7.5
7.0
6.5
6.0
5.5
5.0
4.5
4.0
3.5
3.0
2.5
2.0
1.5
1.0
0.5
0.0
f1 (ppm)

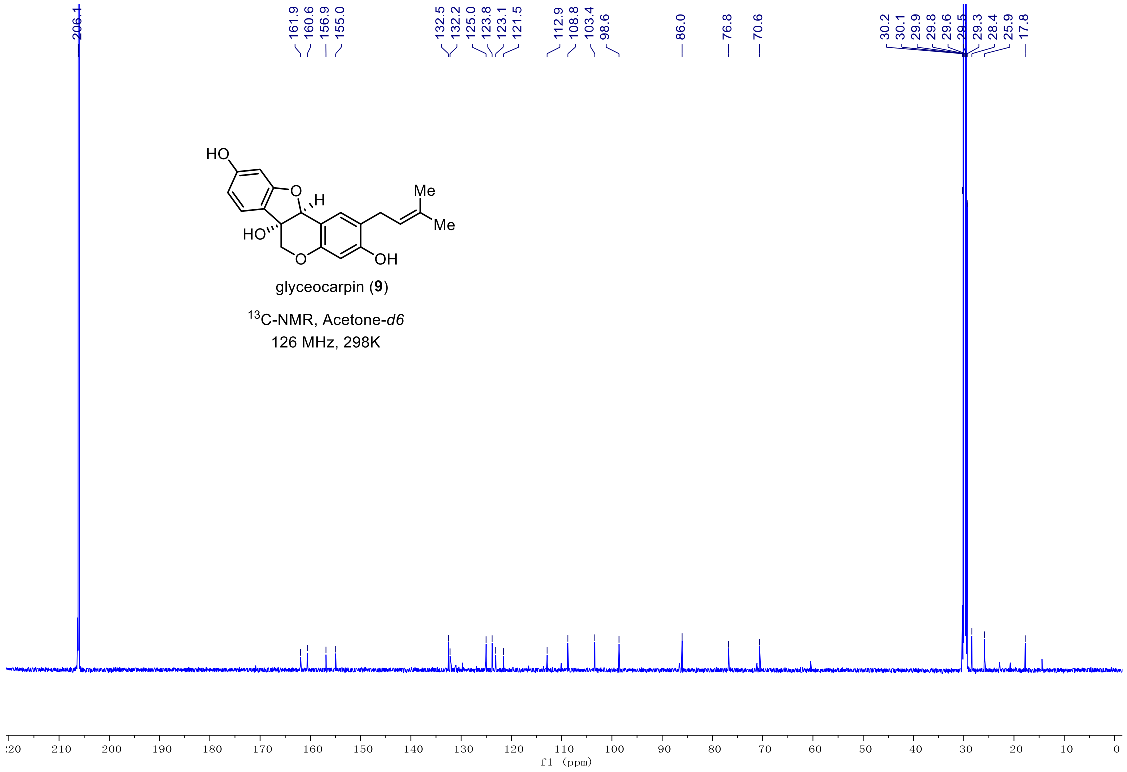
206.1
161.9
160.6
156.9
155.0
132.5
132.2
125.0
123.8
123.1
121.5
112.9
108.8
103.4
98.6
86.0
76.8
70.6
30.2
30.1
29.9
29.8
29.6
29.5
29.3
28.4
25.9
17.8
HO
O
H
Me
Me
HO
O
OH
glyceocarpin (9)
$^{13}$C-NMR, Acetone-*d6*
126 MHz, 298K
210
200
190
180
170
160
150
140
130
120
110
100
90
80
70
60
50
40
30
20
10
0
f1 (ppm)
